# A new tripartite landmark in posterior cingulate cortex

**DOI:** 10.1101/2021.10.30.466521

**Authors:** Ethan H. Willbrand, Benjamin J. Parker, Willa I. Voorhies, Jacob A. Miller, Ilwoo Lyu, Tyler Hallock, Lyndsey Aponik-Gremillion, Alzheimer’s Disease Neuroimaging Initiative, Silvia A. Bunge, Brett L. Foster, Kevin S. Weiner

**Author notes:** co-first author (Ethan H. Willbrand, Benjamin J. Parker). **Corresponding author:** Kevin S. Weiner. Data used in preparation of this article were obtained from the Alzheimer’s Disease Neuroimaging Initiative (ADNI) database (adni.loni.usc.edu). As such, the investigators within the ADNI contributed to the design and implementation of ADNI and/or provided data but did not participate in analysis or writing of this report. A complete listing of ADNI investigators can be found at: http://adni.loni.usc.edu/wp-content/uploads/how_to_apply/ADNI_Acknowledgement_List.pdf.

## Abstract

Understanding brain structure-function relationships, and their development and evolution, is central to neuroscience research. Here, we show that morphological differences in posterior cingulate cortex (PCC), a hub of functional brain networks, predict individual differences in macroanatomical, microstructural, and functional features of PCC. Manually labeling 4,319 sulci in 552 hemispheres, we discovered a consistently localized shallow cortical indentation (termed the inframarginal sulcus; *ifrms*) within PCC that is absent from neuroanatomical atlases, yet co-localized with a region within the cognitive control, but not default mode, network. Morphological analyses in humans and chimpanzees showed that unique properties of the *ifrms* differ across the lifespan and between hominoid species. Intriguingly, the consistency of the *ifrms* also debunks the uniqueness of the morphology of Einstein’s PCC. These findings support a classic theory that shallow, tertiary sulci serve as landmarks in association cortices. They also beg the question: how many other cortical indentations have we missed?

## INTRODUCTION

Elucidating the relationship between morphological and functional features of the cerebral cortex is a central endeavor in neuroscience research. Doing so is particularly important for establishing how individual differences in brain structure-function correspondences develop across the lifespan and evolve across species^1–4^. With these goals in mind, growing evidence shows that individual differences in the morphology of shallow indentations in the cerebral cortex, known as tertiary sulci, co-occur with individual differences in the functional organization of association cortices, as well as individual differences in cognition with translational applications^5–12^. While these findings build on a classic theory^13,14^ suggesting that tertiary sulci are behaviorally meaningful landmarks supporting the functional layout of cognitive representations in association cortices, tertiary sulci have yet to be explored in key association cortices such as posteromedial cortex (PMC).

PMC is routinely considered a central hub of the default mode network^15–18^ and implicated in a broad array of cognitive functions, such as episodic memory, self-referential processing, spatial navigation, and cognitive control^15,17,19–24^. PMC also possesses unique anatomical^19,25^, metabolic^15^, functional^26^, developmental, and evolutionary properties^20,27,28^. Despite this progress in identifying different functions and properties of PMC, the functional neuroanatomy of human PMC remains poorly understood. Precise understanding of its functional-anatomic subdivisions has been impeded by a lack of consensus regarding its basic anatomy. For example, many different anatomical labels and demarcations are employed to refer to the same macroanatomical subdivisions within PMC (**Fig. 1**). Further, previous studies often did not consider sulcal patterning within individual participants^16–18,29^, especially the patterning of shallow tertiary sulci (Methods), which are often overlooked and excluded in neuroimaging software packages^7^.

**Figure 1.**
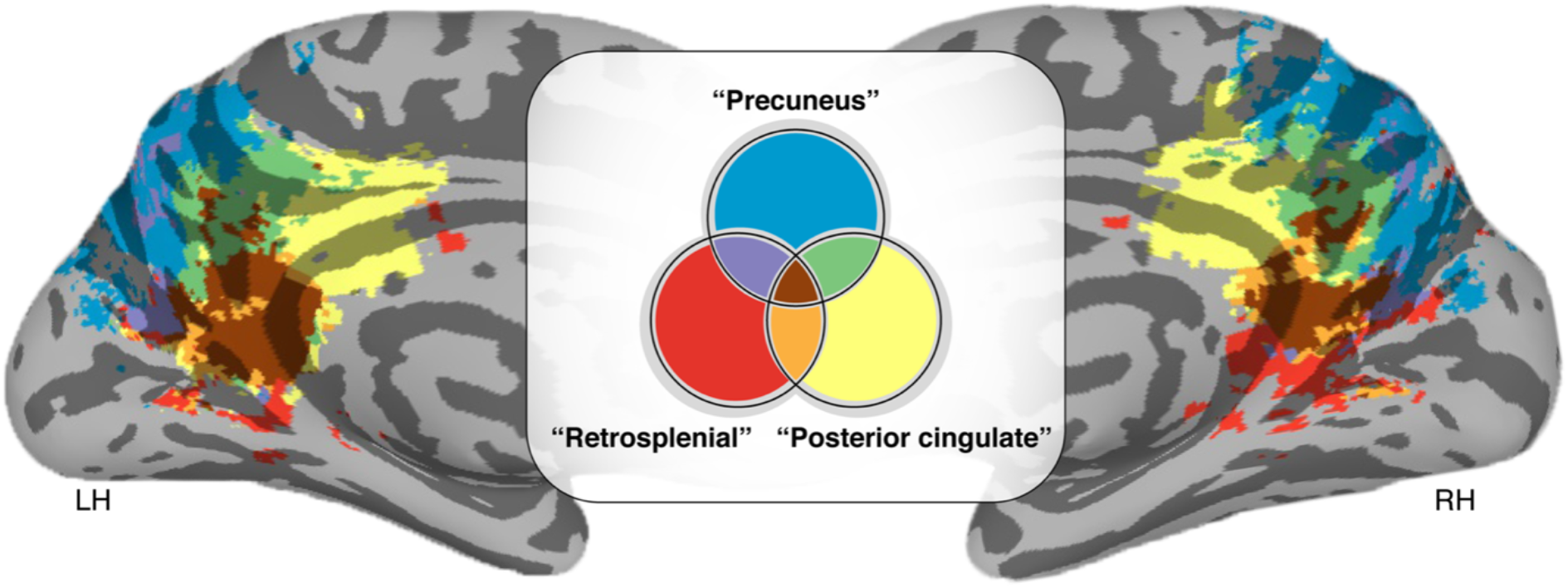
Different names for the same cortical expanse within PMC. MNI152 inflated cortical surface showing *Neurosynth* (https://neurosynth.org) association maps based on the following neuroanatomical search terms: “Posterior cingulate”, “Precuneus,” and “Retrosplenial” in the left (LH) and right (RH) hemispheres. Colors indicate the location, as well as areas of overlap, across the reference studies (947, 1014, and 131 studies, respectively) resulting from each search term. As depicted here, different anatomical labels are used to refer to the same macroanatomical subdivisions of PMC in which 66.3% of the voxels in PMC across these studies have multiple names.

In this work, we implemented a multi-method approach to explore the functional and microstructural relevance of shallow, putative tertiary sulci in PMC. To do so, we manually defined sulci at the individual-participant level in discovery (N=36) and replication (N=36) young adult samples using the most recent definitions of PMC sulci (Methods^30^). Through this process, we discovered a new putative tertiary sulcus (which we refer to as the inframarginal sulcus, *ifrms*) within posterior cingulate cortex (PCC), a subregion of PMC, that is absent from neuroanatomical atlases (Supplementary Materials). In light of this discovery, our subsequent analyses focused on quantifying the structure and function of the *ifrms*. Using structural magnetic resonance imaging (MRI) and cortical surface reconstructions, we first characterized the incidence rates and anatomical features (sulcal depth, cortical thickness, myelination) of the *ifrms* in young adults, relative to neighboring sulci. These analyses indicate that the *ifrms* is a landmark identifying a cortically thick and lightly myelinated locus within PCC. Second, we quantified the relationship between data-driven definitions of functional regions and PMC sulcal definitions in young adults showing that the *ifrms* predicts the location of PCC regions within the cognitive control network^31^. Third, we quantified developmental and evolutionary differences in the morphology of the *ifrms* by performing cross-sectional comparisons across three age groups (juveniles, young adults, and elderly adults) for two hominoid species (humans and chimpanzees). After manually defining over 4,000 sulci, these analyses showed that, while the *ifrms* is identifiable in all human hemispheres examined (even Einstein’s brain), it is only identifiable in a subset of chimpanzee hemispheres. Moreover, we observed differences in cortical thickness and depth between age groups and species. Fourth, we assessed the accuracy with which the *ifrms* could be automatically defined, using novel deep learning algorithms. These algorithms showed that the *ifrms* is more predictable than other PMC sulci that are more prominent in terms of depth and surface area. As we share these algorithms with the field, these tools should help expedite and guide the labeling of PMC sulci in future studies in neurotypical individuals and different patient populations. Together, we identify a new landmark within PCC, providing important progress in elucidating this unique region’s functional organization and further supporting the significance of tertiary sulci in functional brain organization.

## RESULTS

### The inframarginal sulcus: A new shallow indentation in PCC

We first manually defined both deep and shallow sulcal indentations in the posterior cingulate (PCC) and precuneal (PRC) cortices within PMC in discovery (N=36) and replication (N=36) samples of young adults (22-36 years old) from the Human Connectome Project (HCP). 8-11 PMC sulci were identifiable within each hemisphere (**Fig. 2**; **Supplementary Tables 1-2**; **Supplementary Fig. 2.1**; **Supplementary Fig. 3.1-2**). Through this process, we discovered a new putative tertiary sulcus that, through an exhaustive review of historical and modern neuroanatomical atlases (Supplementary Materials), is yet to be named and examined. As it is always located underneath the marginal ramus of the cingulate sulcus (*mcgs*) in every hemisphere, we named this putative tertiary sulcus the inframarginal sulcus (*ifrms*). While our subsequent analyses focused on the structure and function of the *ifrms*, our sulcal definitions in individual participants also identified consistent and variable features of the sulcal patterning within PCC and PRC. In terms of consistency, in PRC, we highlight that while recent studies acknowledge a precuneal sulcus^20,30,32^ (*prcus*), we identified separate posterior (*prcus-p*), intermediate (*prcus-ĩ*), and anterior (*prcus-a*) precuneal sulci in every hemisphere in both datasets (**Supplementary Tables 1-2**; **Supplementary Fig. 2.1; Supplementary Fig. 3.1-2**). In terms of variability, in PCC, we also identified shallow sulci either underneath the splenial sulcus (*spls*), which we refer to as the sub-splenial sulcus (*sspls*), or just anterior to the *ifrms* in a subset of hemispheres (**Supplementary Tables 1-2** for incidence rates). While the latter sulcus was recently identified as the intracingulate sulcus^33^, from our measurements, there are as many as seven shallow sulci along the length of the cingulate gyrus. Thus, we refer to this sulcus as the posterior intracingulate sulcus (*icgs-p*), which, when present, is located underneath the body of the *cgs*, while the *ifrms* is identified more posteriorly underneath the *mcgs*. Unlike the *ifrms*, the incidence rates of *icgs-p* and *sspls* significantly varied by hemisphere (discovery: *X*^2^ = 42.54, df = 2, *p* < 0.001; replication: *X*^2^ = 45, df = 2, *p* < 0.001; **Fig. 2b, 2d; Supplementary Tables 1-2**). As shallowness relative to other sulci is a defining morphological feature of tertiary sulci, we tested if these novel sulci (*sspls, ifrms*, and *icgs-p*) were significantly more shallow than surrounding sulci. Across hemispheres and datasets, the *sspls, ifrms*, and *icgs-p* were shallower than other PMC sulci (*p*-values < 0.001, Tukey’s adjustment; **Fig. 2c, 2e**). Beyond descriptive labeling, we also quantified the intersections between the novel shallow sulci (*ifrms, sspls, icgs-p*) relative to other PMC sulci. Specifically, we correlated the rate of sulcal intersections between hemispheres within and across datasets (Methods; **Supplementary Fig. 2.2; Supplementary Tables 3-6**). This quantitative approach revealed a high positive correlation between hemispheres within each sample (discovery: *r* = 0.74, *p* < 0.001; replication: *r* = 0.9, *p* < 0.001; **Supplementary Fig. 2.2**), as well as between samples (RH: *r* = 0.94, *p* < 0.001; LH: r = 0.82, *p* < 0.001).

**Figure 2.**
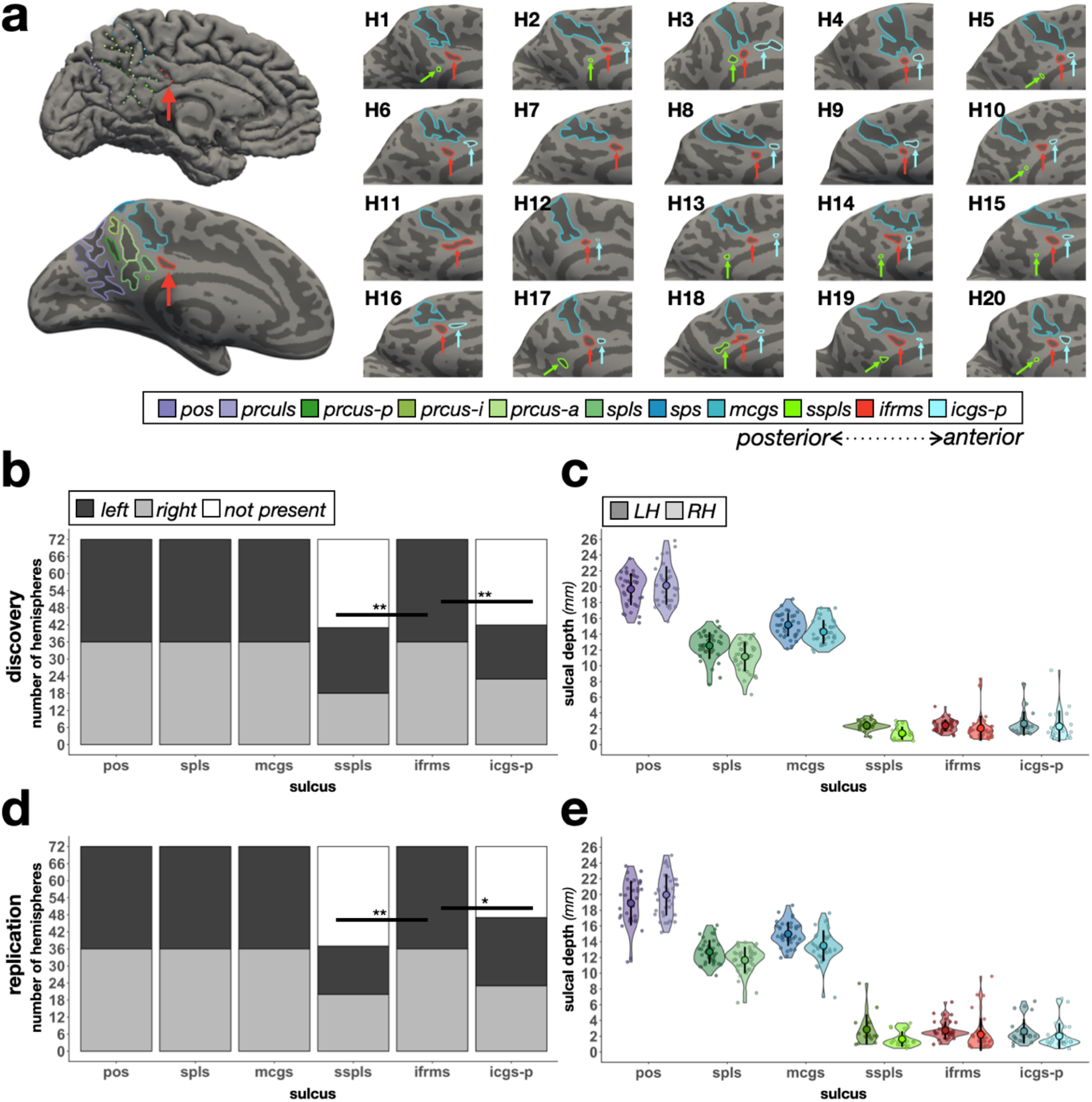
The *ifrms*, but not other shallow sulci in PMC, are identifiable in every hemisphere. **a.** Top left: A cortical surface reconstruction of an individual left hemisphere (LH). Sulci: dark gray; Gyri: light gray. Individual sulci are depicted by dotted colored lines (legend). Bottom left: The same cortical surface but inflated. Right: 20 example hemispheres from the discovery sample. Each hemisphere is indicated by H1, H2, etc. RH images are mirrored so that all images have the same orientation. Each shallow sulcus is designated with an arrow to highlight the consistent location of the *ifrms* underneath the *mcgs*. **b.** Stacked bar plots illustrate the incidence rates of the three shallow sulci (*ifrms, sspls, icgs-p*) relative to three deep sulci (*pos, spls, mcgs*) in the discovery sample (N = 72 hemispheres; dark gray: LH; light gray: RH; white: absent). The *ifrms* is present in every hemisphere, while the *sspls* and *icgs-p* are not (Supplementary Table 1; \**p*<0.05, \*\**p*<0.01). **c.** Sulcal depth (mm) for each individual participant (small colored circles) in the discovery sample. The mean (large colored circles), standard deviation (black line), and kernel density estimate (colored violin) are plotted for each sulcus (LH: darker shades; RH: lighter shades) **d.** The same as **b.**, but for the replication sample (N = 72 hemispheres). As in **b.**, the *ifrms* is present in every hemisphere, while the *sspls* and *icgs-p* are not (Supplementary Table 2). **e.** The same as **c.**, but for the replication sample.

To ensure that the computational processes used to generate the cortical surface reconstruction from individual participant MRI data did not artificially create shallow sulci, we also sought to identify shallow PMC sulci within individual postmortem brains (**Supplementary Fig. 3.3**). In 22 labeled postmortem hemispheres, the *ifrms* was again identifiable in each hemisphere, while the *sspls* and *icgs-p* were not. The *sspls* was present in 90.91% of left and right hemispheres (20/22) and the *icgs-p* was found in 63.64% of left and right hemispheres (14/22). Finally, due to the prevalence of the *ifrms* in all 166 *in-vivo* and postmortem hemispheres, we checked whether this shallow indentation had been identified and labeled in historical publications (**Supplementary Fig. 1**). Our analyses indicated that the *ifrms* was commonly depicted in classic and modern schematics, but to our knowledge, has neither been officially labeled nor empirically explored across age groups and species. In less than a handful of studies, the *ifrms, sspls*, and *icgs-p* were referred to as PCC “dimples”^34^, but otherwise went unreferenced (**Supplementary Fig. 1**).

### The ifrms identifies a cortically thick and lightly myelinated cluster in PCC

Recent work identified a focal cluster within PCC that is cortically thick and lightly myelinated^21,35^, but did not consider covariation with sulcal morphology. As this previously identified cluster was positioned directly under the *mcgs*—in the vicinity of the sulcus we have termed the *ifrms*—we tested the targeted hypothesis that the *ifrms* is cortically thicker and more lightly myelinated than other PMC sulci. To do so, we first extracted cortical thickness and myelination (T_1_w/T_2_w ratio^35^) values from each sulcal label (Methods). We then calculated the ratio between cortical thickness and myelination. Across both samples and hemispheres, the *ifrms* had the greatest thickness/myelination ratio within PCC (**Fig. 3b**). Impressively, when viewing the thickness/myelination map on the cortical surface, the *ifrms* co-localized with this focal anatomical ratio of macroanatomical and microstructural features in PCC (**Fig. 3a**; **Supplementary Fig. 4.1** and **4.2** for the thickness/myelination profiles of all 11 PMC sulci, as well as the individual thickness and myelination values of the four PMC sulci analyzed here).

**Figure 3.**
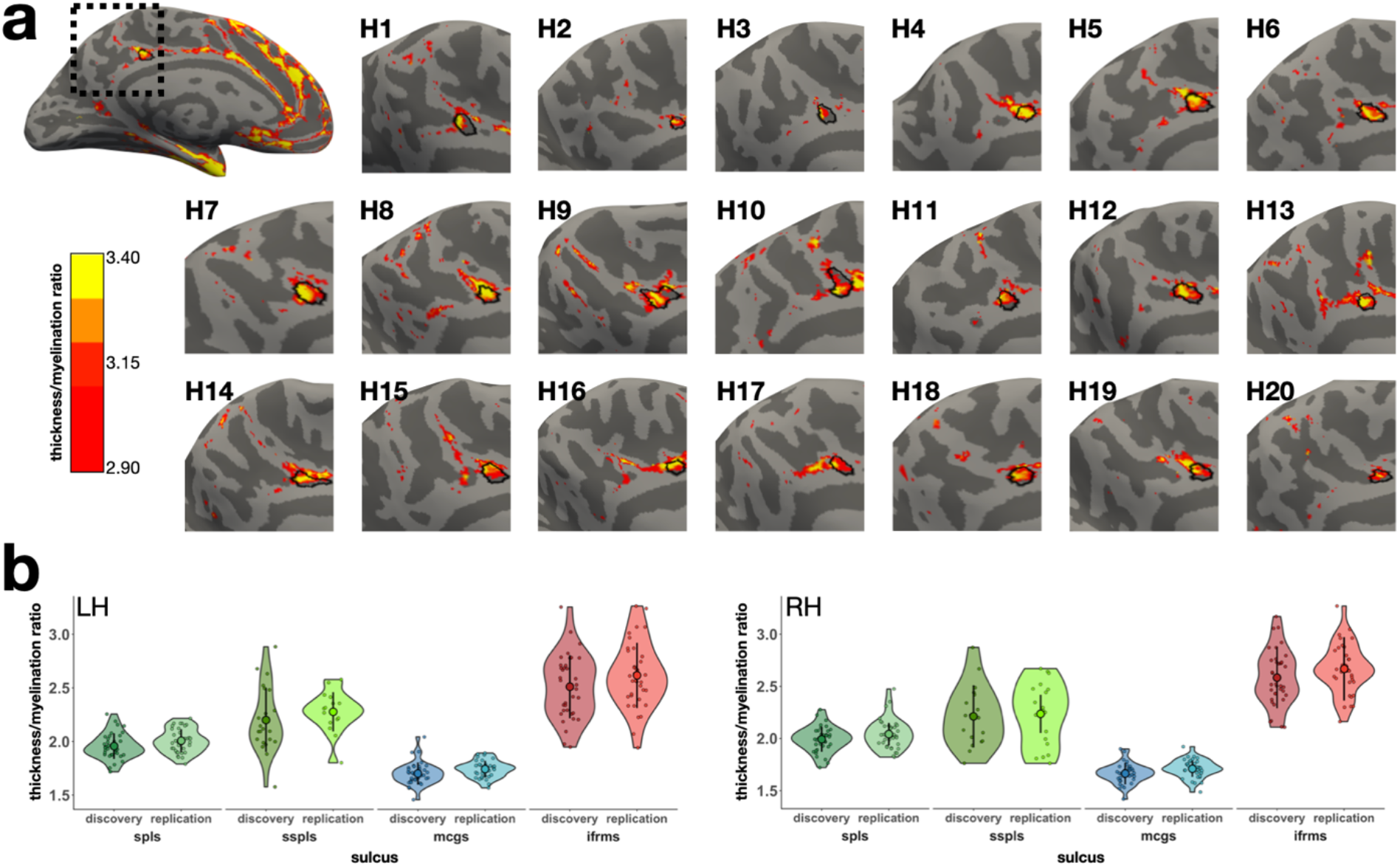
The *ifrms* is a macroanatomical and microstructural landmark in PCC. **a.** A sample left hemisphere, alongside ten left (LH) and ten right (RH) hemispheres (H1, H2, etc.) zoomed in on the PMC, displaying the thickness/myelination ratio (>2.9) relative to the *ifrms* (outlined in black). The right hemispheres have been mirror-reversed to be in alignment with the left hemispheres. The ifrms co-localizes with a portion of PCC that is cortically thick and lightly myelinated. **b.** Thickness/myelination ratio of two deep PMC sulci (*spls* and *mcgs*) and two shallow PMC sulci below them (*sspls* and *ifrms*, respectively) in the discovery and replication samples in the LH and RH. Individual participants from the discovery and replication samples (small colored circles), means (large colored circles), standard deviation (black line), and kernel density estimate (colored violins) are plotted for each sulcus. Each sulcus is colored according to the legend in Figure 2a. The difference in the thickness/myelination ratio is much greater (*p* < 0.001) between the anterior deep and shallow sulci (*mcgs* vs. *ifrms*) compared to the posterior deep and shallow sulci (*spls* vs. *sspls*).

In each participant, the *ifrms* and *sspls* appeared, qualitatively, to be cortically thicker and more lightly myelinated than the deeper sulci positioned above them (*mcgs* and *spls*, respectively). To directly quantify this effect, we conducted 3-way ANOVAs with factors of sulcal type (deep, shallow), PMC position (anterior (*mcgs* and *ifrms*), posterior (*spls* and *sspls*)), and hemisphere (*left, right*) for each sample. In both samples, we observed a main effect of sulcal type (discovery: F(1, 249) = 526.4, *p* < 0.001, η^2^G = 0.68; replication: (F(1, 245) = 682.92, *p* < 0.001, η^2^G = 0.74), in which the shallow sulci (*sspls* and *ifrms*) had a larger thickness/myelination ratio—i.e., were thicker and less myelinated—than the deeper sulci (*spls* and *mcgs*); this was true regardless of hemisphere (**Fig. 3b**). Additionally, there was a sulcal type x position interaction in both samples (discovery: F(1, 249) = 142.14, *p* < 0.001, η^2^G = 0.36; replication: F(1, 245) = 186.6, *p* < 0.001, η^2^G = 0.43), such that the difference in the thickness/myelination ratio was greater in the more anterior PMC, between the *ifrms* and *mcgs*, compared to posterior PMC, between the *sspls* and the *spls*, across hemispheres (*p*-values < 0.001, Tukey’s adjustment in both samples; **Fig. 3b**). While previous work in ventral temporal cortex identified relationships among myelin, curvature, and thickness^36^, this is not necessarily the case across cortex. For example, regarding the *ifrms*, depth only correlated with the thickness/myelination ratio in the right hemisphere of the replication sample (*r* = −0.56, *p* = 0.003, FDR corrected), and cortical thickness did not correlate with myelination in either sample (all *r*-values < 0.31, all *p*-values > 0.12). Taken together, the *ifrms* overlaps with a focal PCC cluster that is cortically thick and lightly myelinated across hemispheres and samples.

### The ifrms predicts the location of functional regions within cognitive control networks

To examine the relationship between the *ifrms* and functional parcellations of PMC, we leveraged resting-state fMRI functional connectivity parcellations from a recently published study^31^ for each individual HCP participant. Importantly, these parcellations were conducted blind not only to cortical folding, but also to our sulcal definitions. **Figure 4a** (left) illustrates the 17-network parcellation on an individual participant’s left hemisphere^31^. To quantitatively determine the relationship between cortical network parcellations and sulcal definitions, we created functional connectivity network profiles (termed “connectivity fingerprints”) by calculating the overlap between each of the 17 networks within the cortical area of a given sulcus on the native hemisphere via the Dice coefficient (Methods; **Fig. 4a**, right) as in our previous work^7^. As it is unclear where the default mode network (DMN) ends and the cognitive control network (CCN) begins from cortical folding alone, we leveraged the functional parcellation in each participant to test if the *ifrms* is a potential landmark identifying the CCN, while the more posterior *spls* is a potential landmark identifying the DMN (connectivity fingerprints for all participants are included in the Supplementary Materials; **Supplementary Fig. 5.1, 5.3**; **Supplementary Fig. 5.4** for connectivity fingerprints of the three precuneal sulci).

**Figure 4.**
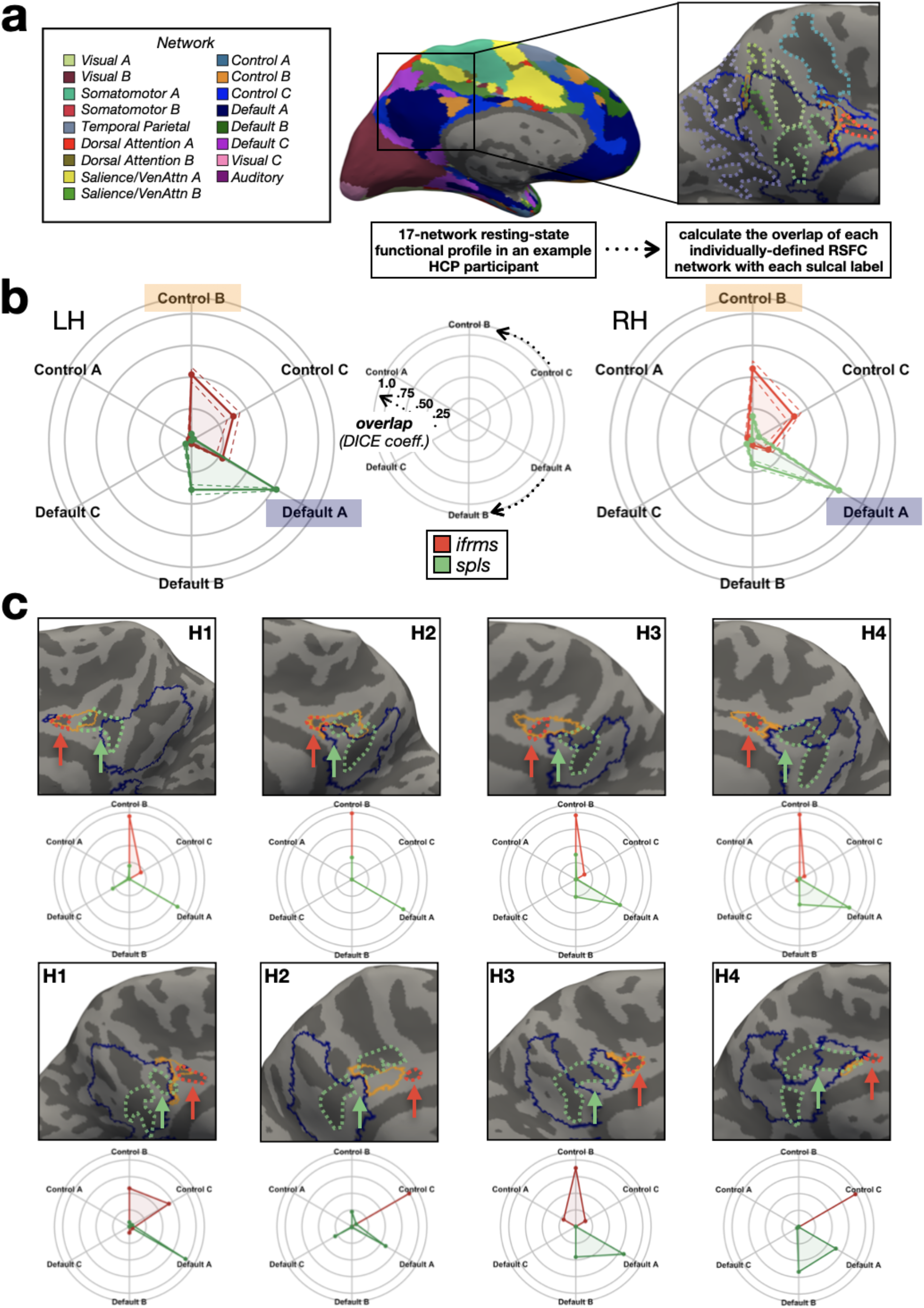
The *ifrms* is a functional landmark in PCC. **a.** Schematic of how resting state functional connectivity (RSFC) profiles were generated for each participant in an example left hemisphere. Individual participant RSFC parcellations were obtained from a recent study^31^, blind to cortical folding, and independent of our sulcal definitions. The connectivity fingerprint represents the overlap of each network within a given sulcal area. **b.** Polar plots showing the mean connectivity fingerprints of the *ifrms* and *spls* in the left hemisphere (LH, left, darker shades) and right hemisphere (RH, right, lighter shades) of the discovery sample. Solid lines: mean. Dashed lines: ±1 sem. Center: Legend for interpreting the polar plots in the left and right images. Arrows denote the direction of each network’s overlap (cognitive control: top; default mode: bottom). The closer to the periphery of the circle, the higher the Dice coefficient. **c.** Connectivity fingerprints for both the RH (top) and LH (bottom) from four individual hemispheres relative to cortical surface reconstructions with CCN-b (orange) and DMN-a (blue) outlines. *ifrms* (red) and *spls* (green) outlines are also included as dotted lines and designated with arrows. The *ifrms* overlaps primarily with CCN-b and CCN-c regions, while the *spls* overlaps more with DMN-a and DMN-b regions with between-hemisphere differences in the different samples (see Supplementary Figure 5.1 for all participants; Supplementary Materials).

Consistent with our hypothesis, the *ifrms* predicted the location of CCN regions, while the *spls* predicted the location of DMN regions in both discovery and replication samples. In both samples, a 3-way repeated measures ANOVA with factors sulcus (*spls, ifrms*), network (*CCN a, b, c* and *DMN a, b, c*), and hemisphere (*left, right*) yielded a sulcus x network interaction (discovery: F(5, 175) = 94.71, *p* < 0.001, η^2^G = 0.41; replication: F(5, 165) = 52.75, *p* < 0.001, η^2^G = 0.31) and a sulcus x network x hemisphere interaction (discovery: F(5, 175) = 3.27, *p* = 0.007, η^2^G = 0.02; replication: F(5, 165) = 8.51, *p* < 0.001, η^2^G = 0.04). Post-hoc analyses in both samples showed that the shallow *ifrms* overlapped significantly more with regions CCN-b and CCN-c than with DMN regions (*p*-values < 0.001, Tukey’s adjustment; **Fig. 4b**; **Supplementary Fig. 5.1-3**), while the *spls* overlapped significantly more with DMN-a and DMN-b than CCN regions (*p*-values < 0.001, Tukey’s adjustment; **Fig. 4b**; **Supplementary Fig. 5.1-3**).

To directly quantify the location of the *ifrms* relative to CCN regions, we performed linear regressions in each hemisphere between *ifrms* coordinates and coordinates of CCN subregions (CCN-b, CCN-c; **Fig. 4, 5; Supplementary Fig. 5.1-3**). *Ifrms* coordinates were predictive of coordinates of CCN regions in both discovery and replication samples (**Supplementary Table 7** for CCN-b and **Supplementary Table 8** for CCN-c). Specifically, the more anterior and superior the *ifrms*, the more anterior and superior the CCN-b (**Fig. 5b**) and CCN-c (**Fig. 5c**) regions. This structure-function correspondence is impressive given the relatively small surface area of both the *ifrms* (average surface area ± std = 77.86 ± 35.83 mm^2^) and the CCN-b (average surface area ± std = 147.53 ± 125.23 mm^2^) compared to the larger CCN-c (average surface area ± std = 390.34 ± 124.94 mm^2^).

**Figure 5.**
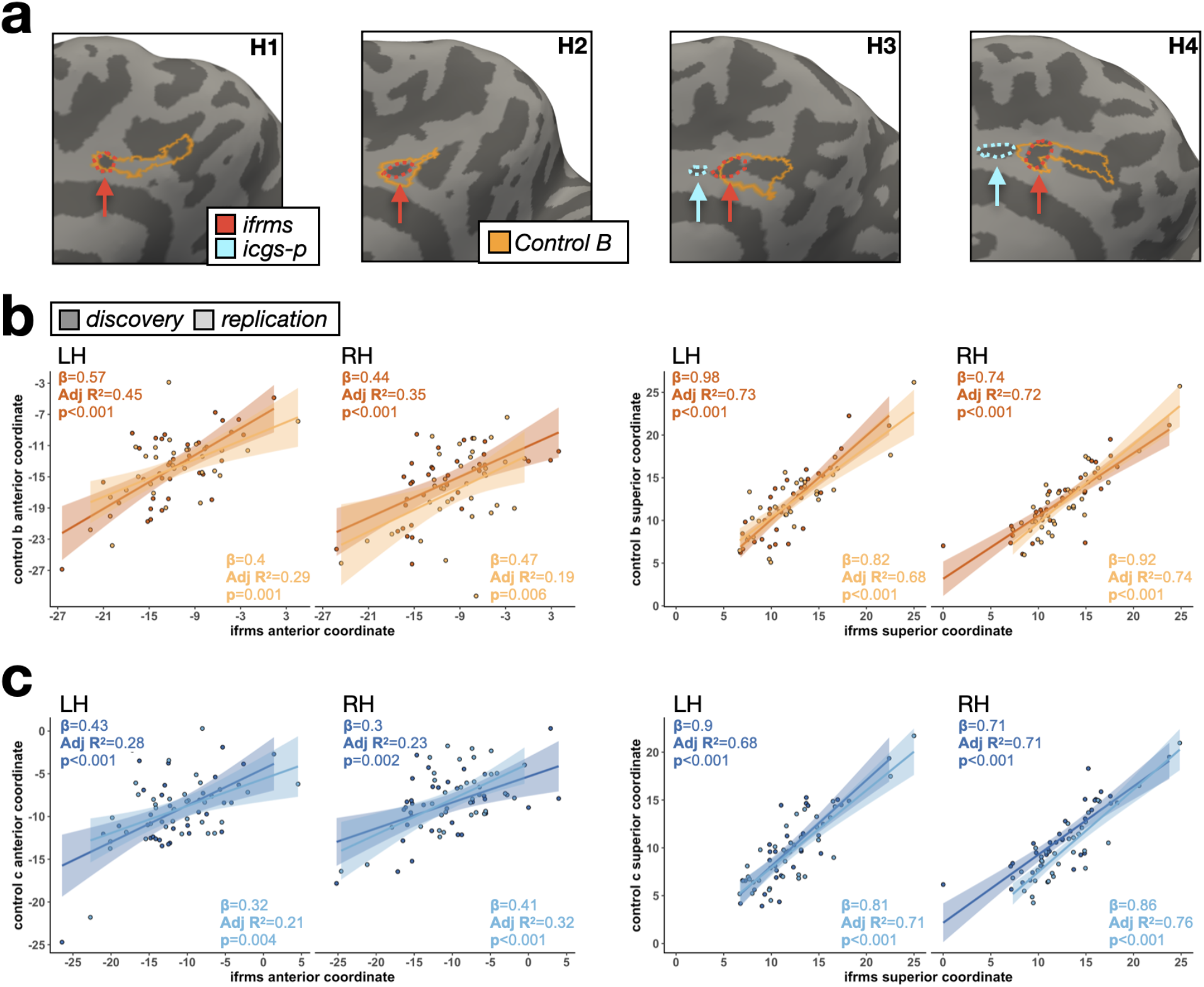
Individual differences in the location of cognitive control regions correlates with variability in the sulcal anatomy. **a.** Four example individual right hemispheres from the discovery sample (two without and two with the *icgs-p*) illustrating the qualitative relationship between the *ifrms* and functionally defined CCN-b (outlined in orange) by Kong and colleagues^31^. The *ifrms* (red) and *icgs-p* (cyan) are outlined and indicated with an arrow. **b.** Linear models (lm) capturing the relationship between the *ifrms* and CCN-b in individual participants (colored circles), from the discovery (dark orange) and replication (light orange) samples, for their mean anterior (left) and superior (right) coordinates for each hemisphere. The best fit line from the regression and ±95% confidence interval are color-matched to each sample. Results for each lm are in the top left corner (slope, adjusted R^2^, and p-value), as well as in Supplementary Table 7. **c.** Same as **b.**, but between the *ifrms* and cognitive control network C (CCN-c; Supplementary Table 8) for both the discovery (dark blue) and replication (light blue) samples. The anterior and superior mean coordinates of the *ifrms* and CCN-b and CCN-c are strongly correlated with one another indicating that variability in the location of the functional CCN-b and CCN-c regions across individuals also correlates with variability in the sulcal anatomy.

### The ifrms is present in human PCC across the lifespan, but is variably present in chimpanzees across age groups

Given the novel discovery of the *ifrms*, we further tested if it is identifiable across the lifespan, as well as in non-human hominoids—and, if so, to compare sulcal depth and cortical thickness of the *ifrms* across age groups and species. To do so, we combined the two young adult samples from the previous section into one group (N = 72; age: range = 22-36, average ± std = 29.06 ± 3.59). We then defined the 8-11 PMC sulci in a juvenile human dataset (N = 72; age: range = 6-18, average ± std = 11.89 ± 3.53) and a healthy older human dataset (N = 72; age: range = 64-90, average ± std = 74.49 ± 5.15), both composed of 72 participants. Lastly, we labeled the *ifrms*— when present—in 60 chimpanzee participants (age: range = 9-51, average ± std = 23.16 ± 9.75), which we also binned into age groups similar to humans: juvenile (age < 22), young adult (22 <= age <= 36), and older adults (age > 36; **Fig. 6a**).

**Figure 6.**
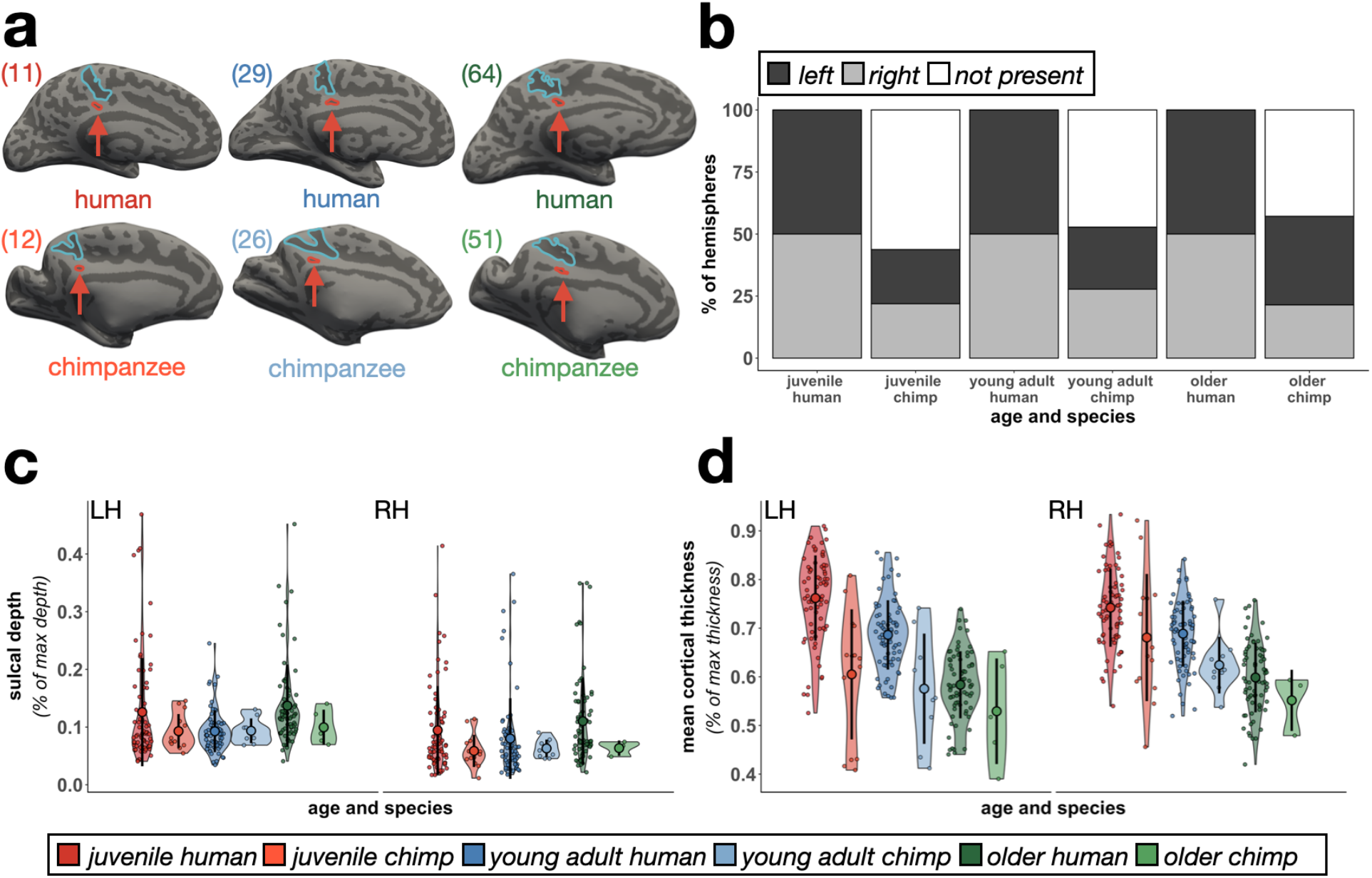
The *ifrms* across species and age groups. **a.** Six example left hemispheres identifying the *ifrms* across age groups (left to right: juvenile, young adult, older adult) and species (top: human; bottom: chimpanzee; cortical surfaces are not to scale). The *ifrms* (red) is outlined and labeled with an arrow below the *mcgs* (blue) in each participant. Ages are in the top left corner of each hemisphere. **b.** The percent of hemispheres with the *ifrms* binned by species and age group. LH: dark gray; RH: light gray; White: absent). The *ifrms* is present in every human, but not chimpanzee, hemisphere measured across age groups. The *ifrms* is only present in about half of chimpanzee hemispheres (juveniles: LH, 43.75% (14/32); RH, 43.75% (14/32); young adults: LH, 50% (9/18); RH, 55.56% (10/18); older adults: LH, 71.43% (5/7); RH, 42.86% (3/7)). **c.** Normalized sulcal depth (% of max depth) of the *ifrms* across age groups and between species plotted for each individual participant in each hemisphere. The mean (large colored circles), standard deviation (black line), and kernel density estimate (colored violin) are also plotted for each sulcus. Each age group and species combination is colored according to the legend. The *ifrms* is deeper in humans compared to chimpanzees and in older adults and juveniles than in young adults across species. **d.** Same layout as **c.**, but for normalized cortical thickness (% of max thickness). The *ifrms* shows an age- and species-related decrease in thickness.

In humans, the *ifrms* was identifiable in every hemisphere examined in juveniles and healthy older adults, as in young adults (**Fig. 6b**; **Supplementary Tables 9 and 10**). Cortical locations (**Supplementary Fig. 6.1** for all 1423 sulcal definitions in juvenile human hemispheres and **Supplementary Fig. 6.2** for all 1386 defined PMC sulci in healthy elderly human hemispheres) were similar to those in the young adult sample and the sulcal intersections (**Supplementary Tables 11-14**) were highly correlated within and between age groups (all *r*s > 0.60, all *p*s < 0.001; **Supplementary Fig. 7.1**). Contrary to the consistent identification of the *ifrms* in humans across age groups, the *ifrms* was identifiable in about half (LH: 50% (30/60); RH: 46.67% (28/60)) of the chimpanzees across age groups (**Fig. 6b**; **Supplementary Fig. 6.3** for all 120 chimpanzee hemispheres).

Morphologically, normalized sulcal depth (Methods) of the *ifrms* is shallower in chimpanzees than humans and differs by age group. Specifically, a 3-way ANOVA with factors of hemisphere (*left, right*), age group (*juvenile, young adult, older*), and species (*human, chimpanzee*) yielded three significant effects. First, there was a main effect of species (F(1, 475) = 6.85, *p* = 0.01, η^2^G = 0.01), wherein the *ifrms* was deeper in humans than in chimpanzees (**Fig. 6c**; **Supplementary Fig. 7.2a** for unnormalized data). Second, there was a main effect of age group (F(2, 475) = 10.68, *p* < 0.001, η^2^G = 0.04), such that the *ifrms* was deeper in the juvenile and older age groups than in the younger adult group (**Fig. 6c**). Third, there was a main effect of hemisphere (F(1, 475) = 15.43, *p* < 0.001, η^2^G = 0.03), wherein the *ifrms* was generally deeper in the left compared to the right hemisphere (**Fig. 6c**).

Additionally, the *ifrms* was relatively cortically thinner in chimpanzees compared to humans across ages groups. A 3-way ANOVA with hemisphere (*left, right*), age group (*juvenile, young adult, older*), and species (*human, chimpanzee*) as factors yielded main effects of age group (F(2, 475) = 135.37, *p* < 0.001, η^2^G = 0.36) and species (F(1, 475) = 63.59, *p* < 0.001, η^2^G = 0.12). There was also a hemisphere x species interaction (F(1, 475) = 6.98, *p* = 0.009, η^2^G = 0.01), such that the *ifrms* in chimpanzees was thicker in the right hemisphere (**Fig. 6d**; **Supplementary Fig. 7.2b** for unnormalized data). Finally, in our historical analyses, we were also able to identify the *ifrms* in a subset of postmortem chimpanzee hemispheres, as well as in other non-human hominoids (gorillas and orangutans). Additionally, we were able to identify a shallow “dimple” in the same location of the brain (which we refer to as the inframarginal dimple, *ifrmd*) in Old World and New World monkeys (Supplementary Materials; **Supplementary Fig. 8**), which is consistent with references to a posterior cingulate dimple in modern research mentioned earlier in these results^34^. Finally, we provide the sulcal depth (mm), surface area (mm^2^), and thickness (mm) values (average ± std) for all 11 PMC sulci across age groups in the Supplementary Materials (**Supplementary Fig. 9** and **Supplementary Tables 15-17**).

### Tools to automatically define the ifrms using convolutional neural networks

As the *ifrms* is a new tripartite landmark in PCC, and due to its prominence across age groups and species, we have created freely available tools to predict the *ifrms* in new datasets. We used recently developed (Methods^37^) spherical convolutional neural networks (CNNs) with context-aware training on the eight PMC sulci identified in the young adult participants. We then quantified the correspondence between the predicted and actual sulcal labels. Here, we report the performance for the *ifrms*; we include the performance for all sulci in the Supplementary Materials.

We found that spherical CNNs were more accurate (t(71) = −3.69, *p* < 0.001, *d* = −0.435) at predicting the *ifrms*’ location than more traditional cortex-based alignment tools (average ± sem; CNN: 0.61 ± 0.03; CBA: 0.5 ± 0.02; **Fig. 7a**). Importantly, the Dice values of the CNN labels were skewed toward higher values, emphasizing that CNNs accurately labeled the *ifrms* a majority of the time (**Fig. 7b**, bottom left). Additionally, while the *ifrms* was, unsurprisingly, not as predictable as the larger, deeper primary sulci (*pos*, *mcgs*, *spls*), the *ifrms* was as predictable as the *prculs*, and even more predictable than the three *prcus* sulci, which were also deeper and larger than the *ifrms* (**Supplementary Fig. 10**). The efficacy of this automated approach speaks to the anatomical consistency of the *ifrms* across individuals. Free dissemination of these tools will expedite the amount of time it takes to define PMC sulci in individual hemispheres in future studies.

**Figure 7.**
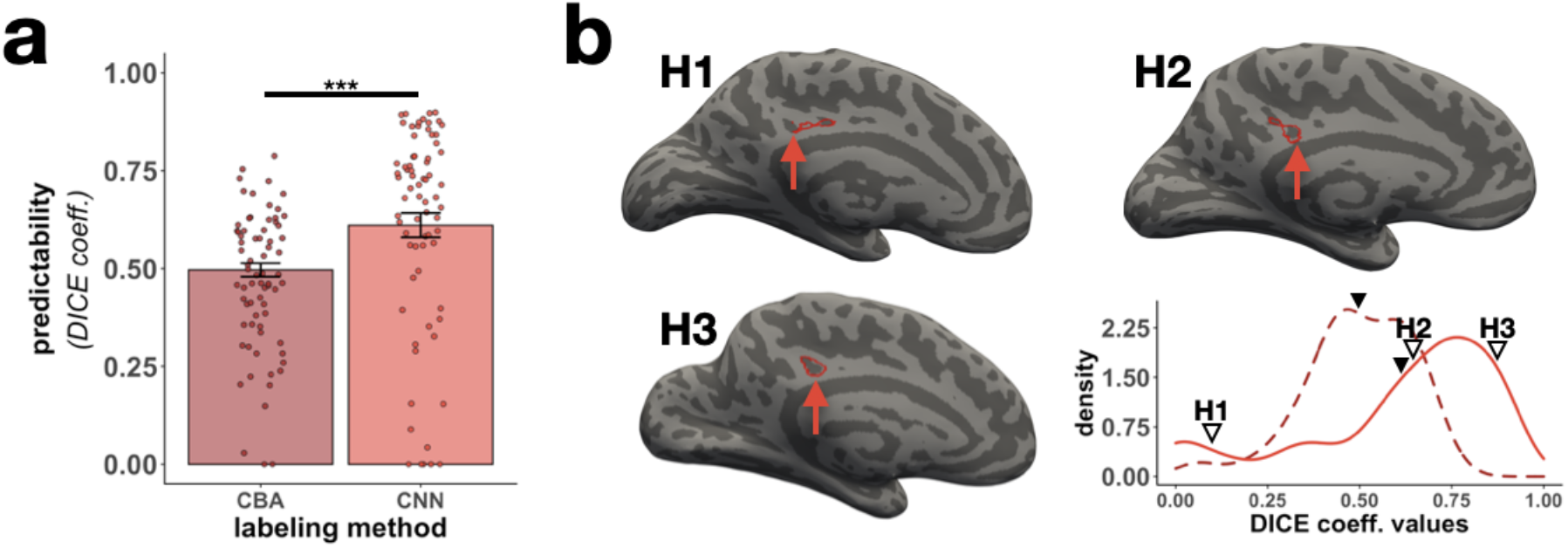
Automatically defining the *ifrms* using deep learning algorithms. **a.** Overlap (Dice coefficient) between predicted and manual location of the *ifrms* for cortex-based alignment (CBA) and spherical convolutional neural network (CNN) methods. Bars represent average values, and the error bars indicate ±1 SEM. Circles represent each individual. Asterisks indicate statistically significant differences between the two approaches (\*\*\**p*<0.001). **b.** Lower right: The density distribution of the *ifrms* Dice coefficient values for the CBA and CNN approaches. The dashed and solid lines represent the CBA and CNN distributions, respectively. The line for each method is colored the same as in **a.** The mean Dice coefficient value is visualized with the solid black triangle for each method (CBA = 0.5, CNN = 0.61). In addition, the Dice coefficient values of three individual participants with the CNN approach (indicated by H1, H2, and H3) are identified with outlined triangles (left to right): low accuracy (H1; Dice = 0.09), mean accuracy (H2; Dice = 0.64), high accuracy (H3; Dice = 0.87). The three participants’ corresponding left hemispheres (indicated by H1, H2, and H3) are provided to visually illustrate the differing degrees of overlap between the automated CNN labels and the manual labels. The automated *ifrms* labels are outlined in red while the manual *ifrms* labels are identified with a red arrow. These data show that the *ifrms* is more accurately defined using the novel CNN approach and importantly, in the majority of hemispheres, the automated labels coincided strongly with the manual labels.

## DISCUSSION

Here, we examined the functional significance of the PMC sulcal organization, focusing on a newly characterized putative tertiary sulcus: the *ifrms*. We report five main findings. First, the *ifrms* is identifiable in every human hemisphere in children, young adults, and elderly adults (even Einstein; **Supplementary Fig. 11**). Second, the *ifrms* is a macroanatomical and microstructural landmark, with the largest thickness/myelination ratio of all PMC sulci. Third, the *ifrms* predicts the location of functional subregions of the cognitive control network (CCN). Fourth, the *ifrms* is present in some, but not all, non-human hominoid hemispheres, as well as appears as a shallow dimple (*ifrmd*) in some non-human primate hemispheres. Fifth, morphological features of the *ifrms* differ across age groups and between species. In the following sections, we discuss these findings in the context of (1) the *ifrms* as a tripartite landmark, (2) tertiary sulci in cortical development and evolution, (3) general limitations and future directions of the present study, and (4) translational applications of tertiary sulci.

We refer to the *ifrms* as a tripartite landmark because it identifies a cortically thick and lightly myelinated portion of PCC, as well as identifies CCN regions. In classic neuroanatomical terms^38^, then, the *ifrms* is an axial sulcus as it co-occurs with a particular cortical area rather than identifies a transition between areas (limiting sulcus). Examining classic and recent microstructural^39,40^ and multimodal parcellations in PCC^35^ shows that the axial definition of the *ifrms* also likely applies to cytoarchitectonic and multimodal areas, which can be explored in future studies (**Fig. 8**)^41^.

**Figure 8.**
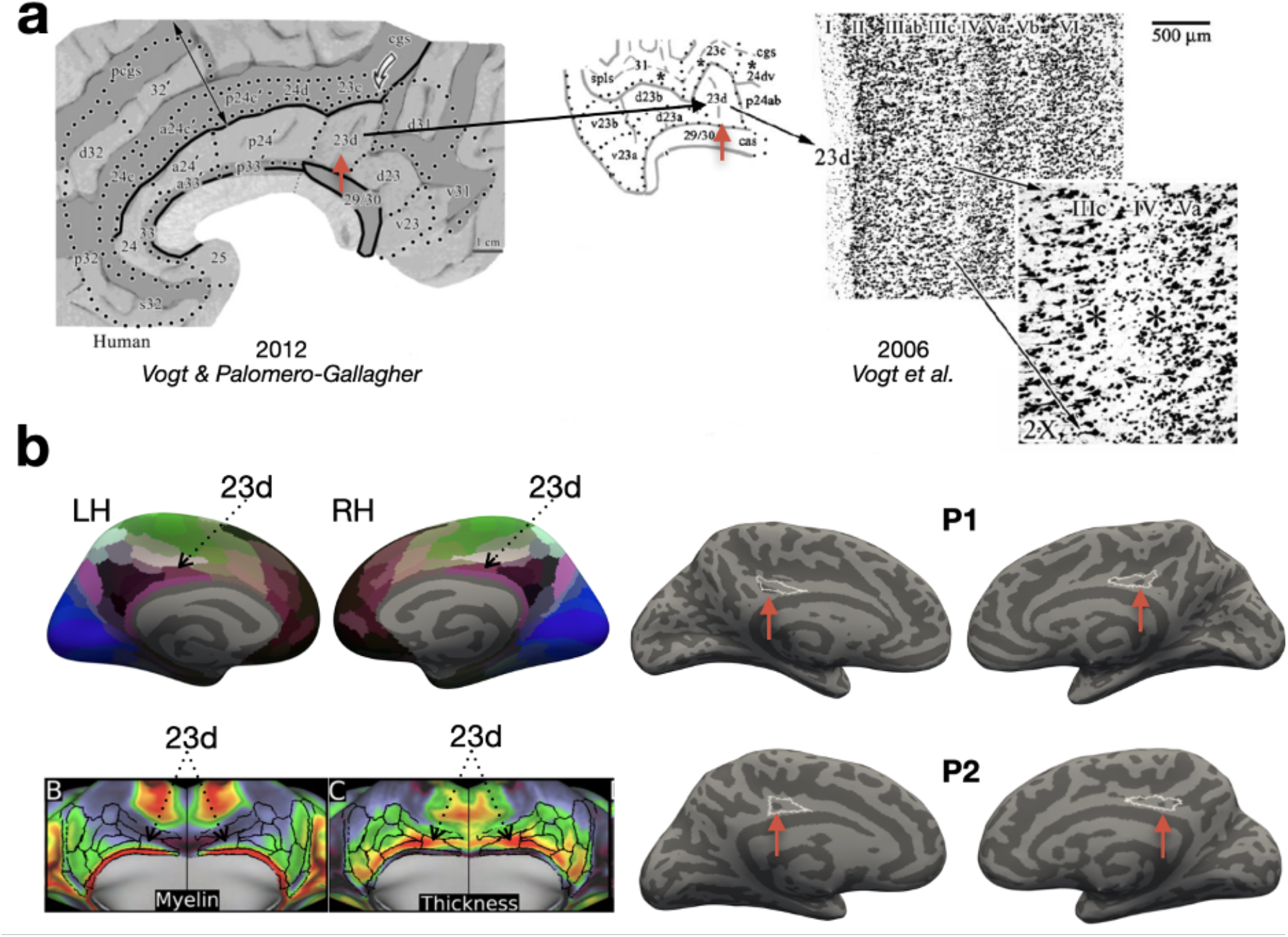
The *ifrms* is a potential cytoarchitectonic and multimodal landmark. **a.** The *ifrms* (red arrow) is located within area 23d, which is dysgranular (as shown by its small layer IV in the inset by Vogt *et al.^39^*). **b.** Left, top: area 23d is defined in the multimodal parcellation by Glasser and colleagues^35^ which is visualized here on the fsaverage cortical surface in magenta (black arrows) in both the left (LH) and right hemisphere (RH). Left, bottom: Like the *ifrms*, area 23d is lightly myelinated (left, green) and cortically thick (right, red), two features which differentiate it from surrounding areas at the group level. Right: The outline of area 23d (in white) defined in the group map projected to two randomly chosen individual participants in both hemispheres. The *ifrms* in individual participants is located within area 23d based on multimodal features at the group level. These observations tentatively identify the *ifrms* as a sulcal landmark based on cytoarchitectonic and multimodal features, which can be tested in individual participants in future studies.

The present findings build on a growing body of work examining the morphological, functional, and cognitive features of tertiary sulci across age groups and species^5,7–12,42,43^. Developmentally, our results extend previous work showing that morphological features of tertiary sulci in other cortical expanses predict performance on different cognitive tasks. For example, the depths of tertiary sulci in lateral prefrontal cortex (LPFC) robustly predict reasoning skills in a developmental cohort^12^. Additionally, the morphology of the paracingulate sulcus (*pcgs*) predicts individual differences in cognition^8–10^, as well as whether individuals with schizophrenia will hallucinate or not^11^. Future studies may identify that morphological features of the *ifrms* may predict behavior and cognition.

Evolutionarily, the present work adds to the growing comparative neuroscience literature classifying the presence/absence of tertiary sulci across species. For example, the mid-fusiform sulcus (*mfs*) was identifiable in every human and non-human hominoid hemisphere examined^43^, while the *pcgs* was much more variable across species^42^. Thus, tertiary sulci are not always identifiable in association cortices, further highlighting the impressive fact that the *ifrms* is identifiable in all hemispheres measured across age groups in humans. Future studies will further assess the prevalence of tertiary sulci among humans and across species in other cortical expanses.

In addition to assessing the incidence of tertiary sulci in a given cortical expanse, future studies could determine whether the presence or absence of a tertiary sulcus directly affects the structure or function of that cortical expanse. For example, the *ifrms* co-localizes with the CCN and may serve as a cytoarchitectonic and multimodal landmark for area 23d in humans. Considering that primate PCC is implicated in cognitive control^20,44,45^ and also possesses area 23d^46^, future research can test if the variably present primate *ifrmd* corresponds to either of these features.

A main limitation of the present work is that the *ifrms* and other tertiary sulci must be manually defined, as they are not included in present approaches that automatically identify most primary and secondary sulci. The main benefit of manual definitions is their precision at the level of individual participants, which allows the most accurate assessment of individual differences in both morphological features themselves and the relationship between morphological features and functional or cognitive significance. However, manual sulcal definitions are slow and arduous, which typically limits sample sizes. For example, while this study includes the manual definition of over 4000 sulci, it only includes 552 hemispheres– a large number for an anatomy study, but a relatively small one for studies exploring the neural basis of individual differences in cognition, functional brain organization, or neurological conditions.

To expedite the labeling process, we benchmarked deep learning algorithms to automatically identify eight of the manually defined PMC sulci, including the *ifrms* (**Fig. 7; Supplementary Fig. 10**). As we share our deep learning methods, these tools can be used in future studies focused not only on tertiary sulci in PMC, but also throughout the cerebral cortex. By expediting sulcal labeling, such methods will facilitate the analysis of larger sample sizes in the service of elucidating the functional significance of variability in tertiary sulci across individuals.

With regards to sulcal classifications, it should be noted that we have classified the novel sulci in this study (*ifrms, sspls*, and *icgs-p*) as putative tertiary sulci based on their morphology (small surface area and shallow depth). However, tertiary sulci are truly classified based on when they emerge during gestation, which is around 36 weeks^1,3,5,6,14,30,47,48^. Therefore, to determine if these sulci are definitively tertiary, future work can seek to identify when exactly they emerge in gestation. Further, to shed light on the relationship between the cross-sectional morphological differences observed here across age groups, future work should quantify the longitudinal interrelations among morphological, microstructural, and functional features of the *ifrms* and other overlooked sulci.

While it is unlikely that all tertiary sulci will serve as landmarks in association cortices, it is important to know which ones do, especially because recent work shows that the morphology of tertiary sulci relates to the symptomatology of several disorders. For example, Garrison *et al*.^11^ showed that the length of the *pcgs* increased the likelihood of hallucinations in individuals with schizophrenia. Additionally, Brun and colleagues^49^ showed that the deepest points within a tertiary sulcus in LPFC were different in individuals with autism spectrum disorder (ASD) compared to neurotypical (NT) individuals. The authors also identified correlations between the depth of these tertiary sulcal points and social communication impairments in ASD individuals^49^. Finally, Ammons and colleagues^50^ recently identified morphological differences in *mfs* morphology between ASD and NT individuals. In addition, *mfs* morphology correlated with the ability of ASD individuals to interpret emotions and mental states from facial features^50^. As such, future studies can build on the findings from our present work to determine whether morphological features of PMC tertiary sulci also have translational applications. For example, there is a considerable amount of research implicating the PCC in Alzheimer’s disease, ASD, attention deficit hyperactivity disorder (ADHD), depression, and schizophrenia (for review see^22^).

In conclusion, through the definition of 4,319 sulci, we not only established a comprehensive description of the sulcal anatomy of human PMC, but also identified a novel sulcus—the *inframarginal* sulcus—that serves as a tripartite landmark in PCC. Methodologically, this study lays the foundation for a myriad of potential PMC and PCC research—whether that be relating the sulci characterized in this study to other anatomical features, additional functions, behavior, or the many disorders that affect this cortical expanse. Evolutionarily and developmentally, morphological analyses between age groups and species show that unique properties of this cortical indentation differ across the lifespan and between species. Theoretically, our findings support Sanides’ classic hypothesis that tertiary sulci may serve as landmarks within association cortices^13,14^. Finally, considering that neuroanatomists have been charting and labeling the outer surface of the human cerebrum for centuries, it is surprising that a sulcus that is observed so consistently across humans was never extensively studied until now. This begs the question: how many other sulci have we yet to uncover?

## METHODS

We describe our methodological approach in three separate sections. The first describes the methods implemented in the main portion of our study examining the anatomical and functional features of sulci in the human posterior cingulate (PCC) and precuneal cortices (PRC) in a young adult cohort, in separate discovery and replication samples. The second focuses on the novel inframarginal sulcus (*ifrms*) and describes the cross-sectional methods implemented to compare the morphological features of the *ifrms* across different age groups (juveniles, young adults, and older adults) and species (humans and chimpanzees). The third details the statistical methods used for the analyses in the previous two sections.

### Examining anatomical and functional features of sulci in the human PCC and PRC in young adults *Participants*

Data for the young adult cohort analyzed in the present study were from the freely available Human Connectome Project database (HCP; https://www.humanconnectome.org/study/hcp-young-adult^51,52^). The discovery dataset consisted of the first five numerically listed HCP participants as well as a random selection of 31 additional participants (17 females, 19 males) whose ages were between 22 and 36 (average ± std = 28.97 ± 3.78). These participants were the same as those used in a previous study examining the anatomical and functional features of sulci within lateral prefrontal cortex^7^. The replication sample consisted of 36 additional participants (19 females, 17 males) randomly selected from the HCP database, with a comparable age range (average ± std = 29.13 ± 3.44, *p* = .84). These data were previously acquired using protocols approved by the Washington University Institutional Review Board.

### Imaging data acquisition

Anatomical T1-weighted (T1-w) MRI scans (0.7 mm voxel resolution) were obtained in native space from the HCP database, along with outputs from the HCP modified FreeSurfer pipeline (v5.3.0)^53–55^. Additional details on image acquisition parameters and image processing can be found in Glasser *et al*.^55^. Maps of the ratio of T1-weighted and T2-weighted scans, which is a measure of tissue contrast enhancement related to myelin content, were downloaded as part of the HCP ‘Structural Extended’ release. All subsequent sulcal labeling and extraction of anatomical metrics were calculated from the cortical surface reconstructions of individual participants generated through the HCP’s custom modified version of the FreeSurfer pipeline^55^.

### Anatomical analyses

#### Macroanatomical boundaries of the PCC and PRC within PMC

Based on previous work, there is extensive variability regarding how PCC and PRC are defined (**Fig. 1**). Here, we defined the PRC by the following posterior, inferior, and anterior boundaries, respectively, in the medial parietal cortex, as shown in **Fig. 1** and **Fig. 2a** (left): 1) the parieto-occipital sulcus (*pos*), 2) splenial sulcus (*spls*), and 3) marginal ramus of the cingulate sulcus (*mcgs*), respectively. It should be noted that while the *spls* nomenclature is used by Vogt *et al*.^34,39^ and our group, due to its typical location superior to the splenium of the corpus callosum, this sulcus is also known as the subparietal sulcus (*sbps*^20,30^). Nevertheless, we adopt the *spls* nomenclature because there is an additional shallow sulcus in PCC underneath the *spls* (expanded on further below), which we refer to as the sub-splenial sulcus (as opposed to sub-subparietal sulcus). We define PCC as the posterior portion of the cingulate gyrus bounded inferiorly by the callosal sulcus (*cas*), superiorly by the cingulate sulcus (*cgs*), *mcgs*, and the *spls*, and posteriorly by the *pos*.

#### PCC and PRC sulci

Based on the most recent and comprehensive atlas of sulcal definitions throughout the cerebral cortex^30^, we consider 11 sulci that are either bounding or are located within PCC and PRC. In addition to the *pos*, *spls*, and *mcgs* as mentioned in the previous section, which bound the PRC, we also defined five sulci within the PRC: the precuneal limiting sulcus (*prculs*), the superior parietal sulcus (*sps*), and three precuneal sulci (*prcus-p*, *prcus-i*, *prcus-a*). The *prculs* branches off of the superior portion of the *pos* within the posterior PRC. The *sps* is located within the medial portion of the superior PRC and often extends into superior portions of lateral parietal cortex. While previous research identifies a single precuneal sulcus^32^ or includes three in their schematic, but only explicitly labels one component as the *prcus*^30^, we reliably identified three different *prcus*—often intersecting in different combinations with the *spls, mcgs, sps*, and each other (**Supplementary Tables 3, 5, 11, 13**). Because we can identify each sulcus in every hemisphere, we propose the following labels for these sulci (mirroring the labeling approach for the three components of the posterior middle frontal sulci): 1) posterior precuneal sulcus (*prcus-p*), 2) intermediate precuneal sulcus (*prcus-i*), and 3) anterior precuneal sulcus (*prcus-a*).

In addition to these eight sulci, we also considered three shallow PCC sulci. Past research has referred to these indentations in a variety of ways, typically referring to them as inconsistent dimples or as branches of the *cas* or *cgs* (**Supplementary Fig. 1**), and most recently identifying an intracingulate sulcus in the middle portion of the cingulate gyrus^33^. We consistently identify a shallow sulcus in every hemisphere. As this sulcus is always inferior to the *mcgs*, we label it the inframarginal sulcus (*ifrms*). We also identify two tertiary sulci anterior and posterior to the *ifrms* in a subset of individuals and hemispheres. Posteriorly, a variable sulcus is sometimes present inferior to the *spls*, which we label the subsplenial sulcus (*sspls*). Anteriorly, a variable sulcus is also sometimes present, which we refer to as the posterior intracingulate sulcus (*icgs-p*). We suggest this label because while a recent study suggests labeling an intracingulate sulcus^33^, our data in individual participants indicate that there are many intracingulate sulci, which necessitates more precise labels discriminating these sulci from one another.

#### Sulcal labeling

Each PCC and PRC sulcus was manually defined within each individual hemisphere on the FreeSurfer *inflated* mesh with tools in *tksurfer* as described in our previous work^7,12^. Specifically, the *curvature* metric in FreeSurfer distinguished the boundaries between sulcal and gyral components, and manual lines were drawn to separate sulcal components, as well as the appearance of sulci across the *inflated, pial*, and *smoothwm* surfaces. The sulcal labels were generated using a two-tiered procedure. The labels were first defined manually by trained raters (E.W., B.P., and T.H.) and then finalized by a neuroanatomist (K.S.W.). In each hemisphere, we first labeled the more stable sulci bounding the PCC and PRC (e.g., *pos*, *spls*, *mcgs*), and then we identified the remaining sulcal components within the PCC and PRC. All anatomical labels for a given hemisphere were fully defined before any morphological or functional analyses of the sulcal labels were performed.

#### Extracting anatomical features from sulcal labels

After all sulci were defined, anatomical features (sulcal depth (mm), cortical thickness (mm), surface area (mm^2^), and myelination (T_1_w/T_2_w ratio)) were extracted. Raw mm values for sulcal depth were calculated from the sulcal fundus to the smoothed outer pial surface using a custom-modified version of a recently developed algorithm building on the FreeSurfer pipeline^57^. Though classic work in post-mortem brains by Ono and colleagues^32^ did not identify all PMC sulci included in the present study, three sulci (*pos*, *spls*, and *mcgs*) overlapped between the two studies and displayed similar ranges for sulcal depth (Supplementary materials). Mean cortical thickness (mm) and surface area (mm^2^) were extracted from each sulcus using the built-in *mris_anatomical_stats* function in FreeSurfer^58^. Average values for myelination were obtained using an *in vivo* proxy for myelination: the T_1_w/T_2_w ratio for each individual hemisphere available in the HCP dataset^59,60^.

#### Quantitative assessment of incidence rates of PCC and PRC tertiary sulci

To compare the incidence rates of tertiary sulci (*ifrms, icgs-p*, and *sspls*) in PCC and PRC, chi-squared tests were implemented, along with follow-up post hoc pairwise comparisons.

#### Qualitative labeling of PCC and PRC sulci in post-mortem hemispheres

To assure that our labeling of PCC and PRC sulci was not an artifact of the cortical surface reconstruction process, we also identified PCC and PRC sulci within *post-mortem* human brains (22 hemispheres total) from a classic neuroanatomy atlas^61^ (**Supplementary Fig. 3.3**). Critically, the *ifrms* was present 100% of the time.

#### Quantitative assessment of PMC sulcal depth

We quantitatively compared PMC sulcal depth using a 2-way ANOVA with sulcus (*pos, spls, mcgs, sspls, ifrms*, and *icgs-p*) and hemisphere (*left, right*) as factors.

#### Quantitatively assessing the macroanatomical and microanatomical properties of PMC sulci along an anterior-posterior dimension

To quantitatively assess how macroanatomical (cortical thickness) and microanatomical (myelination) features of PMC sulci vary along an anterior-posterior dimension, we compared the thickness/myelination ratio of the two deeper sulci (*spls* and *mcgs*) and the two shallow sulci inferior to them (*sspls* and *ifrms*, respectively) with a 3-way ANOVA using sulcal type (primary/secondary and tertiary), anatomical position (anterior (*mcgs* and *ifrms*) and posterior (*spls* and *sspls*)), and hemisphere (*left* and *right*) as factors.

#### Predictive labeling of sulcal location using convolutional neural networks

For full methodological details, please see Lyu *et al*.^37^, which describes the methodological pipeline in full. Briefly, the ability to automatically define PMC sulci was compared using two methods: cortex-based alignment (described in the next paragraph) and deep learning (convolutional neural network, CNN) with context aware training. For the CNN, the algorithms were trained on sulcal labels in a 5-fold cross validation manner (60% of participants for training), and then the trained model was chosen with peak performance on the validation set (20% of participants) for each fold. The overall performance was iteratively calculated on the left-out participant in the test set (20% of participants). This approach was developed previously on sulci in lateral prefrontal cortex by Lyu and colleagues^37^ and applied here to PMC sulci. The implementation of the CNN contains two key modifications compared to other CNNs: during the learning phase, surface data augmentation and context aware training are implemented. The former adds flexibility by implementing intermediate deformations (if needed) to better align sulci across participants and to enhance the model generalizability. The latter incorporates the spatial information of the primary and secondary sulci to guide the automatic labelling of the small and more variable sulci. Prediction performance was determined by calculating the Dice coefficient between the predicted sulcus and the ground truth, manually defined sulcus using the following formula:

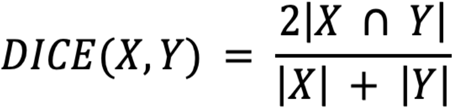

Prediction performances for CBA and deep learning with context aware training were compared using a paired t-test.

#### Predictive labeling of sulcal location using cortex-based alignment

First, all sulcal labels were registered to a common surface template surface (*fsaverage*) using cortex-based alignment^56^. Sulcal probability maps were then calculated to describe the vertices with the highest alignment across participants for a given sulcus. A map was generated for each sulcus by calculating, at each vertex in the *fsaverage* hemisphere, the number of participants with that vertex labeled as a given sulcus, divided by the total number of participants. The map was then made more precise by constraining the original probability maps into *maximum probability maps* (MPMs) by only including vertices where (1) more than 33% of participants were included in the given sulcal label and (2) the sulcus with the highest value of participant overlap was assigned to a given vertex. This step also helped to avoid overlap among sulci. In a leave-one-out cross-validation procedure, probability maps were generated from the combined young adult sample from N = 71 participants. These maps were then registered to (1) the held-out participant’s native cortical surface, separately for each dataset, and (2) the held-out participant’s native cortical surface of the opposing dataset. Prediction performance was again determined by calculating the Dice coefficient between the predicted sulcus and the manually defined sulcus.

### Functional analyses

#### Creating sulcal “connectivity fingerprints” from resting-state network parcellations

To determine if the *ifrms* is functionally distinct from the nearby *spls*, we generated functional connectivity profiles (“connectivity fingerprints”) using a recently developed analysis^7^. This analysis implements a four-pronged approach. First, resting-state network parcellations for each individual participant were used from Kong *et al*.^31^, who generated individual network definitions by applying a hierarchical Bayesian network algorithm to produce maps for each of 17-networks in individual HCP participants^62^. These data were calculated in the template HCP *fs_LR 32k* space. Importantly, this parcellation was conducted blind to both cortical folding and our sulcal definitions. Second, we resampled the network profiles for each participant onto the *fsaverage* cortical surface, and then to each native surface using CBIG tools (https://github.com/ThomasYeoLab/CBIG). Third, we then calculated the overlap between a sulcus with each of the 17 resting-state networks for each participant via the Dice coefficient. Fourth, a 3-way (*sulcus, network*, and *hemisphere*) repeated measures ANOVA was run to determine if the network profiles of the *ifrms* and the more posterior *spls* were differentiable from one another. The functional networks included in the statistical analyses were limited to the default mode (DMN) and cognitive control networks (CCN) since only these networks overlapped prominently with these sulci. The same analyses, but for the three sulcal *prcus* components are included in the Supplementary Materials and visualized in **Supplementary Fig. 5.4**.

#### Determining if the location of the ifrms is predictive of CCN-b and CCN-c

Building off of the functional findings for the *ifrms*, we aimed to quantify if the location of the *ifrms* was predictive of the location of the two CCN subregions that it overlapped with most (CCN-b, CCN-c), especially the smaller CCN-b. To test this, we leveraged the mean RAS values of the *ifrms* and the two cognitive control networks obtained in the previous analyses to conduct linear regressions for each hemisphere between the mean RAS coordinates of the *ifrms* (predictor) and CCN-b or CCN-c (outcome).

### Examining morphological features of the ifrms across the lifespan and between species *Participants*

This part of the study leveraged three human neuroimaging datasets. The first was composed of the combined young adult samples (N = 72 participants) described previously. The second human dataset was composed of juveniles, and the third of healthy older adults. For comparative analyses, one chimpanzee neuroimaging dataset was used.

#### Human (juveniles)

72 participants (30 females, 42 males) were randomly selected from the Neurodevelopment of Reasoning Ability study (NORA) study^63–65^. This sample consists of typically developing individuals between the ages of 6 and 18 (average ± std = 11.89 ± 3.53). All participants were screened for neurological impairments, psychiatric illness, history of learning disability, and developmental delay. Additionally, all participants and their parents gave their informed assent or consent to participate in the study, which was approved by the Committee for the Protection of Human participants at the University of California, Berkeley.

#### Human (healthy older adults)

72 healthy older adult participants (37 females, 35 males) with ages ranging from 64 to 90 years old (average ± std = 74.49 ± 5.15) were randomly selected from the Alzheimer’s Disease Neuroimaging Initiative (ADNI) database (adni.loni.usc.edu).

#### Chimpanzee

60 (37 female, 23 male) chimpanzee (*Pan Troglodytes*) anatomical T1 scans were chosen from the National Chimpanzee Brain Resource (www.chimpanzeebrain.org; supported by NIH grant NS092988). Chimpanzees were between the ages of 9 and 51 (average ± std = 23.16 ± 9.75). The chimpanzees were members of the colony housed at the Yerkes National Primate Research Center (YNPRC) of Emory University. All methods were carried out in accordance with YNPRC and Emory University’s Institutional Animal Care and Use Committee (IACUC) guidelines. Institutional approval was obtained prior to the onset of data collection. Further data collection details are described in Keller *et al*.^66^. These are the same 60 chimpanzee cortical surfaces examined in Miller *et al*.^43^.

### Imaging data Acquisition

#### Human (juveniles)

Two high-resolution T1-weighted MPRAGE anatomical scans (TR = 2300 ms, TE = 2.98 ms, 1 × 1 × 1 mm^3^ voxels) were acquired using the Siemens 3T Trio fMRI scanner at the University of California Berkeley Brain Imaging Center.

#### Human (healthy older adults)

T1-weighted MPRAGE anatomical scans were obtained for cortical morphometric analyses in these participants from the ADNI online repository (http://adni.loni.usc.edu). The exact scanning parameters varied across the sample (see **Supplementary Table 18** for a breakdown of the different scanning parameters used).

#### Chimpanzee

Here we briefly describe the scanning parameters that are described in more thorough detail in Keller *et al*.^66^. The T1-weighted magnetization-prepared rapid-acquisition gradient echo (MPRAGE) MR images were obtained using Siemens 3T Trio MR system (TR = 2300 ms, TE = 4.4 ms, TI = 1100 ms, flip angle = 8, FOV = 200 mm x 200 mm) at YNPRC in Atlanta, Georgia. Prior to reconstructing the cortical surface, each chimpanzee T1 was scaled to the size of the human brain. As described in Hopkins *et al*.^67^, within FSL, (1) the BET function was used to automatically strip away the skull, (2) the FAST function was used to correct for intensity variations due to magnetic susceptibility artifacts and radio frequency field inhomogeneities (i.e., bias field correction), and (3) the FLIRT function was used to normalize the isolated brain to the MNI152 template brain using a seven degree of freedom transformation (i.e., three translations, three rotations, and one uniform scaling), which preserved the shape of individual brains. Afterward, each T1 was segmented using FreeSurfer. The fact that the brains are already isolated, along with bias-field correction and size-normalization, greatly assisted in segmenting the chimpanzee brain in FreeSurfer. Furthermore, the initial use of FSL also has the specific benefit of enabling the individual brains to be spatially normalized with preserved brain shape. Lastly, the values of this transformation matrix and the scaling factor were saved for later use.

### Cortical surface reconstruction

Each T1-weighted image was visually inspected for scanner artifacts. Afterwards, reconstructions of the cortical surfaces were generated for each participant from their T1 scans using a standard FreeSurfer pipeline (FreeSurfer (v6.0.0): surfer.nmr.mgh.harvard.edu/^53,54,56^). Cortical surface reconstructions were created from the resulting boundary made from segmenting the gray and white matter in each anatomical volume with FreeSurfer’s automated segmentation tools^53^. Each reconstruction was inspected for segmentation errors, which were then manually corrected when necessary. As in young adults, all subsequent sulcal labeling and extraction of anatomical metrics were calculated from cortical surface reconstructions from individual participants.

### Developmental and comparative analysis of the ifrms

#### Sulcal labeling

The same 8-11 PCC and PRC sulci defined in young adults were manually identified in individual juvenile and healthy older human hemispheres and the *ifrms* was defined (when present) in chimpanzee hemispheres. Since the *mcgs* and *spls* are present in the PCC of nonhuman hominoids^46^, as in humans, an indentation was labeled as the *ifrms* if it was i) inferior to the *mcgs* and ii) anterior to the *spls*. The specific procedure for sulcal labeling was identical to the methodology described for defining sulci in young adults.

#### Sulcal depth and cortical thickness

Raw values (in mm) for sulcal depth, measured as the distance from the fundus to the smoothed outer pial surface, were calculated with the custom-modified algorithm used in the previous section^57^. These depth values were then normalized to the maximal depth within each individual hemisphere, which is located within the insula for both species^43^. Mean cortical thickness values (in mm) for each sulcus were calculated with the *mris_anatomical_stats* FreeSurfer function and also normalized to the thickest vertex within each individual hemisphere^58^. Similar to previous work^43^, we also demonstrate that the observed sulcal morphological patterns hold regardless of whether raw or normalized values are used (**Supplementary Fig. 7.2**). Like in young adults, the depth (mm), mean cortical thickness values (mm), and mean surface area (mm^2^) were also extracted from all juvenile and healthy older adult human PMC sulci.

#### Quantitatively assessing the developmental and comparative differences of the ifrms morphology

A three-way ANOVA with factors hemisphere (*left, right*), age group (*juvenile, young adult, older*), and species (*human, chimpanzee*) was run for each morphological feature. To address age differences in the chimpanzee dataset, the chimpanzees who had an *ifrms* were divided into similar age ranges reflecting the respective ranges across the human datasets: juvenile chimpanzees (age < 22), young adult chimpanzees (22 <= age <= 36), and older chimpanzees (age > 36). Three chimpanzees did not have ages provided and were therefore excluded from the morphological analyses.

### Identifying the ifrms in post-mortem hominoid and non-human primate brains

To determine if the *ifrms* is present in non-human primates and non-human hominoids, beyond the *in vivo* chimpanzee participants, we identified indentations in the cerebral cortex (when present) inferior to the *mcgs* and anterior to the *spls*, within images of Old World monkey, New World monkey, and non-human hominoid (chimpanzees, gorillas, orangutans) post-mortem hemispheres from a classic atlas by Retzius^68^ (**Supplementary Fig. 8**).

### Statistical methods

All statistical tests were implemented in R (v4.0.1) and RStudio (v1.3.959). Chi squared tests were carried out with the *chisq.test* function and post-hoc pairwise comparisons were run with the *chisq.multcomp* function from the built-in R *stats* and *RVAideMemoire* R packages, respectively. All ANOVAs (regular and repeated measures) were implemented using the *aov* function from the built-in R *stats* package. Post-hoc t-tests were conducted for the significant ANOVA effects and implemented with the *emmeans* R package. Effect sizes for the ANOVA effects are reported with the *generalized* eta-squared (η^2^G) metric and computed with the *anova_summary* function from the *rstatix* R package. The paired t-test was carried out with the *t.test* function from the built-in R *stats* package. The effect size of this t-test is reported with the Cohen’s d (*d*) metric and obtained with the *cohens_d* function from the *rstatix* R package. Linear regression analyses were run using the *lm* function from the built-in R *stats* package.

## Data availability

All data used for this project have been made freely available on GitHub: https://github.com/cnl-berkeley/stable_projects/tree/main/TripartiteLandmark_PosteriorCingulate. The processed data required to perform all statistical analyses and to reproduce all figures are freely available at the above link. Visualizations of all sulcal definitions generated for each participant are provided in the Supplementary Information. Requests for further information or raw data should be directed to the Corresponding Author, Kevin Weiner.

## Code availability

We provide the custom code to perform statistical analyses and to generate all figures on GitHub in an implementable R markdown file. (https://github.com/cnl-berkeley/stable_projects/tree/main/TripartiteLandmark_PosteriorCingulate).

## Acknowledgements

This research was supported by a T32 HWNI training grant and an NSF-GRFP fellowship (Voorhies), start-up funds from UC Berkeley (Weiner) and NIH R01MH116914 (Foster). Funding for some of the original data collection and curation was provided by NINDS R01 NS057156 (Bunge, Ferrer) and NSF BCS1558585 (Bunge, Wendelken). Data analysis was supported by NICHD R21HD100858 (Weiner, Bunge) and NSF CAREER 2042251 (Weiner). We thank Seth Koslov for assistance with *Neurosynth* data figures. We also thank Bennett Landman for his comments and feedback on the deep learning component. Alzheimer’s Disease Neuroimaging Initiative: Data collection and sharing for this project was funded by the Alzheimer’s Disease Neuroimaging Initiative (ADNI) (National Institutes of Health Grant U01 AG024904) and DOD ADNI (Department of Defense award number W81XWH-12-2-0012). ADNI is funded by the National Institute on Aging, the National Institute of Biomedical Imaging and Bioengineering, and through generous contributions from the following: AbbVie, Alzheimer’s Association; Alzheimer’s Drug Discovery Foundation; Araclon Biotech; BioClinica, Inc.; Biogen; Bristol-Myers Squibb Company; CereSpir, Inc.; Cogstate; Eisai Inc.; Elan Pharmaceuticals, Inc.; Eli Lilly and Company; EuroImmun; F. Hoffmann-La Roche Ltd and its affiliated company Genentech, Inc.; Fujirebio; GE Healthcare; IXICO Ltd.; Janssen Alzheimer Immunotherapy Research & Development, LLC.; Johnson & Johnson Pharmaceutical Research & Development LLC.; Lumosity; Lundbeck; Merck & Co., Inc.; Meso Scale Diagnostics, LLC.; NeuroRx Research; Neurotrack Technologies; Novartis Pharmaceuticals Corporation; Pfizer Inc.; Piramal Imaging; Servier; Takeda Pharmaceutical Company; and Transition Therapeutics. The Canadian Institutes of Health Research is providing funds to support ADNI clinical sites in Canada. Private sector contributions are facilitated by the Foundation for the National Institutes of Health (www.fnih.org). The grantee organization is the Northern California Institute for Research and Education, and the study is coordinated by the Alzheimer’s Therapeutic Research Institute at the University of Southern California. ADNI data are disseminated by the Laboratory for Neuro Imaging at the University of Southern California.

