## Supplementary Materials for "A new tripartite landmark in posterior cingulate cortex"

### Supplementary Results

#### Algorithmically determined sulcal depth vs. post-mortem values

To relate algorithmically determined depth (Methods[1]) of posteromedial (PMC) sulci *in-vivo* compared to previously published manual depths of PMC sulci in post-mortem samples, we compared the ranges of algorithmically determined sulcal depth values (in mm) in the present study to post-mortem depth values (in mm) collected by Ono *et al.*[2]. In their work, Ono and colleagues[2] analyzed the sulcal anatomy of 25 autopsy specimens (sex and age information was not available). Of interest to the present study, the authors computed the depth ranges of three PMC sulci analyzed in the present work: the marginal ramus of the cingulate sulcus (*mcgs*), splenial sulcus (*spls*), and parieto-occipital suclus (*pos*)[2]. By comparing the range of the sulcal depths of the *mcgs, spls,* and *pos* (in a subset of 25 participants in each sample), we found that the algorithmically determined depth values coincided with the range of values obtained by Ono *et al.*[2] in the left (*mcgs* = 2-17, *spls* = 3-13, *pos* = 12-33) and right hemisphere (*mcgs* = 4-21, *spls* = 2-18, *pos* = 17-40) in both the discovery (left: *mcgs* = 12.1-18.4, *spls* = 7.6-14.9, *pos* = 15.4-23.6; right: *mcgs* = 11.7-17.3, *spls* = 6.4-14, *pos* = 17.4-25.8) and replication (left: *mcgs* = 12-18.6, *spls* = 9.7-16.1, *pos* = 11.9-23.6; right: *mcgs* = 6.9-17.6, *spls* = 6.3-13.9, *pos* = 15.2-24.3) samples. This supports the accuracy of the depth estimations obtained by the algorithm on PMC sulci.

#### PMC sulci are morphologically distinct

To statistically compare the raw depths (in mm) of every PMC sulcus, we ran 2-way ANOVAs with sulcus (*11 PMC sulci*) and hemisphere (*left, right*) as factors for both the discovery and replication samples.

*Discovery sample:*

We observed a main effect of sulcus (F(10, 706) = 268.58, $p < 0.001$, $\eta^2G = 0.79$) and a trending effect of hemisphere (F(1, 706) = 3.67, $p = 0.056$, $\eta^2G = 0.005$), in which sulci in the left hemisphere were deeper (**Supplementary Fig. 2.1a**). Post hoc tests on the former effect revealed three takeaways regarding the PRC sulci. First, the three *prcus* sulci were the shallowest of the PRC sulci, but deeper on average than the putative PCC tertiary sulci (*p*-values < 0.001, Tukey's adjustment; **Supplementary Fig. 2.1a**). Second, the *prcus-p* was shallower than *prcus-i* and *prcus-a* (*p*-values < 0.001), while *prcus-i* and *prcus-a* had comparable depths ($p > 0.05$, Tukey's adjustment; **Supplementary Fig. 2.1a**). Third, the *prculs* was shallower than the *pos* ($p < 0.001$, Tukey's adjustment; **Supplementary Fig. 2.1a**).

*Replication sample:* Once again, main effects of sulcus (F(10, 702) = 302.94, $p < 0.001$, $\eta^2G = 0.81$) and hemisphere were observed (F(1, 702) = 19.83, $p < 0.001$, $\eta^2G = 0.03$). For the latter main effect, the PMC sulci were once again deeper in the left hemisphere on average (**Supplementary Fig. 2.1b**). Post hoc tests on the former main effect confirmed the three main findings in the discovery sample (**Supplementary Fig. 2.1b**). Lastly, there was an interaction between sulcus and hemisphere (F(10, 702) = 2.17, $p = 0.02$, $\eta^2G = 0.03$). Post hoc analyses indicated that the effect was driven by the *mcgs* and *prcus-a* being significantly deeper in the left hemisphere (*p*-values < 0.05, Tukey's adjustment; **Supplementary Fig. 2.1b**).

**Connectivity fingerprints of the *ifrms* and *spls* differ by hemisphere**

In addition to the sulcus x network interaction discussed in the main text, we also observed a sulcus x network x hemisphere interaction in both samples (discovery: $F(5, 175) = 3.27$, $p = 0.007$, $\eta^2G = 0.02$; replication: $F(5, 165) = 8.51$, $p < 0.001$, $\eta^2G = 0.04$). In the discovery sample, i) the *ifrms* overlapped more with DMN-a in the left hemisphere than the right ($p = 0.002$, Tukey's adjustment; **Fig. 4b,** left), ii) the *spls* with the CCN-b in the right hemisphere than the left ($p = 0.002$, Tukey's adjustment; **Fig. 4b**, left), and iii) the *spls* with the DMN-b in the left hemisphere than the right ($p < 0.001$, Tukey's adjustment; **Fig. 4b**, right). These three findings were replicated in the replication sample (*p*-values < 0.001, Tukey's adjustment; **Supplementary Fig. 5.2**); however, the *ifrms* also overlapped more with CCN-c in the right hemisphere than the left ($p < 0.001$, Tukey's adjustment; **Supplementary Fig. 5.2**, right; **Fig. 4b**; **Supplementary Fig. 5.3** for the connectivity profiles of the *ifrms* and *spls* in all participants).

**The three prcus components are functionally distinct from each other**

Since this was the first time that three separate *prcus* sulcal components were defined within every hemisphere in a large sample, we tested if these sulci were also distinguishable based on their functional connectivity profiles. Thus, we ran a 3-way repeated measures ANOVA with sulcus (*prcus-p*, *prcus-i*, *prcus-a*), network (*17 resting-state networks*[3]), and hemisphere (*left*, *right*) as factors.

*Discovery sample:* We observed an interaction effect between sulcus and network ($F(32, 1120) = 27.98$, $p < 0.001$, $\eta^2G = 0.13$). Post hoc tests indicated that these three sulci differed in their overlap with the different default mode subnetworks. On the one hand, these sulci show a posterior to anterior decrease in the amount of overlap with DMN-a (*p*-values < 0.001, Tukey's adjustment; **Supplementary Fig. 5.4a**). On the other hand, the three *prcus* show a posterior to anterior increase in overlap with DMN-c (*p*-values < 0.01, Tukey's adjustment; **Supplementary Fig. 5.4a**). Each sulcus also overlapped with other networks that were not shared with the other two sulci (**Supplementary Fig. 5.4a**). *Prcus-p* also overlapped with CCN-b (*p*-values < 0.001, Tukey's adjustment). *Prcus-i* and *prcus-a* both overlapped more with dorsal attention network A than *prcus-p* (*p*-values < 0.001, Tukey's adjustment). *Prcus-a* also overlapped more with somatomotor network A (*p*-values < 0.01, Tukey's adjustment) and ventral attention network B (*p*-values < 0.001, Tukey's adjustment) than the other two *prcus* components, as well as overlapped more with ventral attention network A (*p* = 0.001, Tukey's adjustment) and visual network B (*p* = 0.03, Tukey's adjustment) than *prcus-p*. Altogether, the three *prcus* are functionally dissociable.

*Replication sample:* Here we also observed a sulcus x network interaction ($F(32, 1056) = 27.34$, $p < 0.001$, $\eta^2G = 0.2$). Post hoc tests revealed somewhat similar relationships to those observed in the discovery sample. Similar to the discovery sample, these sulci showed a posterior to anterior decrease in DMN-a overlap and a posterior to anterior increase in overlap with DMN-c (*p*-values < 0.001, Tukey's adjustment; **Supplementary Fig. 5.4b**). Each sulcus also overlapped with other networks than the DMN (**Supplementary Fig. 5.4b**). *Prcus-a* overlapped more with CCN-b than *prcus-i* (*p* = 0.04, Tukey's adjustment). Furthermore, *prcus-a* overlapped more with ventral attention network A, ventral attention network B, and visual network B than the other two sulci (*p*-values < 0.01, Tukey's adjustment). Finally, *prcus-i* and *prcus-a* overlapped more with dorsal attention network A than *prcus-p* (*p*-values < 0.05, Tukey's adjustment).

**Inframarginal cortical indentations in Old World monkeys, New World monkeys, and non-human hominoids: From dimple to tertiary sulcus**

To determine if cortical indentations below the *mcgs* were also present beyond our *in vivo* chimpanzee and human hemispheres, we leveraged Retzius' classic atlas[4] that contained photographs of post-mortem brains from Old and New World monkeys, as well as a variety of non-human hominoids (gorillas, orangutans, and chimpanzees). Here we found that a shallow dimple (which we refer to as an inframarginal dimple, *ifrmd)* was also variably present in 63.83% (30/47) of Old World monkey hemispheres and 40% (4/10) of New World monkey hemispheres, which is consistent with references to a posterior cingulate dimple in modern research mentioned in the main text[5,6]. The *ifrms* was also present in a majority of non-human hominoid hemispheres examined. Specifically, we could identify the *ifrms* in post-mortem chimpanzees (*Troglodytes Niger*; 83.33% (15/18 hemispheres)), gorillas (*Anthropopithecus Gorilla;* 75% (3/4 hemispheres), and orangutans (*Simia Satyrus*; 75% (6/8 hemispheres)) examined. Interestingly, when the *ifrmd* or *ifrms* was present, Retzius sometimes depicted it in the schematic without a label, while in others, he excluded it entirely. **Supplementary Fig. 8** contains some example hemispheres with the *ifrmd* or *ifrms* identified in these species. We also will include a collection of all inspected hemispheres on our lab website with the publication of this paper.

**On the historical use of the term "inframarginal"**

To our knowledge, throughout neuroanatomical history, a label of "inframarginal sulcus" has not been proposed previously. Nevertheless, from our historical analyses, "inframarginal convolution" or "gyrus inframarginalis inferior" was proposed in the 1800s. Specifically, Ecker[7] credited Huschke[8] for the inframarginal label. However, Huschke did not label the sulcus of interest in the present work. Instead, Huschke provided an alternative label (*gyrus inframarginalis*) for the Superior Temporal Gyrus (STG). In the description of the *Lobulus supra-marginalis*, Ecker writes (from the 1873 English translation[9]):

"This lobule lies between the lower end of the posterior central convolution and the upper end of the *fissura Sylvii*, and arises from the lower end of the former, which forms the posterior part of the operculum, then develops into a lobule, consisting of several convolutions, arched around the end of the *fissura Sylvii*, in order to become the lower boundary of this fissure as the *gyrus marginalis inferior* or *temporalis superior* ($T_1$)."

In the description of the gyrus temporalis superior, Ecker directly references Huschke when he writes[9]: "1. *Gyrus temporalis superior* (Huschke) *seu infra-marginalis*, upper temporal convolution ($T_1$)."

Together, these historical analyses reveal that the term "inframarginal" has been used to label a part of the cortex, the STG, but not the sulcus of interest in the present study. Finally, the term inframarginal convolution has been largely removed from the modern nomenclature[10,11] and therefore, will not be confusable with the name we propose for the present sulcus of interest in the human brain.

**Even Einstein has an ifrms**

Historically, there has been great interest in "rare" brains – whether from those who have assassinated political figures or from "geniuses"[12–14]. In terms of the latter, in the last few decades, several papers have been published regarding Albert Einstein's (**Supplementary Figure 11**) brain[15–21], including one which aimed to identify sulcal patterns that differed in Einstein compared to "typical" brains[15]. This study highlighted the sulcal pattern within PCC as "abnormal" compared to "typical" sulcal patterns. The authors write:

"(F) Figure 8 of the left medial surface of Einstein's brain with unusual features highlighted in yellow. The cingulate gyrus has a long unnamed sulcus, and the cingulate sulcus gives off four inferiorly directed branches (two of which are tiny), which suggest that the cingulate gyrus may be relatively convoluted. The cuneus appears to be unusually convoluted."

Upon inspection of the published images, we have been able to identify one of these "tiny" sulci as the *ifrms*, and the other as the putative tertiary sulcus labeled here as the *icgs-p* (**Supplementary Fig. 11**). An additional sulcus labeled "u" for unnamed sulcus, is an additional sulcus within the cingulate gyrus that was not explicitly quantified in the present study but that is rather common in individual hemispheres. We include this point because it stresses the importance of identifying all sulci – including shallow, tertiary sulci – of the cerebral cortex in order to accurately assess typicality and atypicality, as well as how individual differences in sulcal patterning relate to function, anatomy, and cognition in both typical and atypical brains. In this particular case, the omission of the *ifrms* and other "tiny" or shallow sulci in PCC in neuroanatomical atlases resulted in the inaccurate conclusion that this was a special feature of Einstein's brain. Instead, this sulcal patterning in Einstein's PCC is actually common in humans, and also in many chimpanzee brains, as we quantify in the present paper.

**a** depicted but unlabeled

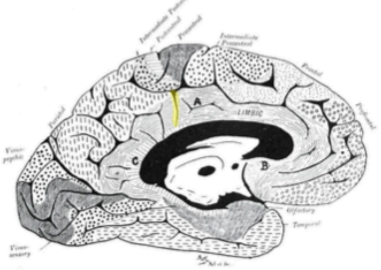

1905
*Campbell*

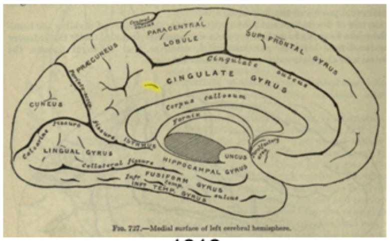

1918
*Gray*

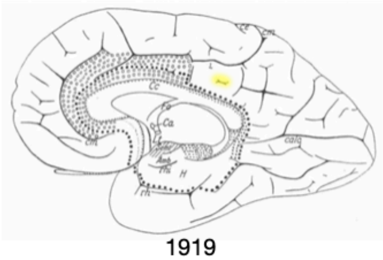

1919
*Vogt & Vogt*

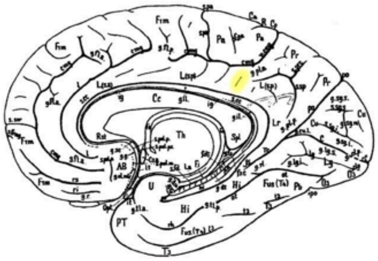

1925
*von Economo & Koskinas*

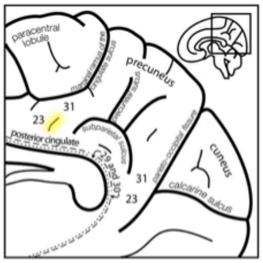

2009
*Margulies et al.*

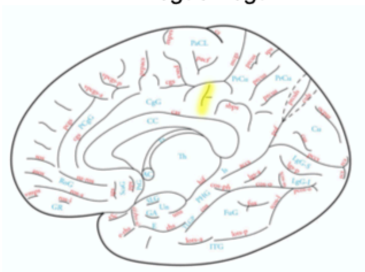

2019
*Petrides*

**b** dimple

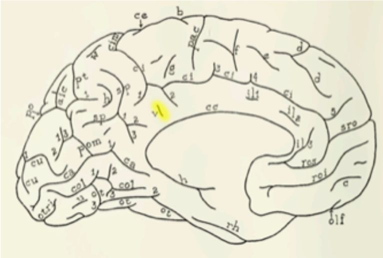

1951
*Bailey & Von Bonin*

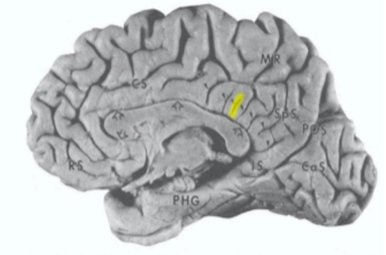

1993
*Vogt*

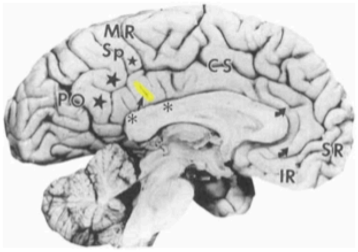

1995
*Vogt et al.*

**c** branch of cas or cgs

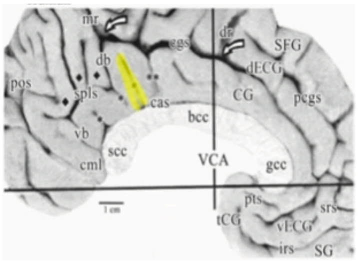

2009
*Vogt*

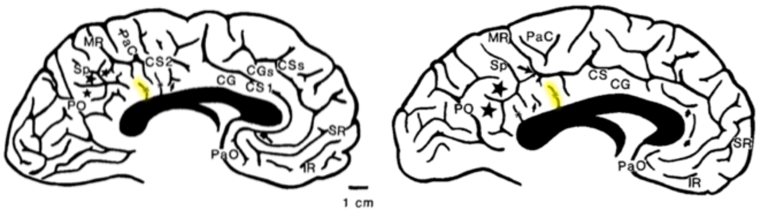

1995
*Vogt et al.*

**Supplementary Figure 1. Shallow PCC tertiary sulci depicted, but without formal names: A synopsis of historical and modern images**. It is important to note that, while this is the first time the *ifrms* and other shallow tertiary sulci were defined and labeled in a large sample, this is not the first time they have been depicted. **a**. Classic and modern studies have noted the presence of a cortical indentation below the *mcgs* but did not explicitly label it. Schematic illustrations from Campbell[22], Gray[23], Vogt and Vogt[24], von Economo and Koskinas[25], Marguiles *et al.*[26], and Petrides[27] are depicted. Yellow shading has been added to each of these images to indicate the location of the *ifrms* in the present study. **b**. In other situations, past research has also referred to the *ifrms* and the other shallow PCC tertiary sulci as inconsistent dimples. For example, Bailey and Von Bonin[28] referred to this indentation underneath the *mcgs* as "dimple v," while Vogt and colleagues[5,6] referred to it as one of many shallow dimples in PCC (arrowhead and arrow in the middle and right images, respectively). **c**. Finally, previous work has referred to cortical indentations underneath the *mcgs* as branches of the callosal sulcus (*cas*) or cingulate sulcus (*cgs*; asterisk[6,29]).

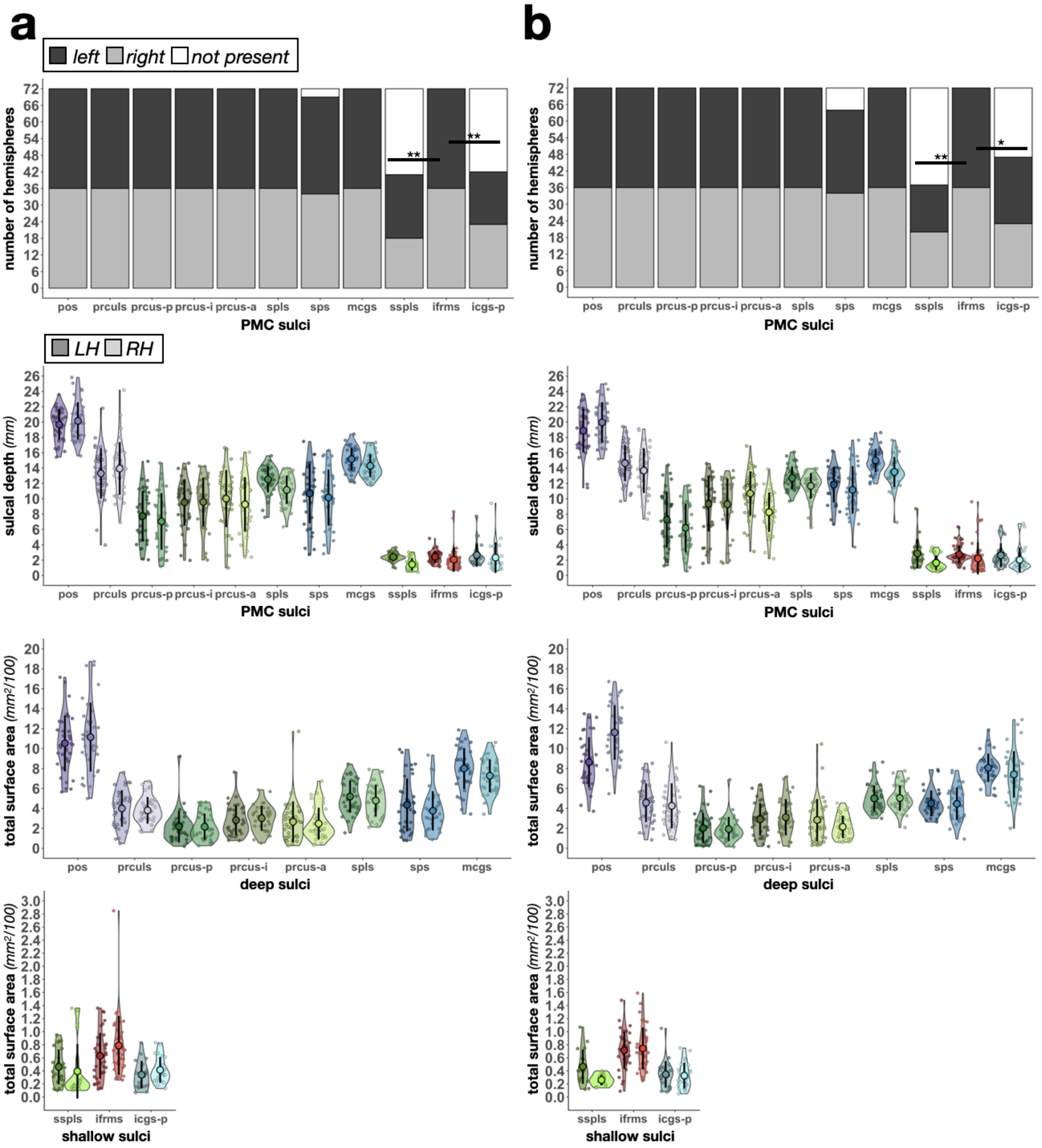


**Supplementary Figure 2.1. The *ifrms*, but not other shallow sulci in PCC, are identifiable in every hemisphere.** The same layout as Fig. 2b-e, but for all 11 PMC sulci. **a.** Incidence and morphology of PMC sulci in the discovery sample. **b.** Same as **a** but for the replication sample. First row: Stacked bar plots illustrate the incidence rates of three shallow sulci (*ifrms, sspls, icgs-p*) relative to the other manually defined PMC sulci ($N_{total}$ = 72 hemispheres each). Dark gray, light gray, and white indicate the number of hemispheres that contain that given sulcus (LH: dark gray; RH: light gray; white: absent). Asterisks indicate statistically significant incidence rates between the *ifrms* and the two other shallow sulci (*$p$<.05, **$p$<0.01; the same as in Fig. 2b and 2d). Second row: Sulcal depth (mm) plotted for each individual participant (small colored circles). The mean (large colored circles), standard deviation (black line), and kernel density estimate (colored violin) are also plotted for each sulcus. The PMC sulci are each colored according to the legend in Fig. 2a, with darker shades indicating LH values and lighter shades indicating RH values. Third row: Same as the second row but for the total surface area ($mm^2$) of the deep sulci. Note that these values are scaled down by a factor of 100. Fourth row: Same as the third row, but for the three shallow sulci.

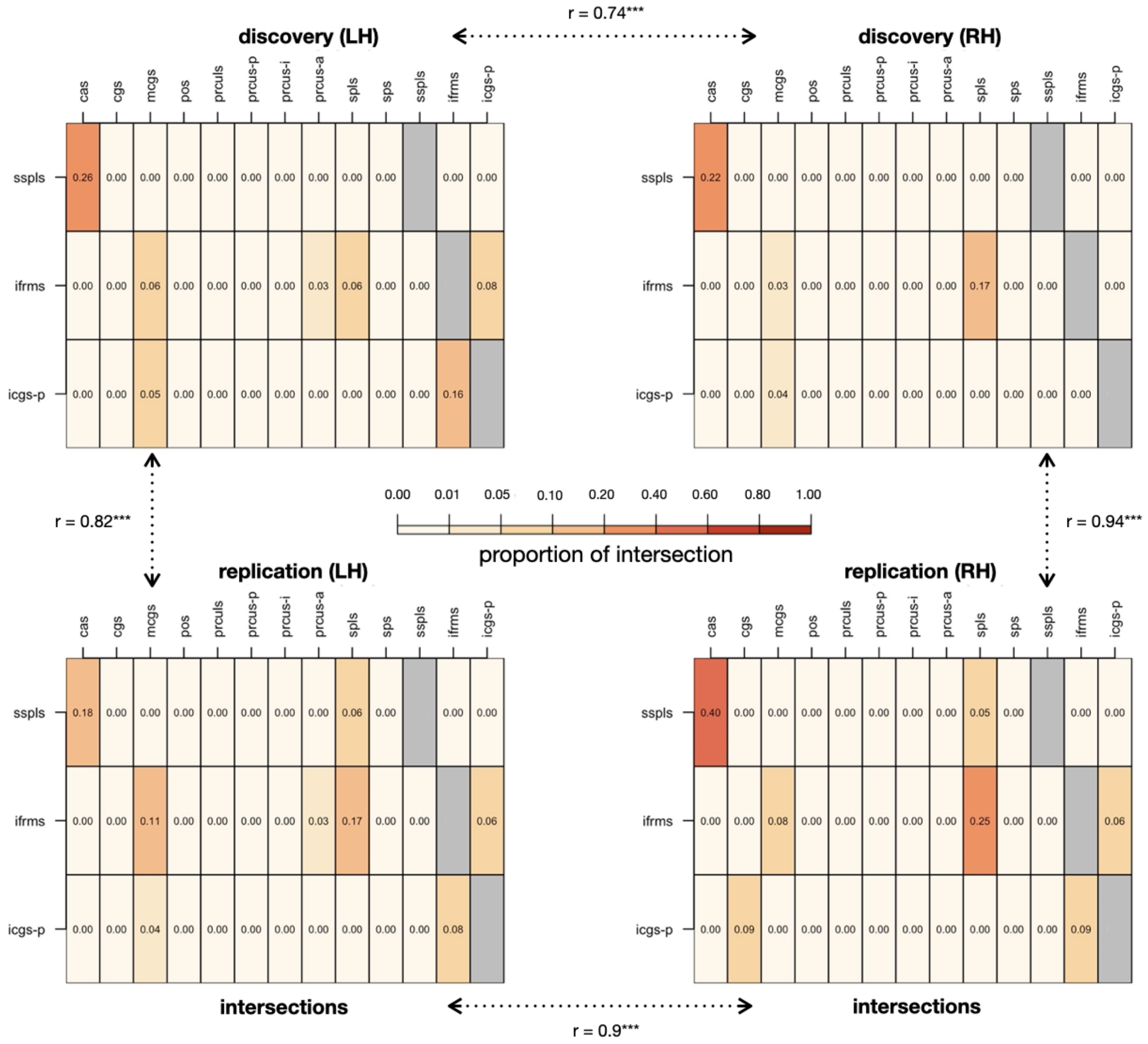


**Supplementary Figure 2.2. Intersections of shallow PCC sulci are similar between hemispheres and samples.** Rates of intersection with surrounding sulci were quantified for each PCC shallow sulcus to identify common sulcal patterns in each young adult sample. For each shallow PCC sulcus (*sspls, ifrms, icgs-p*), we report the proportion of intersection (frequency of occurrence/total number of observations) with each PMC sulcus. Note that the callosal sulcus (cas) and cingulate sulcus (cgs) were also included as the sspls intersected with the cas and the icgs-p intersected with the cgs frequently. Calculating the correlation between matrices shows that intersections of these sulci is comparable (all *r*s > .70; all *p*s < 0.001) between hemispheres and samples. The three most prevalent types for each shallow sulcus in each sample are included in Supplementary Tables 4 and 6.

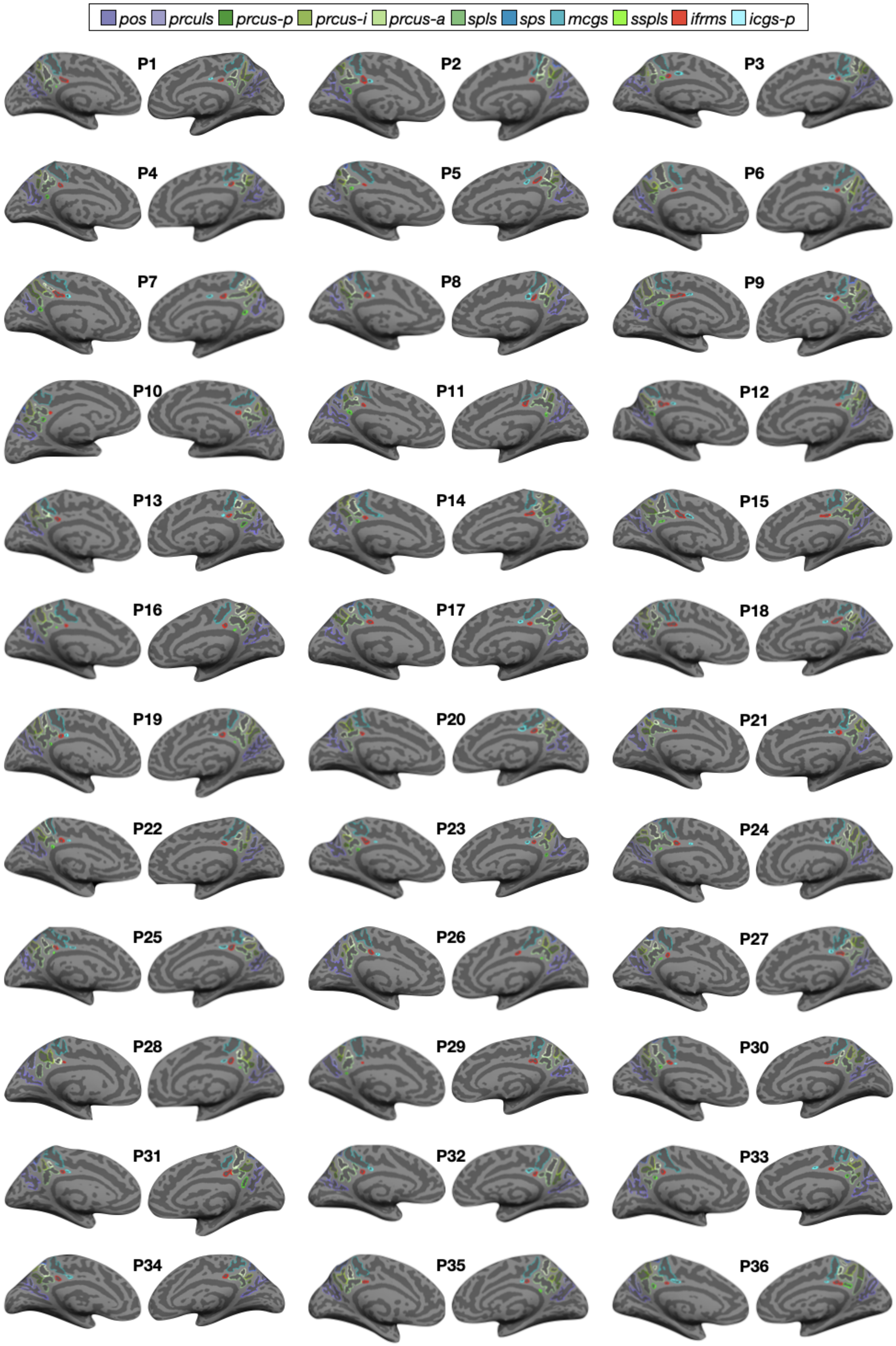
pos prculs prcus-p prcus-i prcus-a spls sps mcgs sspls ifrms icgs-p
P1 P2 P3 P4 P5 P6 P7 P8 P9 P10 P11 P12 P13 P14 P15 P16 P17 P18 P19 P20 P21 P22 P23 P24 P25 P26 P27 P28 P29 P30 P31 P32 P33 P34 P35 P36

**Supplementary Figure 3.1. Manual PMC sulcal labels in the left and right hemispheres of each participant in the discovery sample.** Each sulcus is displayed on the inflated cortical surface in FreeSurfer 6.0.0 and is colored according to the key at the top. Each hemisphere contains at least eight sulci (from posterior to anterior): *pos, prculs, prcus-p, prcus-i, prcus-a, spls, mcgs,* and *ifrms*. The *sps*, *sspls*, and *icgs-p* are all variably present.

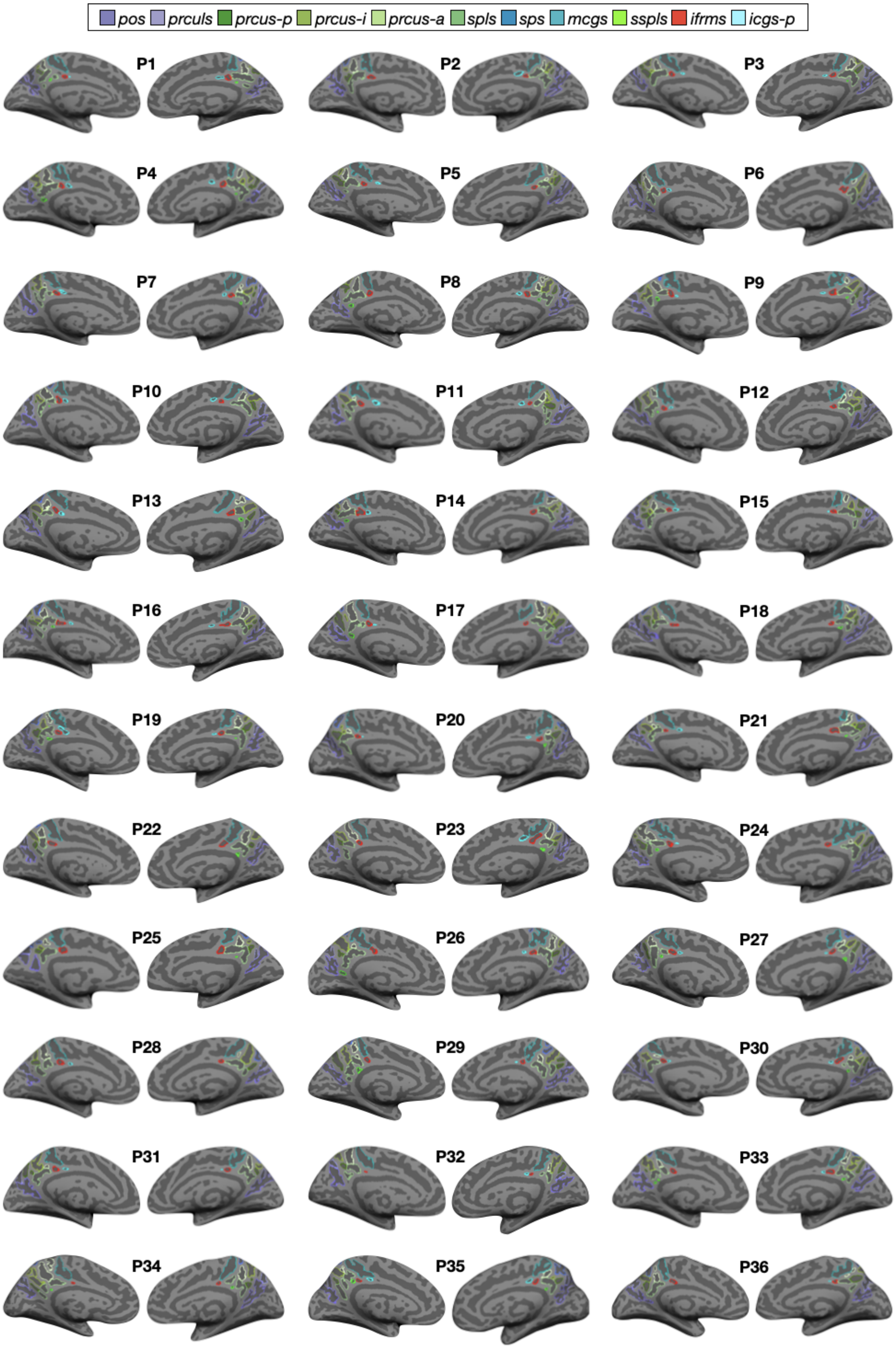
pos
prculs
prcus-p
prcus-i
prcus-a
spls
sps
mcgs
sspls
ifrms
icgs-p
P1
P2
P3
P4
P5
P6
P7
P8
P9
P10
P11
P12
P13
P14
P15
P16
P17
P18
P19
P20
P21
P22
P23
P24
P25
P26
P27
P28
P29
P30
P31
P32
P33
P34
P35
P36

**Supplementary Figure 3.2. Manual PMC sulcal labels in the left and right hemispheres of each participant in the replication sample.** Same layout as Supplementary Fig. 3.1, but for the *replication* sample.

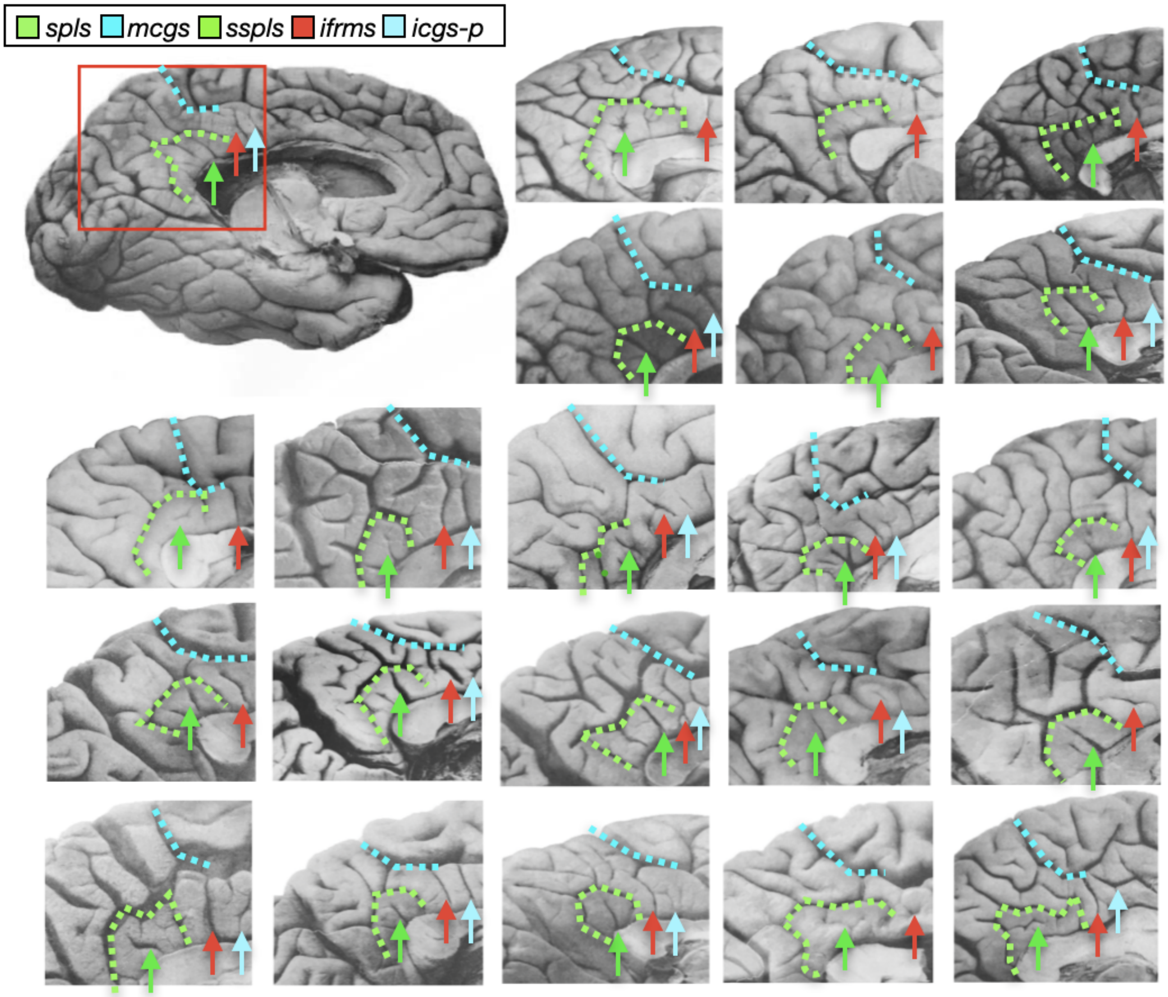


**Supplementary Figure 3.3. The *ifrms* is present in post-mortem hemispheres.** 22 post-mortem hemispheres (11 left hemispheres, 11 right hemispheres) labeled from a classic neuroanatomy atlas[30]. The *mcgs* and *spls* are labeled with colored dotted lines while the *ifrms* is identified with a red arrow. When present, the *sspls* and *icgs-p* are identified with green and cyan arrows, respectively. The *ifrms* is present in all hemispheres labeled while the other two shallow sulci have more variable appearances, which is consistent with our findings from *in-vivo* analyses.

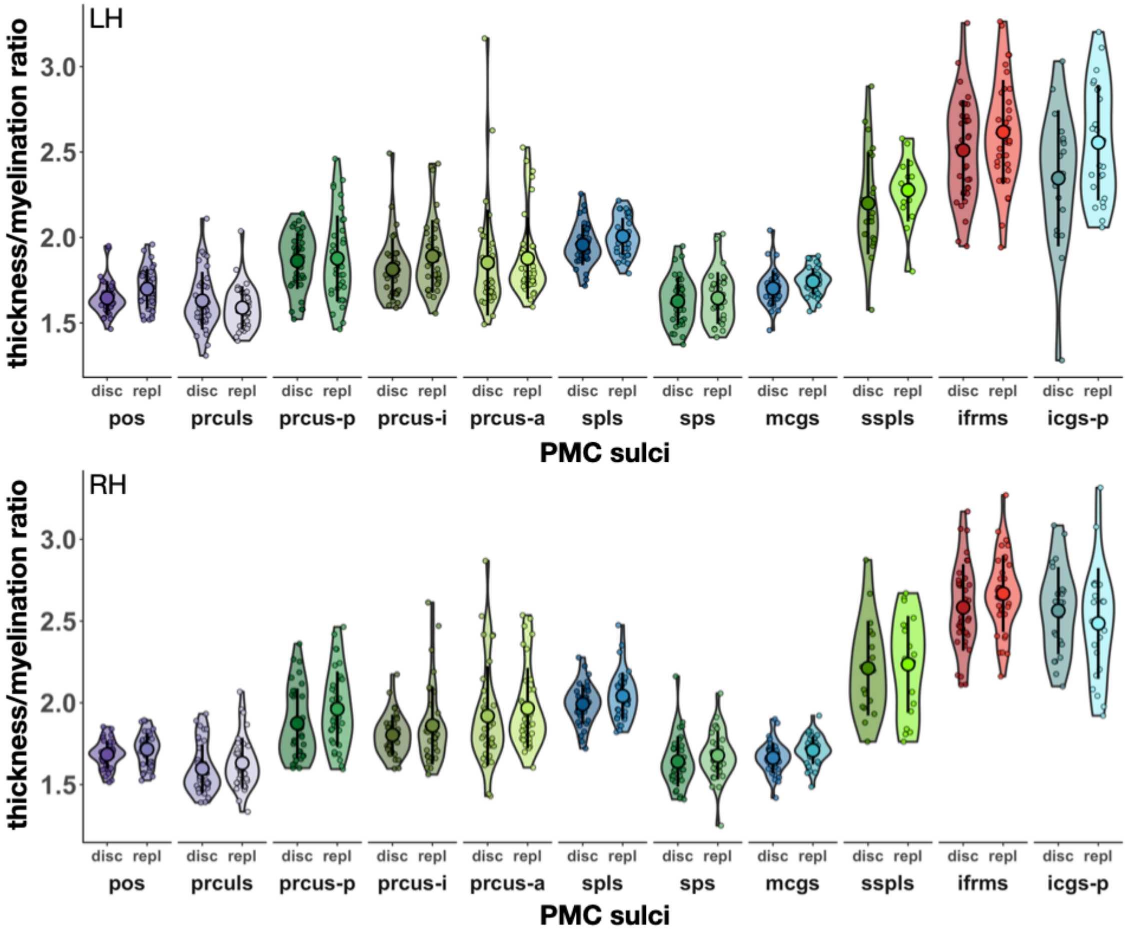


**Supplementary Figure 4.1**. **The *ifrms* is a macroanatomical and microanatomical landmark in PCC.** Same layout as Fig. 3b. Thickness/myelination ratio for all 11 PCC sulci in the discovery (disc) and replication (repl) samples in the LH (top) and RH (bottom). Individual participants from the discovery and replication samples (small colored circles), means (large colored circles), standard deviation (black line), and kernel density estimate (colored violins) are plotted for each sulcus. For all PCC sulci, the *ifrms* has the largest thickness/myelination ratio.

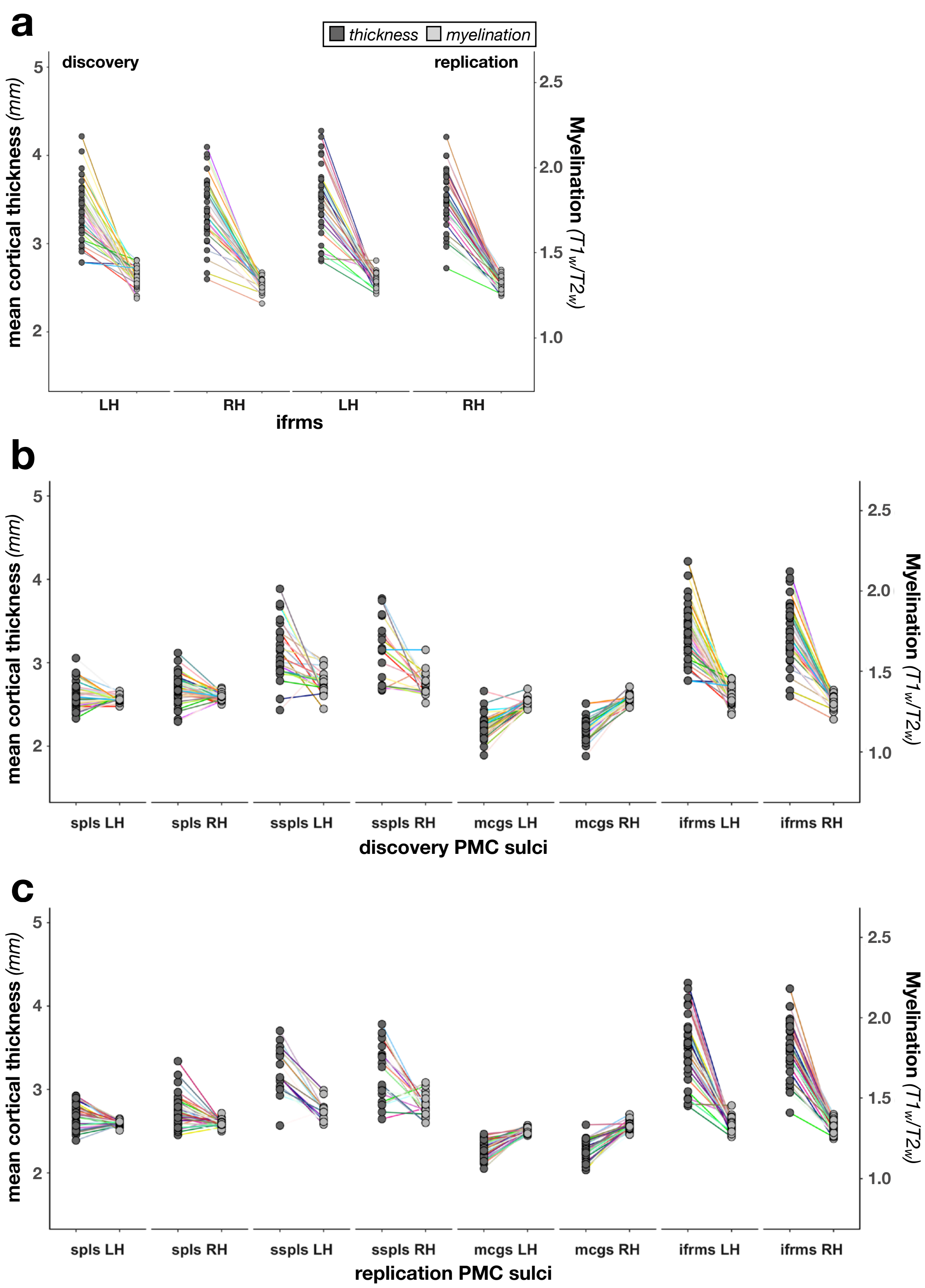


**Supplementary Figure 4.2**. **Individual cortical thickness and myelination values of PMC sulci. a.** Thickness (mm; left axis; dark gray) and myelination ($T_1$w/$T_2$w; right axis; light gray) values for the *ifrms* only in the discovery (left) and replication (right) samples in both the left (LH) and right hemispheres (RH). The thickness and myelination values for each individual participant (small circles) are plotted with a uniquely colored line connecting them. **b**. Similar layout as **a**, but also including the other three sulci (*spls*, *sspls*, and *mcgs*) analyzed in Fig. 3b (discovery sample only). **c.** Same layout as **b** but for the replication sample.

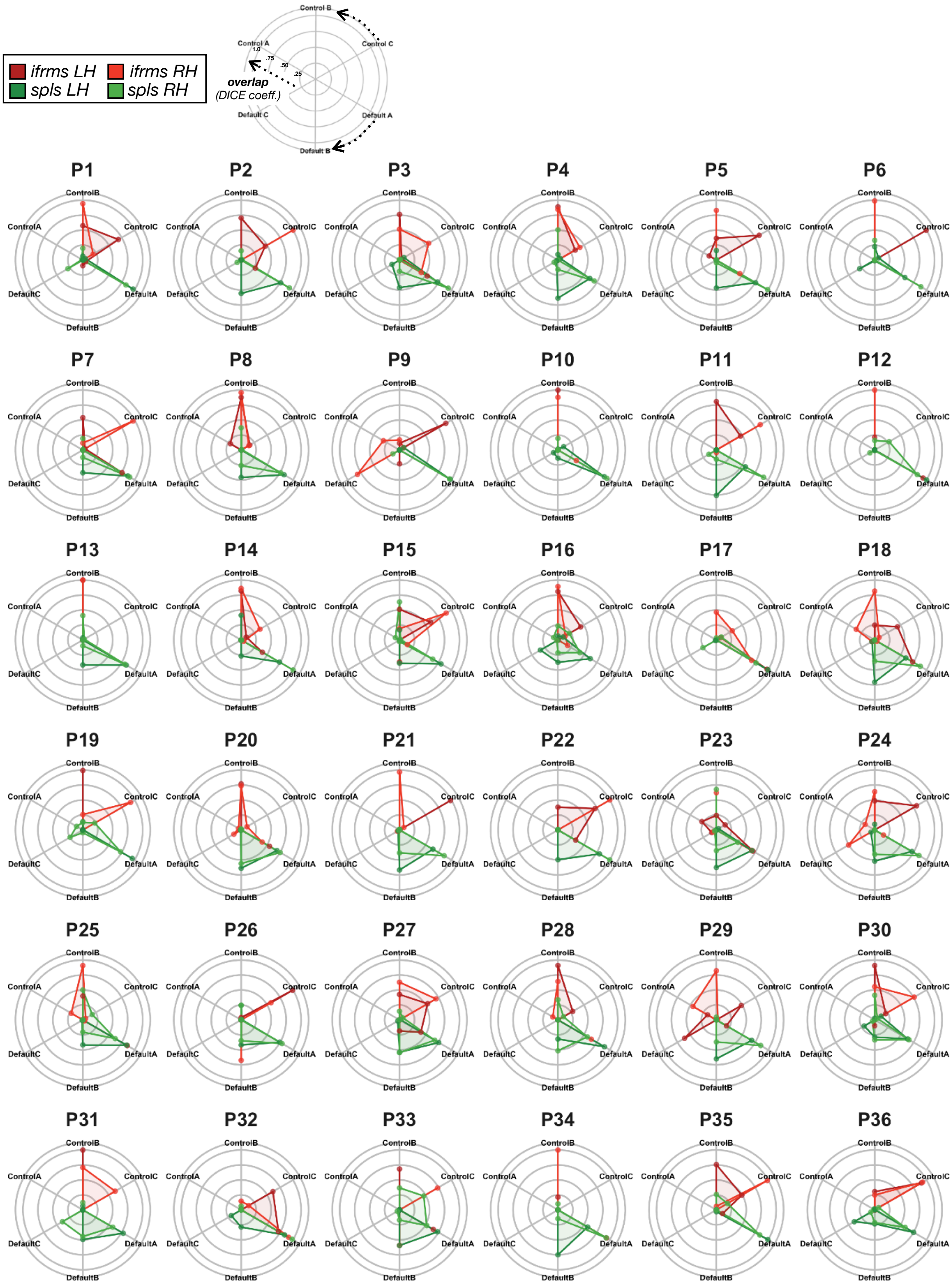
ifrms LH
ifrms RH
spls LH
spls RH
Control B
Control C
Control A
Default A
Default B
Default C
1.0
.75
.50
.25
overlap
(DICE coeff.)
P1
P2
P3
P4
P5
P6
P7
P8
P9
P10
P11
P12
P13
P14
P15
P16
P17
P18
P19
P20
P21
P22
P23
P24
P25
P26
P27
P28
P29
P30
P31
P32
P33
P34
P35
P36
ControlB
ControlC
ControlA
DefaultA
DefaultB
DefaultC

**Supplementary Figure 5.1. Individual participant connectivity fingerprints of the *ifrms* and *spls* in the discovery sample.** Top left: Legend for interpreting the polar plots. Arrows denote the direction of each network's overlap (CCN: top; DMN: bottom). The more the fingerprint extends to the periphery of the circle, the higher the dice coefficient. Individual participant resting state functional connectivity parcellations were obtained from a recent study[3], blind to cortical folding, and independent of our PMC sulcal definitions. The connectivity fingerprint represents the overlap of each network within a given sulcus. Bottom: Polar plots showing the connectivity fingerprints of the *ifrms* (red) and *spls* (green) in individual participants for the left hemisphere (LH, darker shade) and right hemisphere (RH, lighter shade) of the discovery sample. Solid lines: mean. Dashed lines: ±1 sem.

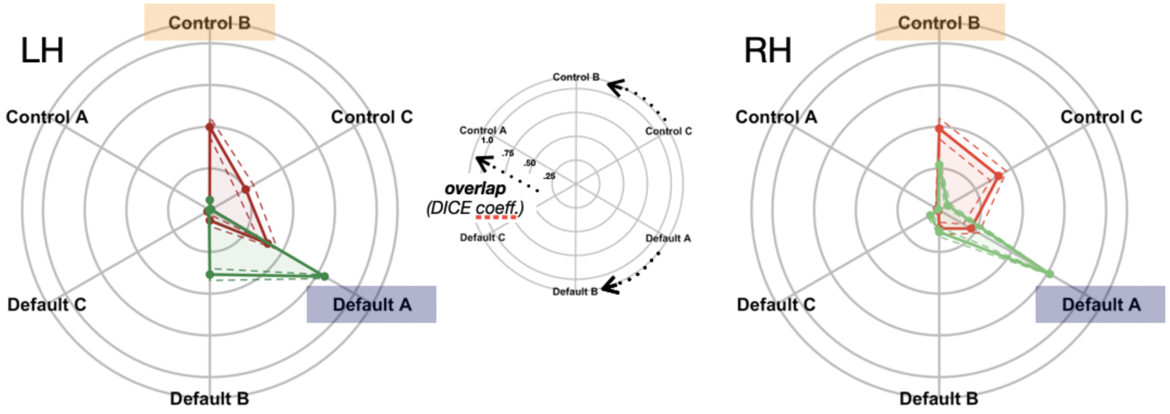


**Supplementary Figure 5.2. Mean connectivity fingerprints of the *ifrms* and *spls* in the replication sample.** Same layout as Fig. 4b, but for the replication sample. See Supplementary Fig. 5.3 for all individuals in this sample.

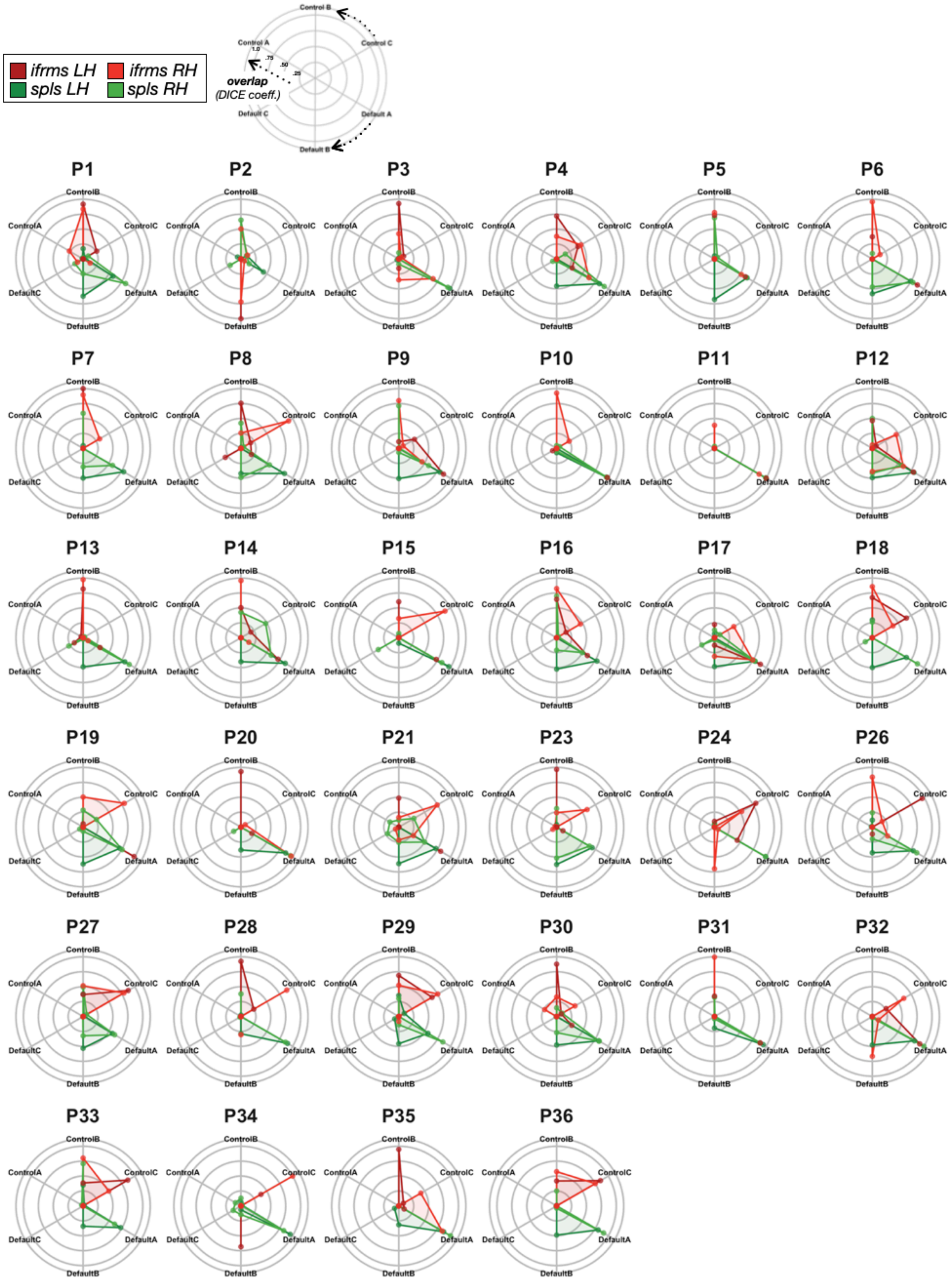
ifrms LH
ifrms RH
spls LH
spls RH
overlap
(DICE coeff.)
Control B
Control C
Control A
Default C
Default A
Default B
1.0
.75
.50
.25
P1
P2
P3
P4
P5
P6
P7
P8
P9
P10
P11
P12
P13
P14
P15
P16
P17
P18
P19
P20
P21
P23
P24
P26
P27
P28
P29
P30
P31
P32
P33
P34
P35
P36
ControlB
ControlC
ControlA
DefaultC
DefaultA
DefaultB

**Supplementary Figure 5.3. Individual participant connectivity fingerprints of the *ifrms* and *spls* in the replication sample.** Same layout as Supplementary Fig. 5.1, but for each individual in the replication sample. Note that two participants were excluded (P22 and P25) due to not having resting-state parcellations available.

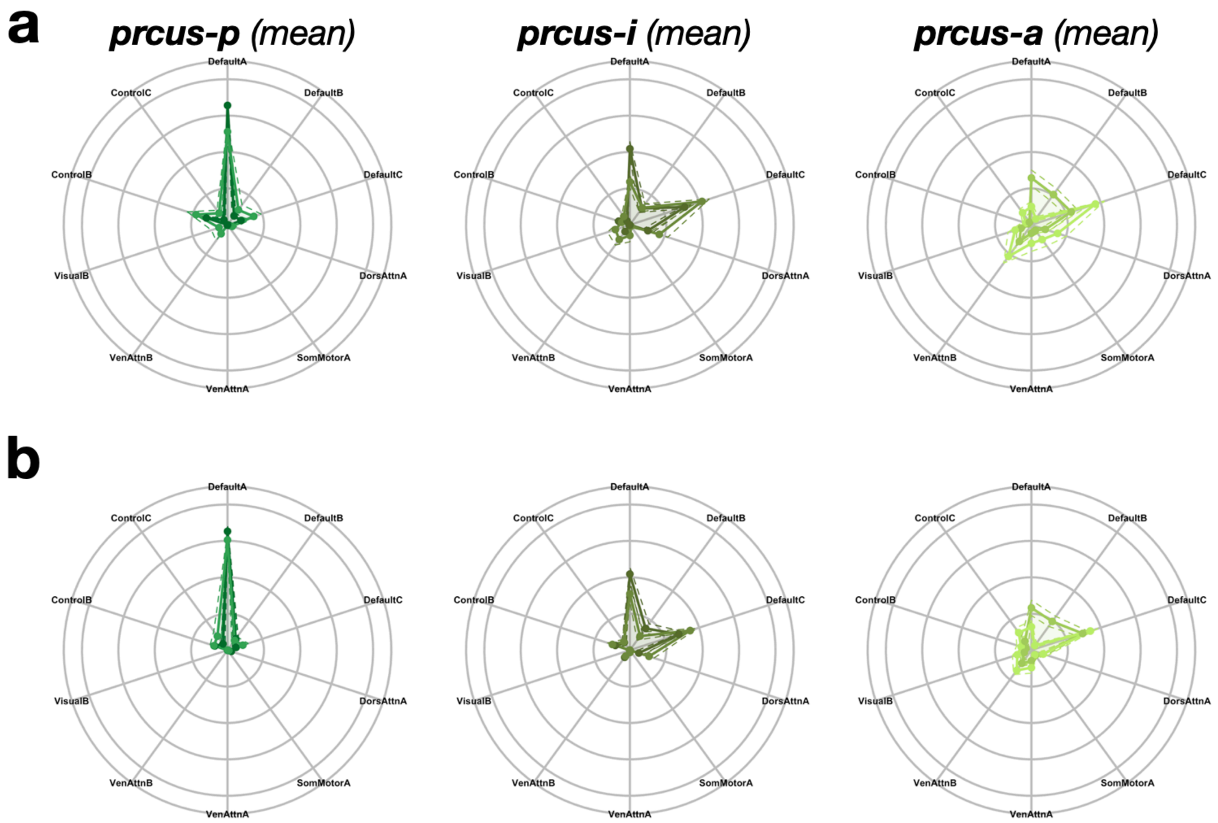


**Supplementary Figure 5.4. The three *prcus* sulci have different connectivity fingerprints. a.** The mean connectivity fingerprints of the three *prcus* sulci in the discovery sample. Polar plots visualize the mean connectivity fingerprint of the three *prcus* sulci (posterior to anterior) for both hemispheres. The polar plots follow the same layout as Fig. 4b and Supplementary Fig. 5.2. The solid-colored lines connect the means, and the dashed colored lines indicate ±1 sem. Each *prcus* component is colored according to the legend in Fig. 2a, with the darker shades indicating the left hemisphere (LH) and lighter shades indicating the right hemisphere (RH). **b.** The same layout as **a.**, but for the replication sample.

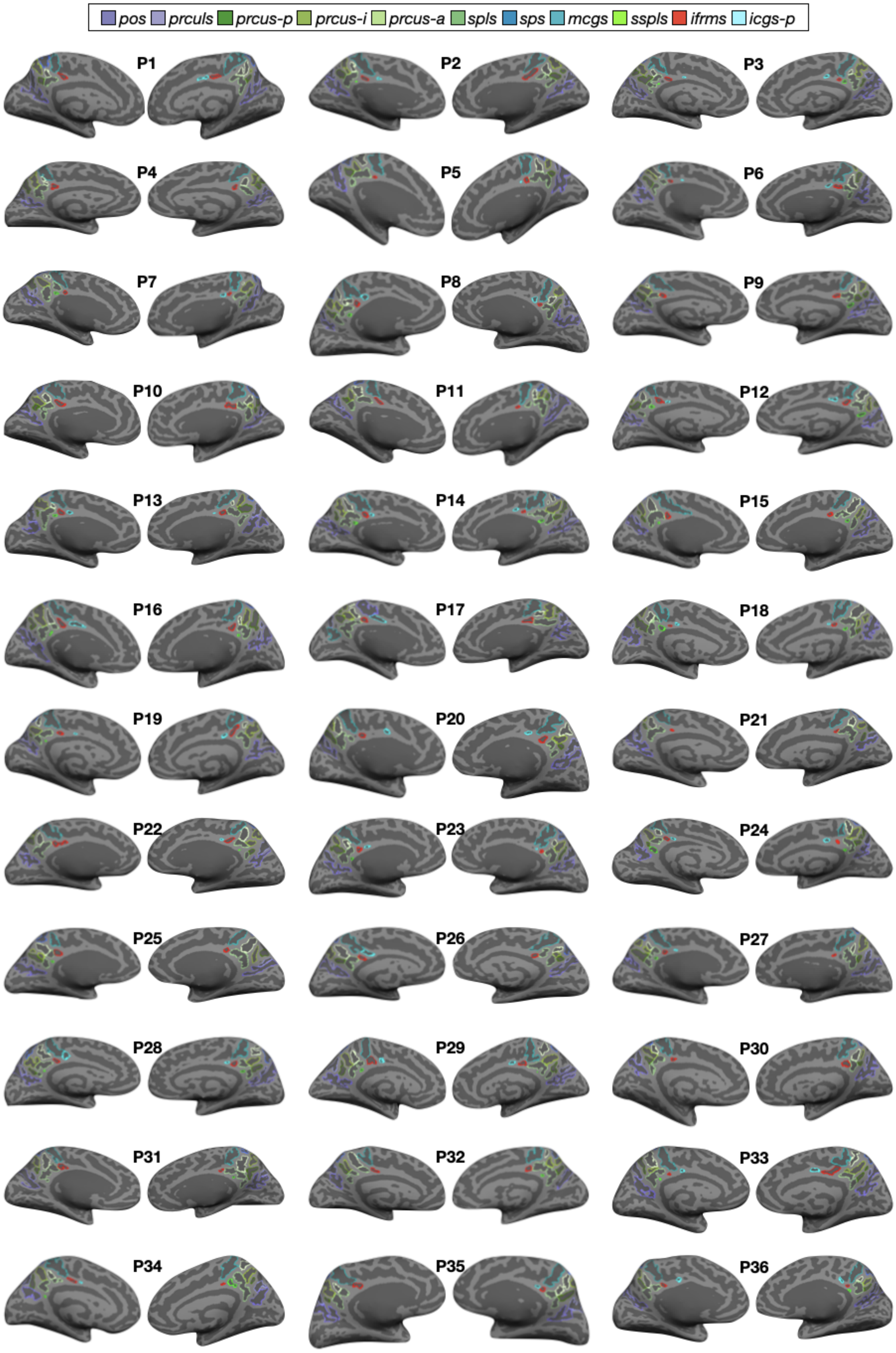
pos
prculs
prcus-p
prcus-i
prcus-a
spls
sps
mcgs
sspls
ifrms
icgs-p
P1
P2
P3
P4
P5
P6
P7
P8
P9
P10
P11
P12
P13
P14
P15
P16
P17
P18
P19
P20
P21
P22
P23
P24
P25
P26
P27
P28
P29
P30
P31
P32
P33
P34
P35
P36

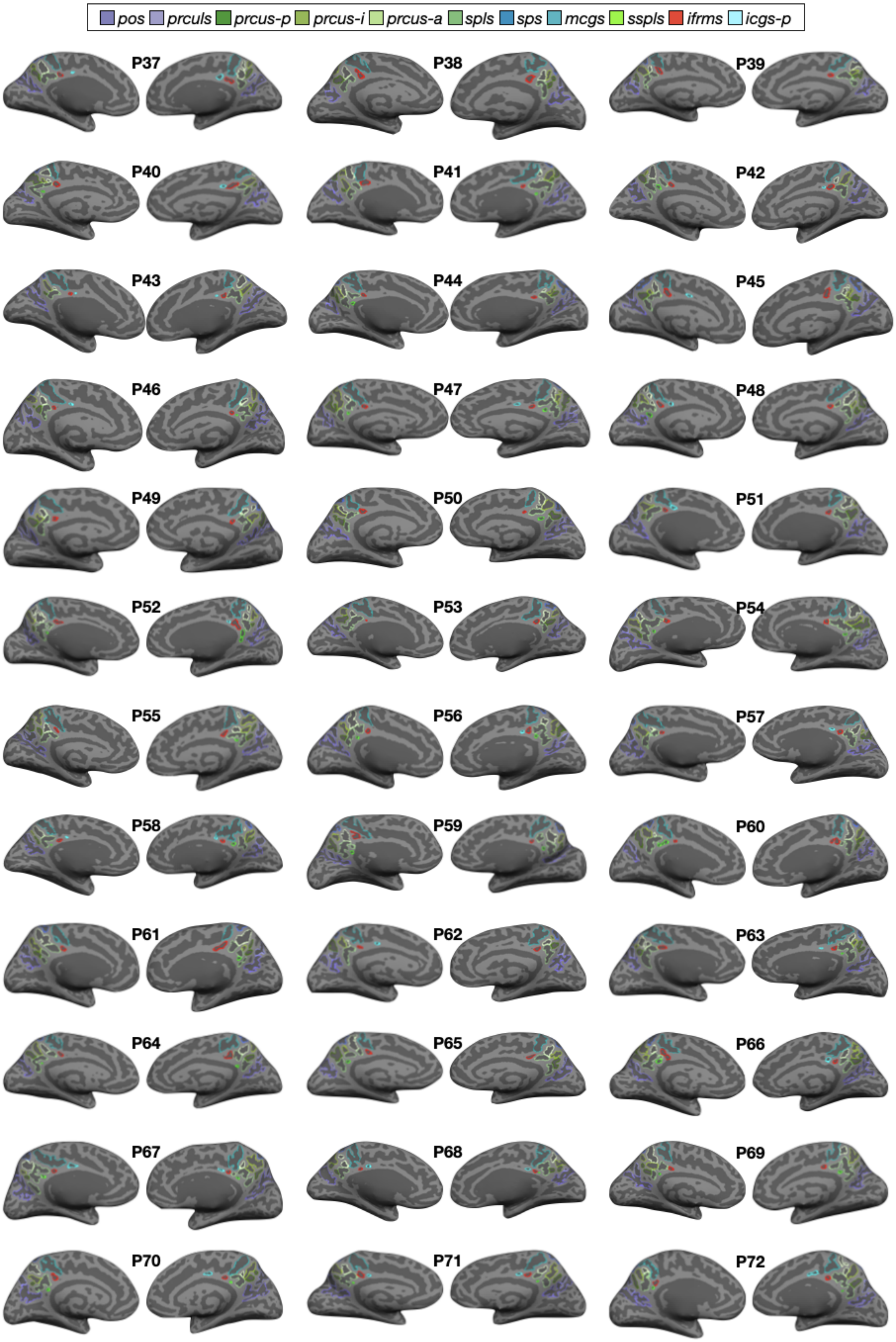
pos
prculs
prcus-p
prcus-i
prcus-a
spls
sps
mcgs
sspls
ifrms
icgs-p
P37
P38
P39
P40
P41
P42
P43
P44
P45
P46
P47
P48
P49
P50
P51
P52
P53
P54
P55
P56
P57
P58
P59
P60
P61
P62
P63
P64
P65
P66
P67
P68
P69
P70
P71
P72

**Supplementary Figure 6.1. Manual PMC sulcal labels in the left and right hemispheres of each human juvenile participant.** Same layout as Supplementary Fig. 3.1, but for the human juvenile participants included in the present study.

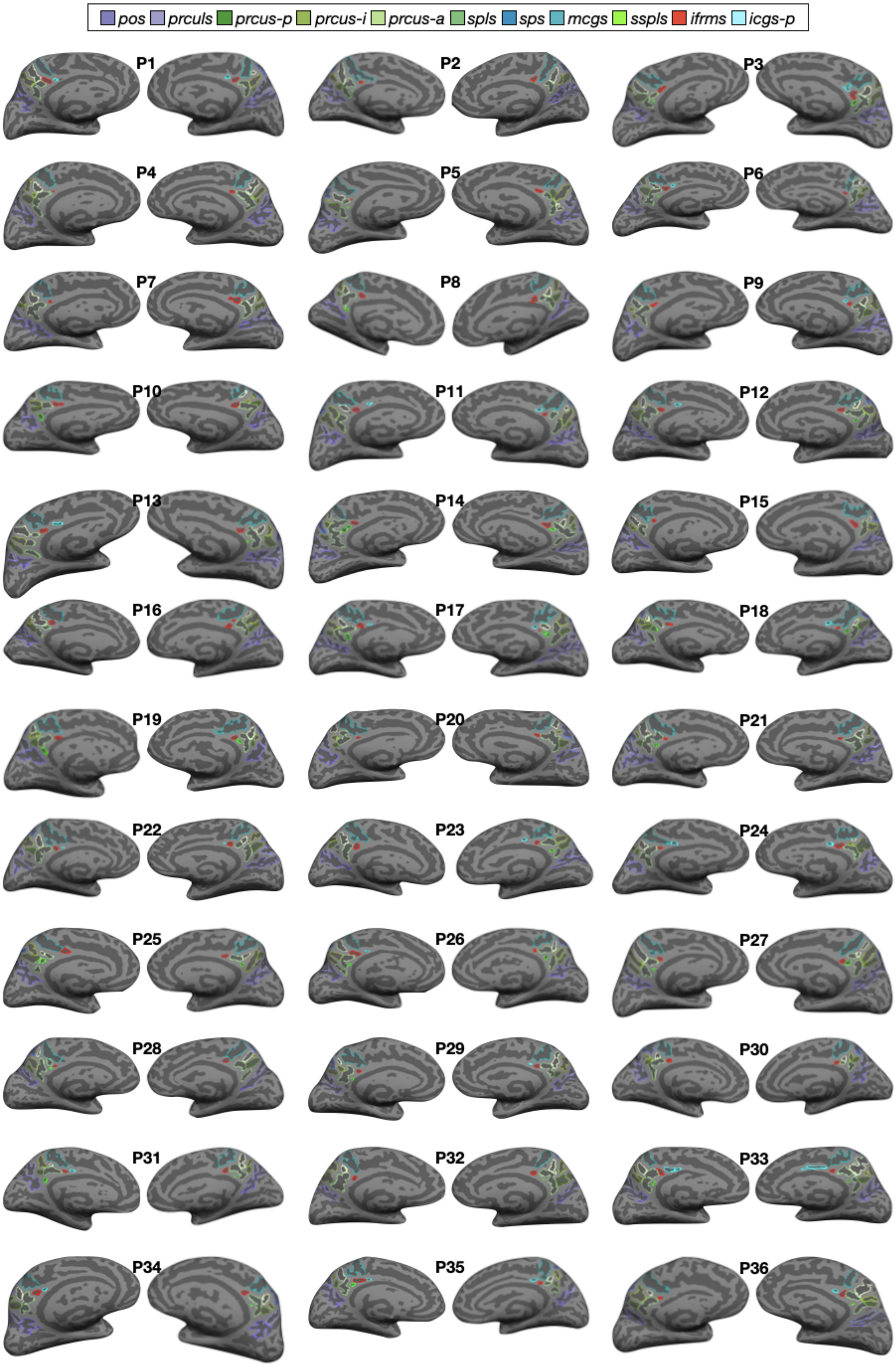

pos
prculs
prcus-p
prcus-i
prcus-a
spls
sps
mcgs
sspls
ifrms
icgs-p
P1
P2
P3
P4
P5
P6
P7
P8
P9
P10
P11
P12
P13
P14
P15
P16
P17
P18
P19
P20
P21
P22
P23
P24
P25
P26
P27
P28
P29
P30
P31
P32
P33
P34
P35
P36

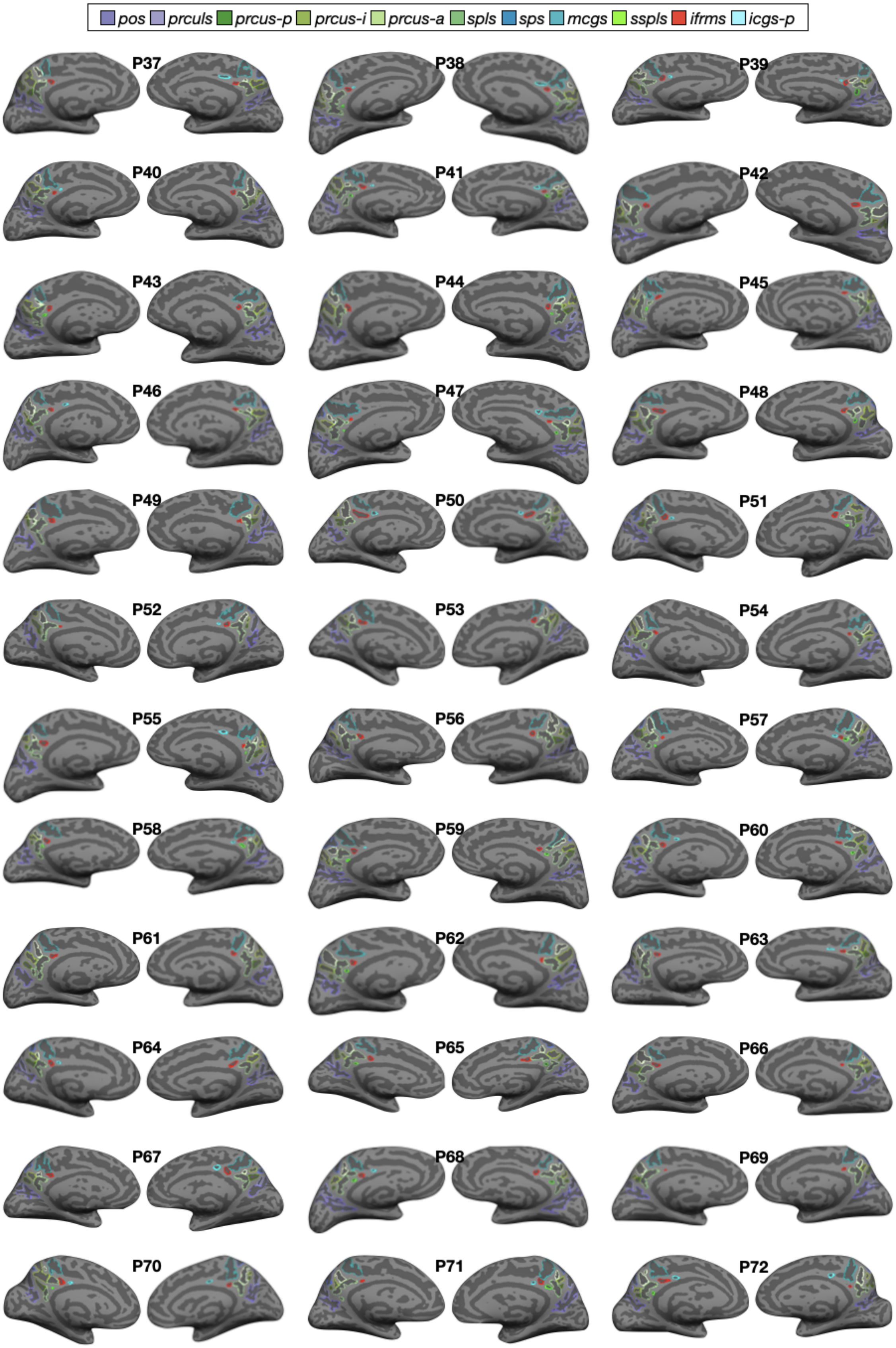
pos prculs prcus-p prcus-i prcus-a spls sps mcgs sspls ifrms icgs-p
P37
P38
P39
P40
P41
P42
P43
P44
P45
P46
P47
P48
P49
P50
P51
P52
P53
P54
P55
P56
P57
P58
P59
P60
P61
P62
P63
P64
P65
P66
P67
P68
P69
P70
P71
P72

**Supplementary Figure 6.2. Manual PMC sulcal labels in the left and right hemispheres of each human elderly participant.** Same layout as Supplementary Fig. 3.1, but for the human elderly participants included in the present study.

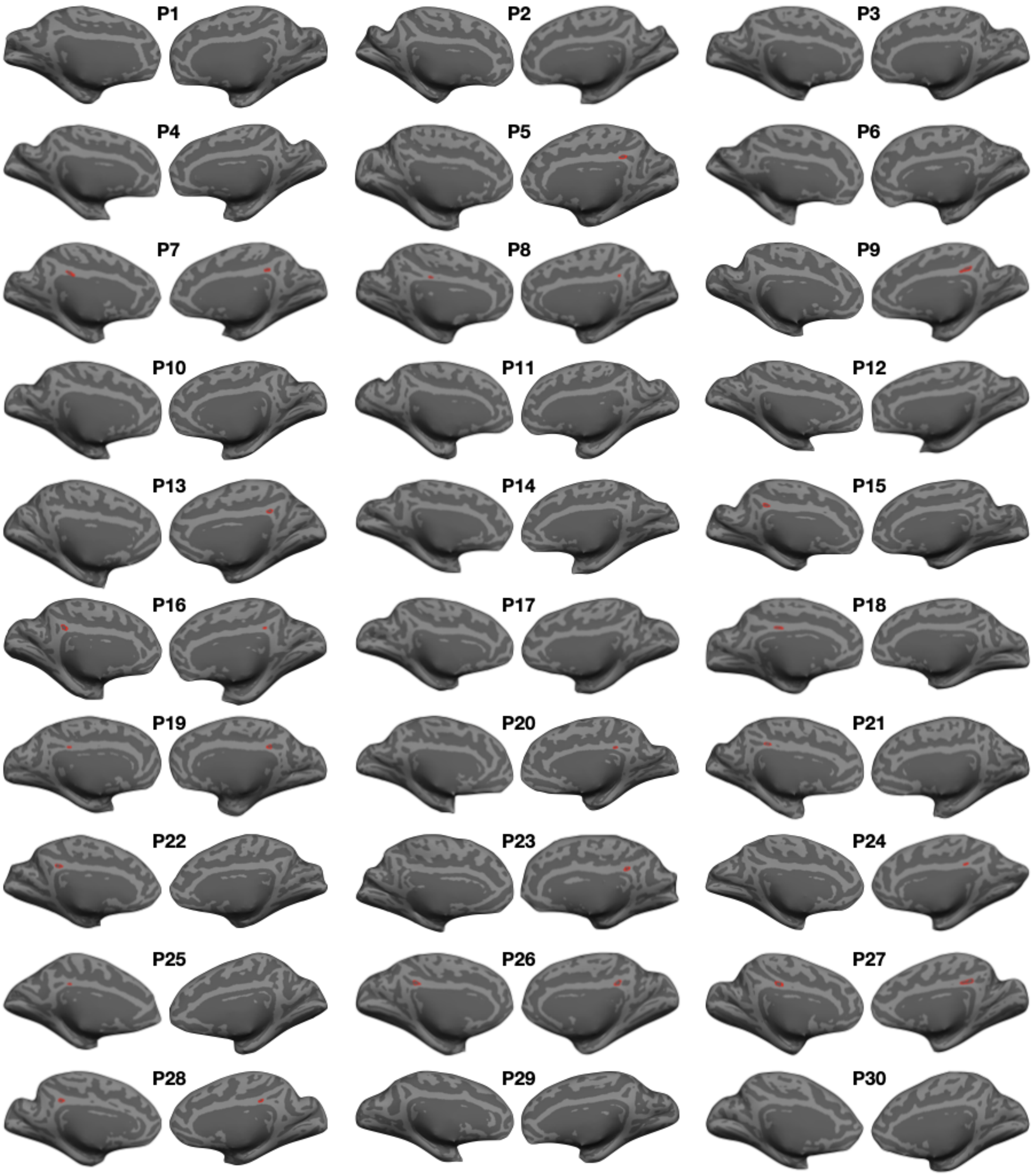
P1
P2
P3
P4
P5
P6
P7
P8
P9
P10
P11
P12
P13
P14
P15
P16
P17
P18
P19
P20
P21
P22
P23
P24
P25
P26
P27
P28
P29
P30

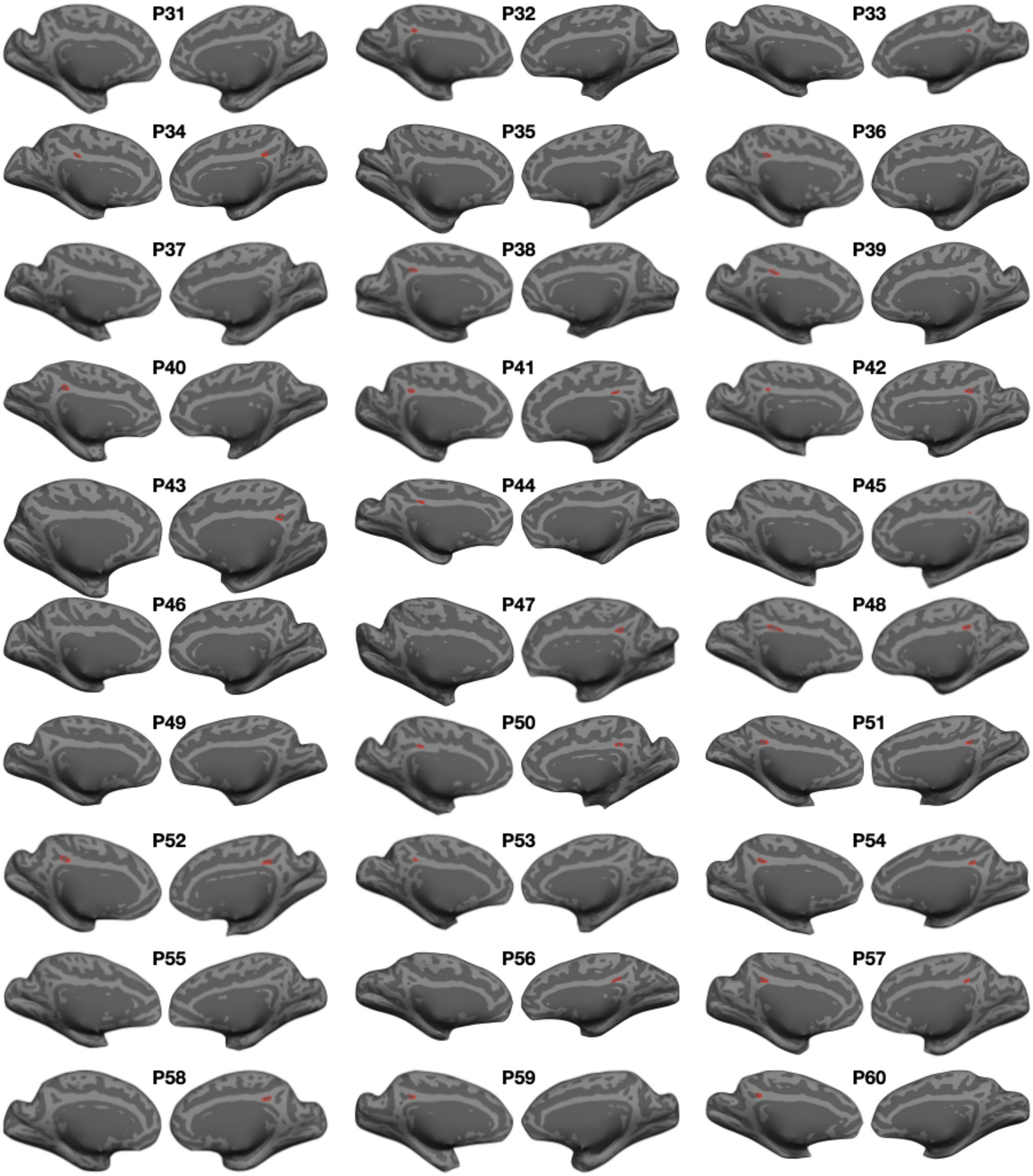


**Supplementary Figure 6.3. Manual *ifrms* labels in the left and right hemispheres in chimpanzees.** Same layout as Supplementary Fig. 3.1, but for each chimpanzee included in the present study. Unlike the human participants, we only labeled the *ifrms* (not all PMC sulci) when it was identifiable.

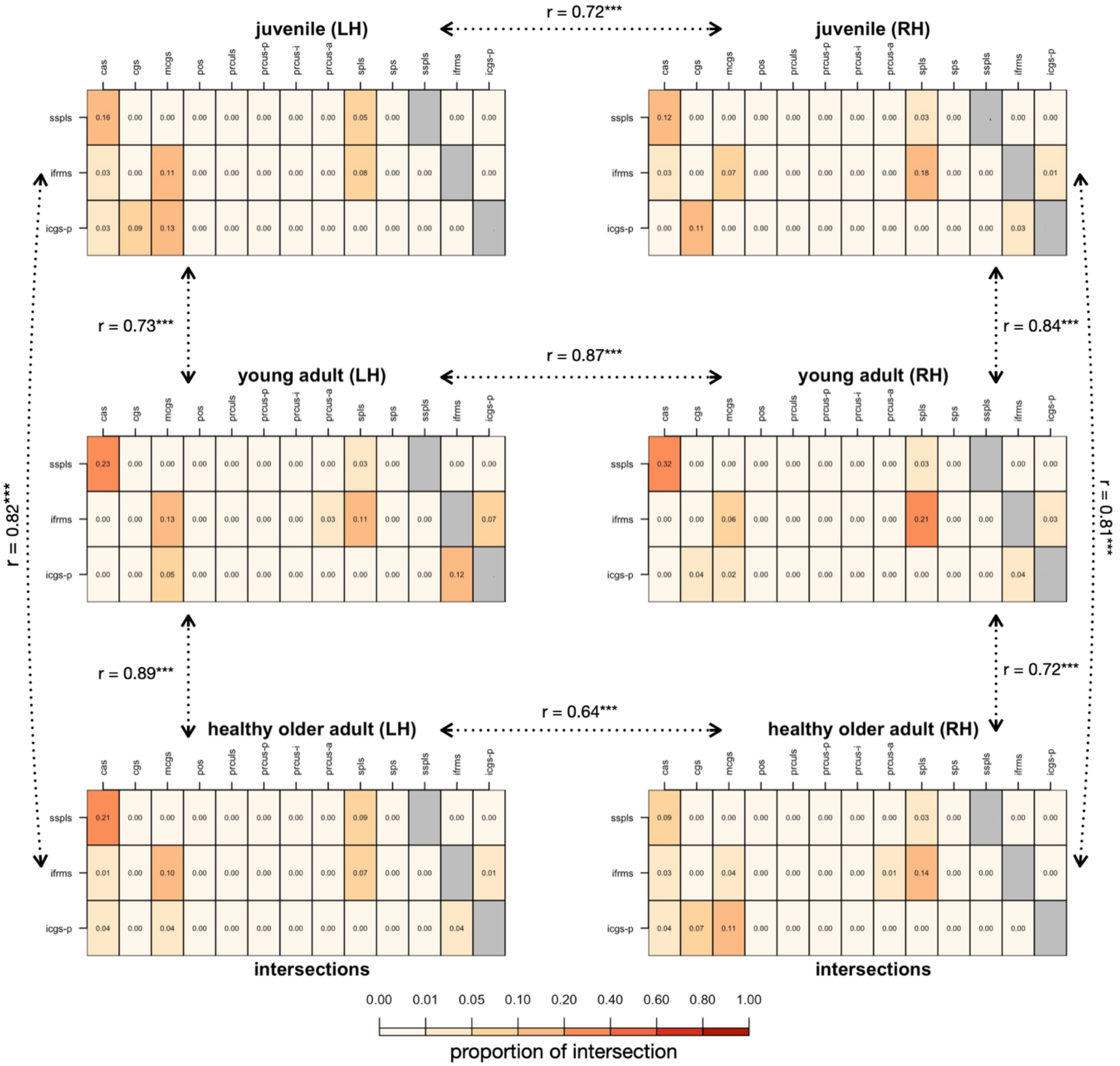


**Supplementary Figure 7.1. The sulcal patterns of shallow PCC sulci are similar between hemispheres and age groups.** Same format as Supplementary Fig. 2.2. For each shallow PCC sulcus (*sspls, ifrms, icgs-p*), we report the proportion of intersection (frequency of occurrence/total number of observations) with each PMC sulcus for each age group (juvenile, young adult, healthy older adult). Note that the callosal sulcus (cas) and cingulate sulcus (cgs) were also included as the sspls intersected with the cas and the icgs-p intersected with the cgs frequently. Calculating the correlation between matrices shows that the intersections of these sulci is comparable (all *r*s > .60; all ps < 0.001) between hemispheres and age groups. The three most prevalent types for each shallow sulcus in the juvenile and healthy older adult samples are included in Supplementary Tables 12 and 14, respectively.

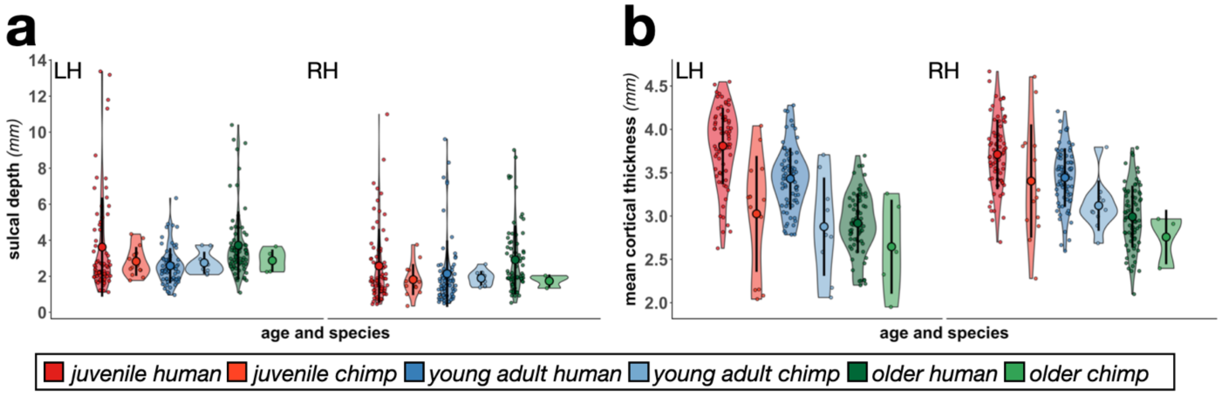


**Supplementary Figure 7.2. The morphological trends of the *ifrms* across age and species are the same regardless of normalization. a.** Same layout as Fig. 6c. Raw sulcal depth of the *ifrms,* calculated using a recent algorithm[1], across the lifespan and between species plotted for each individual participant in each hemisphere. The mean (large colored circles), standard deviation (black line), and kernel density estimate (colored "violin") are also plotted for each sulcus. There are also significant differences in raw depth of the *ifrms* between species and age groups as we found with normalized depth (Fig. 6c). **b.** Same layout as Fig. 6d, but for raw cortical thickness (mm). The *ifrms* shows an age- and species-related decrease in raw cortical thickness as for normalized cortical thickness (Fig. 6d).

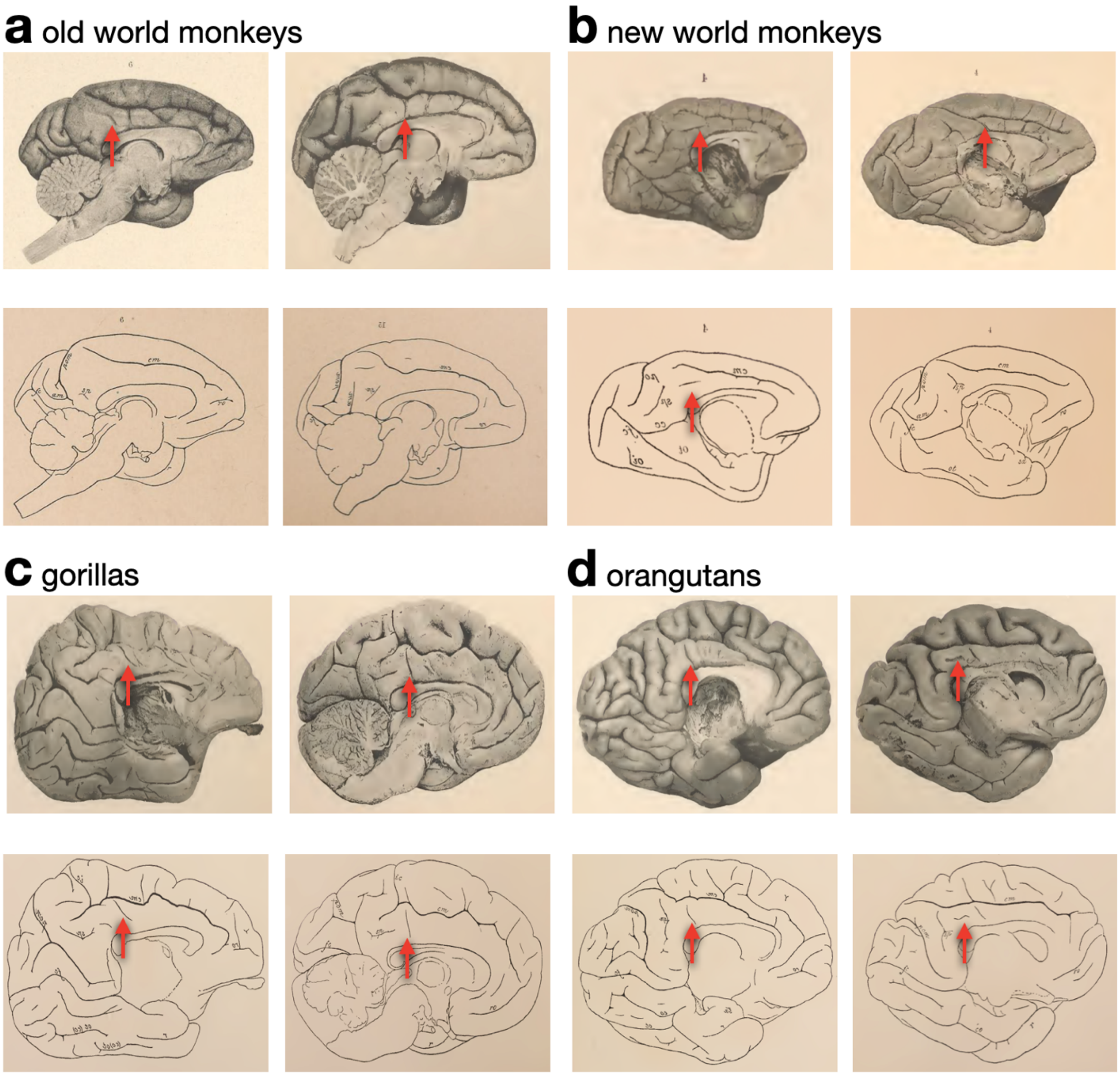


**Supplementary Figure 8. The *ifrms:* From dimple to sulcus across evolution? a.** Top: A shallow dimple (*ifrmd*; red arrow) is identifiable underneath the *mcgs* in Old World monkeys. Bottom: Schematic illustration of the sulcal patterning provided by Retzius[4]. Note that a shallow dimple is present underneath the *mcgs* in each photograph (red arrow), but not included in the schematic illustration. **b.** Same as **a.**, but for New World monkeys. Bottom left: Note that Retzius does include an unlabeled indentation in his schematic (red arrow). **c.** The *ifrms* labeled in two example gorilla hemispheres. **d.** The *ifrms* labeled in two example orangutan hemispheres. Overall, a shallow indentation (dimple) located underneath the *mcgs* was present in 63.83% (30/47) and 40% (4/10) of New World monkey and Old World monkey hemispheres, respectively. A sulcal indentation underneath the *mcgs* was identifiable in 75% (3/4) of gorilla and 75% (6/8) of orangutan hemispheres present in Retzius' atlas. Not pictured: the *ifrms* was identifiable in 83.33% (15/18) of chimpanzee hemispheres in the atlas. All post-mortem hemispheres inspected from Retzius[4] will be included on our lab website with the publication of this paper. Broadly speaking, the culmination of these data support the idea that cortical indentations beneath the *mcgs* are shallow dimples (the *ifrmd*) in

Old World and New World monkeys, which then deepen and become a tertiary sulcus (the *ifrms*) in human and non-human hominoids.

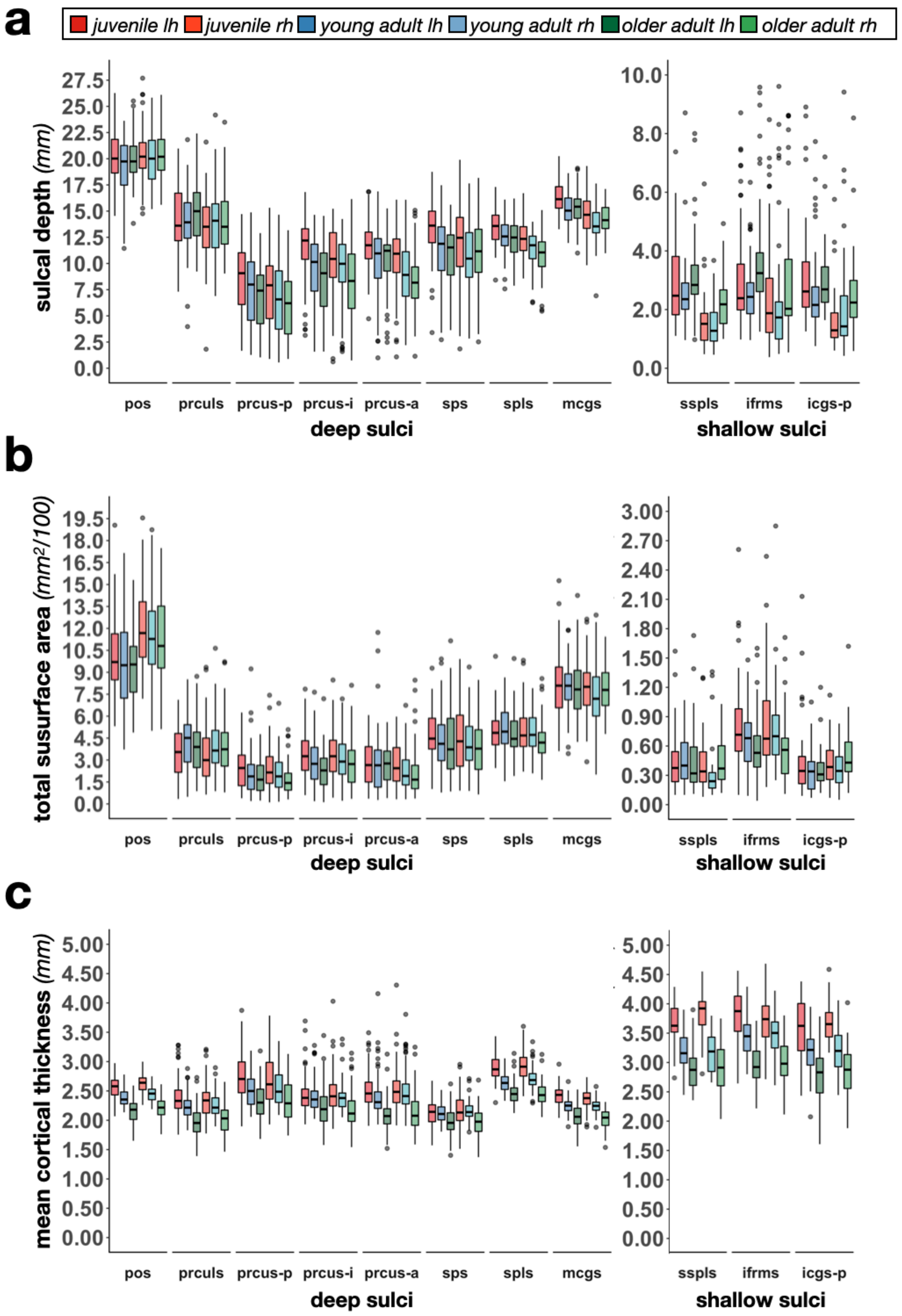

a
juvenile lh
juvenile rh
young adult lh
young adult rh
older adult lh
older adult rh
sulcal depth (mm)
pos
prculs
prcus-p
prcus-i
prcus-a
sps
spls
mcgs
deep sulci
sspls
ifrms
icgs-p
shallow sulci
b
total susurface area (mm2/100)
c
mean cortical thickness (mm)

**Supplementary Figure 9. Morphological features of PMC sulci across age groups in humans. a.** Sulcal depth (mm) of all 11 PMC sulci across the three age groups (juvenile, young adult, and healthy older adult) and hemispheres. The measures of central tendency for each sulcus are visualized with a box plot (outliers shown as dark gray circles). Sulci are ordered posterior to anterior along the x-axis and separated into deep and shallow sulci to appreciate the range of values for the shallower sulci. Each age group and hemisphere combination are colored according to the legend. **b.** Same as **a** but for total surface area ($mm^2$). Note that these values are scaled by a factor of 100. **c.** Same as **a** but for mean cortical thickness (mm). The mean and standard deviation values are also provided in Supplementary Tables 15-17.

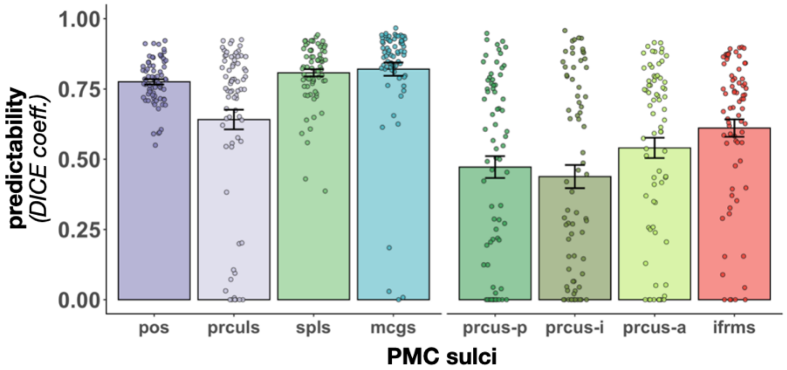


**Supplementary Figure 10. Automatically defining PMC sulci using deep learning algorithms.** Overlap (DICE coefficient) between predicted and manual location of PMC sulci that are identifiable in every hemisphere in young adults for spherical convolutional and context aware training. Bars represent average values, and the error bars indicate ±1 SEm. Circles represent each individual. Large, deep sulci are positioned to the left of the x-axis, while smaller, shallower sulci are positioned to the right. Predictability for the latter is lower than the former. However, predictability of the *ifrms* is higher than the three larger precuneal sulci (see Results).

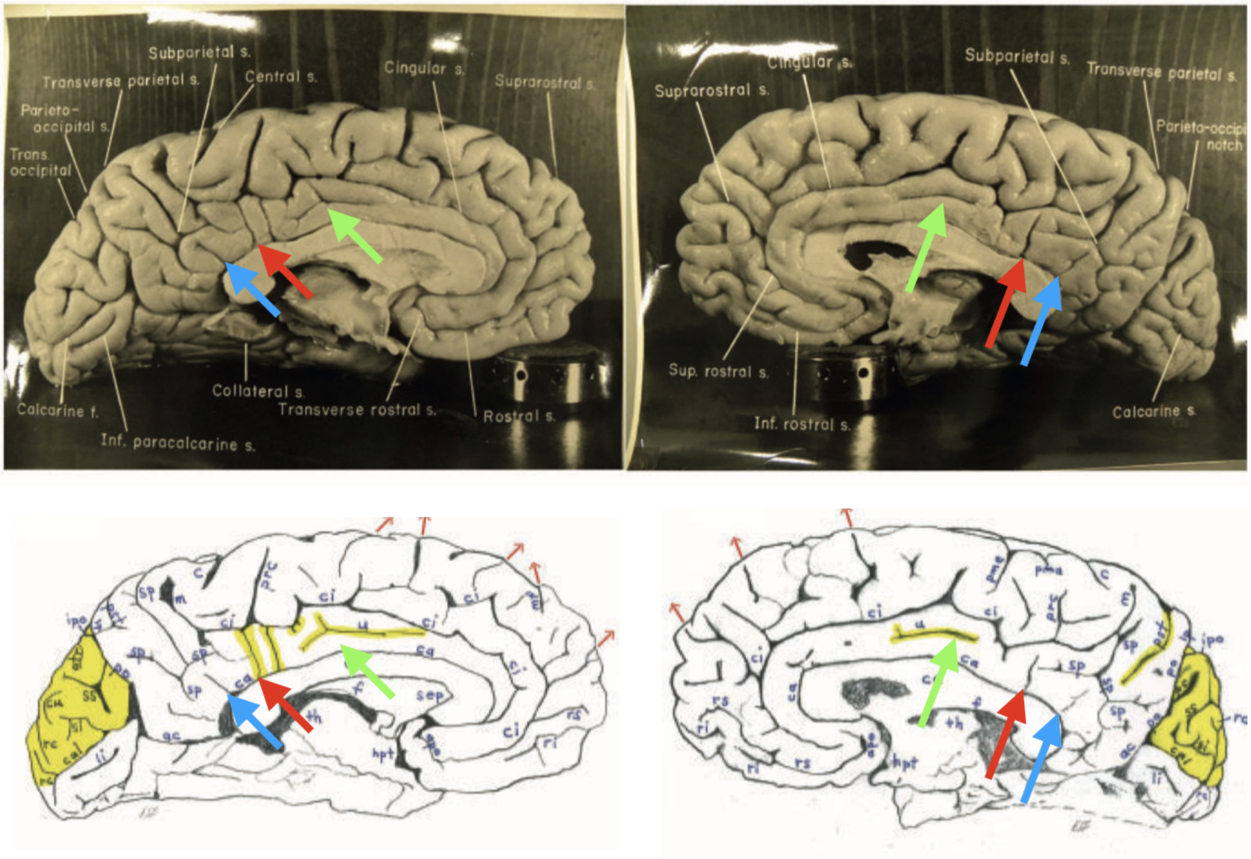


**Supplementary Figure 11. Even Einstein had an ifrms.** Top: Photographs of Einstein's brain. Blue arrow: sspls. Red arrow: ifrms. Green arrow: intracingulate sulcus as identified by Borne and colleagues[31]. Bottom: Schematic illustration highlighting (in yellow) differences in the sulcal patterning in Einstein's brain compared to typical sulcal patterning (from Falk *et al.*[15]). We highlight that the ifrms, and nearbhy shallow sulci, examined in the present study, are identified as abnormal in Einstein's brain. This highlights the necessity to identify all sulci and dimples of the cerebral cortex in order to accurately assess typicality and atypicality, as well as how individual differences in sulcal patterning relate to function, anatomy, and cognition in both typical and atypical brains. In this particular case, the omission of the ifrms and other "tiny" or shallow sulci in PCC resulted in the inaccurate conclusion that this was a special feature of Einstein's brain. Instead, this sulcal patterning in Einstein's PCC is actually common in humans and also present in many chimpanzee brains as we quantify in the present paper.

### Supplementary Tables

**Supplementary Table 1**

*Incidence rates of PMC sulci in the discovery sample*

| sulci | appearance in LH | LH % | appearance in RH | RH % |
|---|---|---|---|---|
| *pos* | 36/36 | 100.00 | 36/36 | 100.00 |
| *prculs* | 36/36 | 100.00 | 36/36 | 100.00 |
| *prcus-p* | 36/36 | 100.00 | 36/36 | 100.00 |
| *prcus-i* | 36/36 | 100.00 | 36/36 | 100.00 |
| *prcus-a* | 36/36 | 100.00 | 36/36 | 100.00 |
| *spls* | 36/36 | 100.00 | 36/36 | 100.00 |
| *sps* | 35/36 | 97.22 | 34/36 | 94.44 |
| *mcgs* | 36/36 | 100.00 | 36/36 | 100.00 |
| *sspls* | 23/36 | 63.89 | 18/36 | 50.00 |
| *ifrms* | 36/36 | 100.00 | 36/36 | 100.00 |
| *icgs-p* | 19/36 | 52.78 | 23/36 | 63.89 |

*Note.* This table illustrates the incidence rates of the 8-11 definable PMC sulci in the *discovery* young adult sample (N = 36 participants). The incidence rate of each sulcus (posterior to anterior) is provided in number and percent for both the left (LH) and right hemispheres (RH). The abbreviations used are as follows: anterior precuneal sulcus (*prcus-a*); intermediate precuneal sulcus (*prcus-i*); inframarginal sulcus (*ifrms*); marginal ramus of the cingulate sulcus (*mcgs*); parieto-occipital sulcus (*pos*); posterior intracingulate sulcus (*icgs-p*); posterior precuneal sulcus (*prcus-p*); precuneal limiting sulcus (*prculs*); splenial sulcus (*spls*); subsplenial sulcus (*sspls*); superior parietal sulcus (*sps*).

**Supplementary Table 2**

*Incidence rates of PMC sulci in the replication sample*

| sulci | appearance in LH | LH % | appearance in RH | RH % |
|---|---|---|---|---|
| *pos* | 36/36 | 100.00 | 36/36 | 100.00 |
| *prculs* | 36/36 | 100.00 | 36/36 | 100.00 |
| *prcus-p* | 36/36 | 100.00 | 36/36 | 100.00 |
| *prcus-i* | 36/36 | 100.00 | 36/36 | 100.00 |
| *prcus-a* | 36/36 | 100.00 | 36/36 | 100.00 |
| *spls* | 36/36 | 100.00 | 36/36 | 100.00 |
| *sps* | 30/36 | 83.33 | 34/36 | 94.44 |
| *mcgs* | 36/36 | 100.00 | 36/36 | 100.00 |
| *sspls* | 17/36 | 47.22 | 20/36 | 55.56 |
| *ifrms* | 36/36 | 100.00 | 36/36 | 100.00 |
| *icgs-p* | 24/36 | 66.67 | 23/36 | 63.89 |

*Note.* This table illustrates the incidence rates of the 8-11 definable PMC sulci in the *replication* young adult sample (N = 36 participants). The incidence rate of each sulcus (posterior to anterior) is provided in number and percent for both the left (LH) and right hemispheres (RH). The abbreviations used are as follows: anterior precuneal sulcus (*prcus-a*); intermediate precuneal sulcus (*prcus-i*); inframarginal sulcus (*ifrms*); marginal ramus of the cingulate sulcus (*mcgs*); parieto-occipital sulcus (*pos*); posterior intracingulate sulcus (*icgs-p*); posterior precuneal sulcus (*prcus-p*); precuneal limiting sulcus (*prculs*); splenial sulcus (*spls*); subsplenial sulcus (*sspls*); superior parietal sulcus (*sps*).

**Supplementary Table 3**

*Most common intersections of the discovery sample's precuneal sulci*

| Sulci | First | % | Second | % | Third | % |
|---|---|---|---|---|---|---|
| prcus-a | | | | | | |
| LH (n=36) | *spls* | 77.8 | *prcus-i* | 13.9 | *free* | 11.1 |
| RH (n=36) | *spls* | 58.3 | *mcgs* | 27.8 | *prcus-i* | 22.2 |
| prcus-i | | | | | | |
| LH (n=36) | *spls* | 47.2 | *free* | 38.9 | *prcus-p* | 13.9 |
| RH (n=36) | *spls* | 61.1 | *prcus-a* | 22.2 | *prcus-p* | 13.9 |
| prcus-p | | | | | | |
| LH (n=36) | *spls* | 55.6 | *free* | 36.1 | *prcus-i* | 13.9 |
| RH (n=36) | *spls* | 41.7 | *free* | 38.9 | *prcus-i* | 13.9 |

*Note.* This table illustrates the different sulcal patterns, or types, of the three precuneal sulci (*prcus*) identified in the *discovery* young adult sample (N = 36 participants). For each sulcus, the top three most prevalent sulcal patterns and their percent of occurrence are provided for both the left (LH) and right hemispheres (RH). The incidence of each sulcus in this sample is also provided for each hemisphere for reference. The abbreviations used are as follows: anterior precuneal sulcus (*prcus-a*); intermediate precuneal sulcus (*prcus-i*); marginal ramus of the cingulate sulcus (*mcgs*); no intersections (*free*); posterior precuneal sulcus (*prcus-p*); splenial sulcus (*spls*).

**Supplementary Table 4**

*Most common intersections of the discovery sample's shallow PCC sulci*

| Sulci | First | % | Second | % | Third | % |
|---|---|---|---|---|---|---|
| icgs-p | | | | | | |
| LH (n=19) | *free* | 78.9 | *ifrms* | 15.8 | *mcgs* | 5.3 |
| RH (n=23) | *free* | 95.7 | *mcgs* | 4.3 | --- | 0.0 |
| ifrms | | | | | | |
| LH (n=36) | *free* | 83.3 | *icgs-p* | 8.3 | *mcgs* | 5.6 |
| RH (n=36) | *free* | 80.6 | *spls* | 16.7 | *mcgs* | 2.8 |
| sspls | | | | | | |
| LH (n=23) | *free* | 73.9 | *cas* | 26.1 | --- | 0.0 |
| RH (n=18) | *free* | 77.8 | *cas* | 22.2 | --- | 0.0 |

*Note.* This table illustrates the different sulcal patterns, or types, of the three shallow PCC sulci identified in the *discovery* young adult sample (N = 36 participants). For each sulcus, the top three most prevalent sulcal patterns and their percent of occurrence are provided for both the left (LH) and right hemispheres (RH). The incidence of each sulcus in this sample is also provided for each hemisphere for reference. The abbreviations used are as follows: callosal sulcus (*cas*); inframarginal sulcus (*ifrms*); marginal ramus of the cingulate sulcus (*mcgs*); no intersections (*free*); no other option (---); posterior intracingulate sulcus (*icgs-p*); splenial sulcus (*spls*); subsplenial sulcus (*sspls*).

**Supplementary Table 5**

*Most common intersections of the replication sample's precuneal sulci*

| Sulci | First | % | Second | % | Third | % |
|---|---|---|---|---|---|---|
| prcus-a | | | | | | |
| LH (n=36) | *spls* | 83.3 | *prcus-i* | 22.2 | *mcgs* | 13.9 |
| RH (n=36) | *spls* | 72.2 | *prcus-i* | 19.4 | *mcgs* | 16.7 |
| prcus-i | | | | | | |
| LH (n=36) | *spls* | 63.9 | *prcus-a* | 22.2 | *free* | 22.2 |
| RH (n=36) | *spls* | 63.9 | *free* | 25.0 | *prcus-a* | 19.4 |
| prcus-p | | | | | | |
| LH (n=36) | *spls* | 55.6 | *free* | 33.3 | *prcus-i* | 13.9 |
| RH (n=36) | *spls* | 55.6 | *free* | 36.1 | *prcus-i* | 11.1 |

*Note.* This table illustrates the different sulcal patterns, or types, of the three precuneal sulci (*prcus*) identified in the *replication* young adult sample (N = 36 participants). For each sulcus, the top three most prevalent sulcal patterns and their percent of occurrence are provided for both the left (LH) and right hemispheres (RH). The incidence of each sulcus in this sample is also provided for each hemisphere for reference. The abbreviations used are as follows: anterior precuneal sulcus (*prcus-a*); intermediate precuneal sulcus (*prcus-i*); marginal ramus of the cingulate sulcus (*mcgs*); no intersections (*free*); posterior precuneal sulcus (*prcus-p*); splenial sulcus (*spls*).

**Supplementary Table 6**

*Most common intersections of the replication sample's shallow PCC sulci*

| Sulci | First | % | Second | % | Third | % |
|---|---|---|---|---|---|---|
| icgs-p | | | | | | |
| LH (n=24) | *free* | 87.5 | *ifrms* | 8.3 | *mcgs* | 4.2 |
| RH (n=23) | *free* | 82.6 | *ifrms* | 8.7 | *cgs* | 8.7 |
| ifrms | | | | | | |
| LH (n=36) | *free* | 66.7 | *spls* | 16.7 | *mcgs* | 11.1 |
| RH (n=36) | *free* | 61.1 | *spls* | 25.0 | *mcgs* | 8.3 |
| sspls | | | | | | |
| LH (n=17) | *free* | 76.5 | *cas* | 17.6 | *spls* | 5.9 |
| RH (n=20) | *free* | 55.0 | *cas* | 40.0 | *spls* | 5.0 |

*Note.* This table illustrates the different sulcal patterns, or types, of the three shallow PCC sulci identified in the *replication* young adult sample (N = 36 participants). For each sulcus, the top three most prevalent sulcal patterns and their percent of occurrence are provided for both the left (LH) and right hemispheres (RH). The incidence of each sulcus in this sample is also provided for each hemisphere for reference. The abbreviations used are as follows: callosal sulcus (*cas*); cingulate sulcus (*cgs*); inframarginal sulcus (*ifrms*); marginal ramus of the cingulate sulcus (*mcgs*); no intersections (*free*); posterior intracingulate sulcus (*icgs-p*); splenial sulcus (*spls*); subsplenial sulcus (*sspls*).

**Supplementary Table 7**

*Regression analysis summary for ifrms location predicting CCN-b location*

| RAS coordinate | df | $\beta$ | SE | t value | *p*-value | adjusted $R^2$ | F-statistic | adjusted *p*-value |
|---|---|---|---|---|---|---|---|---|
| discovery | | | | | | | | |
| right (LH) | 1, 33 | 0.61 | 0.16 | 3.77 | <0.001 | 0.28 | 14.23 | <0.001 |
| right (RH) | 1, 34 | 0.50 | 0.17 | 2.87 | 0.007 | 0.17 | 8.23 | 0.007 |
| anterior (LH) | 1, 33 | 0.57 | 0.11 | 5.37 | <0.001 | 0.45 | 28.84 | <0.001 |
| anterior (RH) | 1, 34 | 0.44 | 0.10 | 4.46 | <0.001 | 0.35 | 19.91 | <0.001 |
| superior (LH) | 1, 33 | 0.98 | 0.10 | 9.68 | <0.001 | 0.73 | 93.71 | <0.001 |
| superior (RH) | 1, 34 | 0.74 | 0.08 | 9.65 | <0.001 | 0.72 | 93.04 | <0.001 |
| replication | | | | | | | | |
| right (LH) | 1, 31 | 0.77 | 0.16 | 4.96 | <0.001 | 0.42 | 24.61 | <0.001 |
| right (RH) | 1, 32 | 1.14 | 0.39 | 2.91 | 0.007 | 0.18 | 8.44 | 0.007 |
| anterior (LH) | 1, 31 | 0.40 | 0.11 | 3.71 | 0.001 | 0.29 | 13.77 | 0.001 |
| anterior (RH) | 1,32 | 0.47 | 0.16 | 2.96 | 0.006 | 0.19 | 8.74 | 0.006 |
| superior (LH) | 1, 31 | 0.82 | 0.10 | 8.29 | <0.001 | 0.68 | 68.80 | <0.001 |
| superior (RH) | 1, 32 | 0.92 | 0.09 | 9.69 | <0.001 | 0.74 | 93.95 | <0.001 |

*Note.* This table provides the output of each linear regression run between each of the RAS (right, anterior, superior) coordinates of the inframarginal sulcus (*ifrms*; predictor variable) and cognitive control network B (CCN-b; outcome variable). A separate linear regression was run for each coordinate in each hemisphere (left (LH) and right (RH)) and sample (*discovery* and *replication*). The *p*-values presented in this table are FDR corrected for multiple comparisons. Exclusions: The LH of one participant in each sample was not included due to not having a CCN-b node near the *ifrms* and two participants from the *replication* sample were not included due to not having resting-state parcellations available. The other abbreviations used are as follows: degrees of freedom (df); regression beta coefficient ($\beta$); standard error (SE).

**Supplementary Table 8**

*Regression analysis summary for ifrms location predicting CCN-c location*

| RAS coordinate | df | $\beta$ | SE | t value | *p*-value | adjusted $R^2$ | F-statistic | adjusted *p*-value |
|---|---|---|---|---|---|---|---|---|
| discovery | | | | | | | | |
| right (LH) | 1, 34 | 0.80 | 0.16 | 5.07 | <0.001 | 0.41 | 25.75 | <0.001 |
| right (RH) | 1, 34 | 0.32 | 0.09 | 3.43 | 0.002 | 0.23 | 11.74 | 0.002 |
| anterior (LH) | 1, 34 | 0.43 | 0.11 | 3.87 | <0.001 | 0.28 | 14.94 | <0.001 |
| anterior (RH) | 1, 34 | 0.30 | 0.09 | 3.38 | 0.002 | 0.23 | 11.43 | 0.002 |
| superior (LH) | 1, 34 | 0.90 | 0.10 | 8.69 | <0.001 | 0.68 | 75.47 | <0.001 |
| superior (RH) | 1, 34 | 0.71 | 0.08 | 9.31 | <0.001 | 0.71 | 86.62 | <0.001 |
| replication | | | | | | | | |
| right (LH) | 1, 32 | 0.56 | 0.09 | 6.02 | <0.001 | 0.52 | 36.21 | <0.001 |
| right (RH) | 1, 32 | 0.55 | 0.14 | 3.82 | <0.001 | 0.29 | 14.62 | <0.001 |
| anterior (LH) | 1, 32 | 0.32 | 0.10 | 3.11 | 0.004 | 0.21 | 9.67 | 0.004 |
| anterior (RH) | 1,32 | 0.41 | 0.10 | 4.02 | <0.001 | 0.32 | 16.19 | <0.001 |
| superior (LH) | 1, 32 | 0.81 | 0.09 | 8.96 | <0.001 | 0.71 | 80.29 | <0.001 |
| superior (RH) | 1, 32 | 0.86 | 0.08 | 10.24 | <0.001 | 0.76 | 104.90 | <0.001 |

*Note.* This table provides the output of each linear regression run between each of the RAS (right, anterior, superior) coordinates of the inframarginal sulcus (*ifrms*; predictor variable) and cognitive control network C (CCN-c; outcome variable). A separate linear regression was run for each coordinate in each hemisphere (left (LH) and right (RH)) and sample (*discovery* and *replication*). The *p*-values presented in this table are FDR corrected for multiple comparisons. Exclusions: Two participants from the *replication* sample were not included due to not having resting-state parcellations available. The other abbreviations used are as follows: degrees of freedom (df); regression beta coefficient ($\beta$); standard error (SE).

**Supplementary Table 9**

*Incidence rates of PMC sulci in the juvenile sample*

| sulci | appearance in LH | LH % | appearance in RH | RH % |
|---|---|---|---|---|
| *pos* | 72/72 | 100.00 | 72/72 | 100.00 |
| *prculs* | 72/72 | 100.00 | 72/72 | 100.00 |
| *prcus-p* | 72/72 | 100.00 | 72/72 | 100.00 |
| *prcus-i* | 72/72 | 100.00 | 72/72 | 100.00 |
| *prcus-a* | 72/72 | 100.00 | 72/72 | 100.00 |
| *spls* | 72/72 | 100.00 | 72/72 | 100.00 |
| *sps* | 64/72 | 88.89 | 67/72 | 93.06 |
| *mcgs* | 72/72 | 100.00 | 72/72 | 100.00 |
| *sspls* | 38/72 | 52.78 | 34/72 | 47.22 |
| *ifrms* | 72/72 | 100.00 | 72/72 | 100.00 |
| *icgs-p* | 32/72 | 44.44 | 36/72 | 50.00 |

*Note.* This table illustrates the incidence rates of the 8-11 definable PMC sulci in the juvenile sample (N = 72 participants). The incidence rate of each sulcus (posterior to anterior) is provided in number and percent for both the left (LH) and right hemispheres (RH). The abbreviations used are as follows: anterior precuneal sulcus (*prcus-a*); intermediate precuneal sulcus (*prcus-i*); inframarginal sulcus (*ifrms*); marginal ramus of the cingulate sulcus (*mcgs*); parieto-occipital sulcus (*pos*); posterior intracingulate sulcus (*icgs-p*); posterior precuneal sulcus (*prcus-p*); precuneal limiting sulcus (*prculs*); splenial sulcus (*spls*); subsplenial sulcus (*sspls*); superior parietal sulcus (*sps*).

**Supplementary Table 10**

*Incidence rates of PMC sulci in the healthy older adult sample*

| sulci | appearance in LH | LH % | appearance in RH | RH % |
|---|---|---|---|---|
| *pos* | 72/72 | 100.00 | 72/72 | 100.00 |
| *prculs* | 72/72 | 100.00 | 72/72 | 100.00 |
| *prcus-p* | 72/72 | 100.00 | 72/72 | 100.00 |
| *prcus-i* | 72/72 | 100.00 | 72/72 | 100.00 |
| *prcus-a* | 72/72 | 100.00 | 72/72 | 100.00 |
| *spls* | 72/72 | 100.00 | 72/72 | 100.00 |
| *sps* | 59/72 | 81.94 | 55/72 | 76.39 |
| *mcgs* | 72/72 | 100.00 | 72/72 | 100.00 |
| *sspls* | 33/72 | 45.83 | 32/72 | 44.44 |
| *ifrms* | 72/72 | 100.00 | 72/72 | 100.00 |
| *icgs-p* | 27/72 | 37.50 | 28/72 | 38.89 |

*Note.* This table illustrates the incidence rates of the 8-11 definable PMC sulci in the healhty older adult sample (N = 72 participants). The incidence rate of each sulcus (posterior to anterior) is provided in number and percent for both the left (LH) and right hemispheres (RH). The abbreviations used are as follows: anterior precuneal sulcus (*prcus-a*); intermediate precuneal sulcus (*prcus-i*); inframarginal sulcus (*ifrms*); marginal ramus of the cingulate sulcus (*mcgs*); parieto-occipital sulcus (*pos*); posterior intracingulate sulcus (*icgs-p*); posterior precuneal sulcus (*prcus-p*); precuneal limiting sulcus (*prculs*); splenial sulcus (*spls*); subsplenial sulcus (*sspls*); superior parietal sulcus (*sps*).

**Supplementary Table 11**

*Most common intersections of the juvenile sample's precuneal sulci*

| Sulci | First | % | Second | % | Third | % |
|---|---|---|---|---|---|---|
| prcus-a | | | | | | |
| LH (n=72) | *spls* | 79.2 | *prcus-i* | 19.4 | *mcgs* | 9.7 |
| RH (n=72) | *spls* | 68.1 | *prcus-i* | 22.2 | *free* | 20.8 |
| prcus-i | | | | | | |
| LH (n=72) | *spls* | 68.1 | *prcus-a* | 22.2 | *free* | 20.8 |
| RH (n=72) | *spls* | 61.1 | *free* | 31.9 | *prcus-a* | 22.2 |
| prcus-p | | | | | | |
| LH (n=72) | *spls* | 55.6 | *free* | 33.3 | *prcus-i* | 15.3 |
| RH (n=72) | *spls* | 52.8 | *free* | 37.5 | *prcus-i* | 19.4 |

*Note.* This table illustrates the different sulcal patterns, or types, of the three precuneal sulci (*prcus*) identified in the juvenile sample (N = 72 participants). For each sulcus, the top three most prevalent sulcal patterns and their percent of occurrence are provided for both the left (LH) and right hemispheres (RH). The incidence of each sulcus in this sample is also provided for each hemisphere for reference. The abbreviations used are as follows: anterior precuneal sulcus (*prcus-a*); intermediate precuneal sulcus (*prcus-i*); marginal ramus of the cingulate sulcus (*mcgs*); no intersections (*free*); posterior precuneal sulcus (*prcus-p*); splenial sulcus (*spls*).

**Supplementary Table 12**

*Most common intersections of the juvenile sample's shallow PCC sulci*

| Sulci | First | % | Second | % | Third | % |
|---|---|---|---|---|---|---|
| icgs-p | | | | | | |
| LH (n=32) | *free* | 75.0 | *mcgs* | 12.5 | *cgs* | 9.4 |
| RH (n=36) | *free* | 83.3 | *cgs* | 11.1 | *ifrms* | 2.8 |
| ifrms | | | | | | |
| LH (n=72) | *free* | 80.6 | *mcgs* | 11.1 | *spls* | 8.3 |
| RH (n=72) | *free* | 73.6 | *spls* | 18.1 | *mcgs* | 5.6 |
| sspls | | | | | | |
| LH (n=38) | *free* | 78.9 | *cas* | 15.8 | *spls* | 5.3 |
| RH (n=34) | *free* | 85.3 | *cas* | 11.8 | *spls* | 2.9 |

*Note.* This table illustrates the different sulcal patterns, or types, of the three shallow PCC sulci identified in the juvenile sample (N = 72 participants). For each sulcus, the top three most prevalent sulcal patterns and their percent of occurrence are provided for both the left (LH) and right hemispheres (RH). The incidence of each sulcus in this sample is also provided for each hemisphere for reference. The abbreviations used are as follows: callosal sulcus (*cas*); cingulate sulcus (*cgs*); inframarginal sulcus (*ifrms*); marginal ramus of the cingulate sulcus (*mcgs*); no intersections (*free*); posterior intracingulate sulcus (*icgs-p*); splenial sulcus (*spls*); subsplenial sulcus (*sspls*).

**Supplementary Table 13**

*Most common intersections of the healthy older adult sample's precuneal sulci*

| Sulci | First | % | Second | % | Third | % |
|---|---|---|---|---|---|---|
| prcus-a | | | | | | |
| LH (n=72) | *spls* | 70.8 | *free* | 20.8 | *prcus-i* | 9.7 |
| RH (n=72) | *spls* | 75.0 | *free* | 15.3 | *mcgs* | 9.7 |
| prcus-i | | | | | | |
| LH (n=72) | *spls* | 54.2 | *free* | 30.6 | *prcus-a* | 9.7 |
| RH (n=72) | *spls* | 52.8 | *free* | 38.9 | *prcus-p* | 13.9 |
| prcus-p | | | | | | |
| LH (n=72) | *free* | 52.4 | *spls* | 44.4 | *prcus-i* | 6.9 |
| RH (n=72) | *spls* | 56.9 | *free* | 34.7 | *prcus-i* | 13.9 |

*Note.* This table illustrates the different sulcal patterns, or types, of the three precuneal sulci (*prcus*) identified in the healthy older adult sample (N = 72 participants). For each sulcus, the top three most prevalent sulcal patterns and their percent of occurrence are provided for both the left (LH) and right hemispheres (RH). The incidence of each sulcus in this sample is also provided for each hemisphere for reference. The abbreviations used are as follows: anterior precuneal sulcus (*prcus-a*); intermediate precuneal sulcus (*prcus-i*); marginal ramus of the cingulate sulcus (*mcgs*); no intersections (*free*); posterior precuneal sulcus (*prcus-p*); splenial sulcus (*spls*).

**Supplementary Table 14**

*Most common intersections of the healthy older adult sample's shallow PCC sulci*

| Sulci | First | % | Second | % | Third | % |
|---|---|---|---|---|---|---|
| icgs-p | | | | | | |
| LH (n=27) | *free* | 88.9 | *mcgs* | 3.7 | *cas* | 3.7 |
| RH (n=28) | *free* | 82.1 | *mcgs* | 10.7 | *cgs* | 7.1 |
| ifrms | | | | | | |
| LH (n=72) | *free* | 80.6 | *mcgs* | 9.7 | *spls* | 6.9 |
| RH (n=72) | *free* | 77.8 | *spls* | 13.9 | *mcgs* | 4.2 |
| sspls | | | | | | |
| LH (n=33) | *free* | 69.7 | *cas* | 21.2 | *spls* | 9.1 |
| RH (n=32) | *free* | 87.5 | *cas* | 9.4 | *spls* | 3.1 |

*Note.* This table illustrates the different sulcal patterns, or types, of the three shallow PCC sulci identified in the healthy older adult sample (N = 72 participants). For each sulcus, the top three most prevalent sulcal patterns and their percent of occurrence are provided for both the left (LH) and right hemispheres (RH). The incidence of each sulcus in this sample is also provided for each hemisphere for reference. The abbreviations used are as follows: callosal sulcus (*cas*); cingulate sulcus (*cgs*); inframarginal sulcus (*ifrms*); marginal ramus of the cingulate sulcus (*mcgs*); no intersections (*free*); posterior intracingulate sulcus (*icgs-p*); splenial sulcus (*spls*); subsplenial sulcus (*sspls*).

**Supplementary Table 15**
*Mean ± sd depth of each PMC sulcus between human age groups*

| | mean | sd |
|---|---|---|
| pos | | |
| j lh | 20.00 | 2.51 |
| ya lh | 19.29 | 2.48 |
| oa lh | 19.87 | 2.30 |
| j rh | 20.36 | 2.51 |
| ya rh | 20.06 | 2.53 |
| oa rh | 20.34 | 2.09 |
| prculs | | |
| j lh | 14.27 | 2.98 |
| ya lh | 13.97 | 2.92 |
| oa lh | 14.88 | 2.61 |
| j rh | 13.34 | 3.03 |
| ya rh | 13.82 | 3.17 |
| oa rh | 14.03 | 2.96 |
| prcus-p | | |
| j lh | 8.67 | 3.51 |
| ya lh | 7.52 | 3.45 |
| oa lh | 6.92 | 3.06 |
| j rh | 7.50 | 3.49 |
| ya rh | 6.62 | 3.45 |
| oa rh | 6.00 | 3.11 |
| prcus-i | | |
| j lh | 11.49 | 3.06 |
| ya lh | 9.42 | 3.30 |
| oa lh | 8.59 | 3.30 |
| j rh | 10.27 | 3.15 |
| ya rh | 9.43 | 3.26 |
| oa rh | 8.13 | 3.52 |
| prcus-a | | |
| j lh | 11.76 | 2.47 |
| ya lh | 10.36 | 3.33 |
| oa lh | 10.26 | 2.71 |
| j rh | 10.19 | 2.97 |
| ya rh | 8.75 | 3.12 |
| oa rh | 8.30 | 2.79 |
| sps | | |
| j lh | 13.42 | 2.78 |
| ya lh | 11.21 | 3.28 |
| oa lh | 10.71 | 3.19 |
| j rh | 11.94 | 3.53 |
| ya rh | 10.66 | 3.41 |
| oa rh | 10.84 | 3.36 |
| spls | | |
| j lh | 13.48 | 1.74 |
| ya lh | 12.63 | 1.61 |
| oa lh | 12.35 | 1.91 |
| j rh | 12.37 | 1.77 |
| ya rh | 11.41 | 1.78 |
| oa rh | 10.73 | 1.88 |
| mcgs | | |
| j lh | 16.26 | 1.62 |
| ya lh | 15.09 | 1.51 |
| oa lh | 15.26 | 1.58 |
| j rh | 14.70 | 1.87 |
| ya rh | 13.90 | 1.79 |
| oa rh | 14.22 | 1.39 |
| sspls | | |
| j lh | 2.93 | 1.47 |
| ya lh | 2.60 | 1.32 |
| oa lh | 3.49 | 1.95 |
| j rh | 1.68 | 1.11 |
| ya rh | 1.55 | 0.91 |
| oa rh | 2.26 | 0.91 |
| ifrms | | |
| j lh | 3.64 | 2.77 |
| ya lh | 2.59 | 0.98 |
| oa lh | 3.77 | 1.90 |
| j rh | 2.61 | 2.06 |
| ya rh | 2.14 | 1.85 |
| oa rh | 2.94 | 1.95 |
| icgs-p | | |
| j lh | 3.59 | 2.33 |
| ya lh | 2.64 | 1.54 |
| oa lh | 3.07 | 1.31 |
| j rh | 1.67 | 1.10 |
| ya rh | 2.18 | 1.81 |
| oa rh | 2.61 | 1.61 |

*Note.* Depth values are in millimeters. Abbreviations are as follows: juvenile (j), young adult (ya), older adult (oa), left hemisphere (lh), right hemisphere (rh)

**Supplementary Table 16**

*Mean ± sd surface area of each PMC sulcus between human age groups*

| | mean | sd |
|---|---|---|
| pos | | |
| j lh | 1006.85 | 270.56 |
| ya lh | 958.54 | 280.86 |
| oa lh | 947.81 | 247.88 |
| j rh | 1206.11 | 289.68 |
| ya rh | 1138.71 | 311.39 |
| oa rh | 1117.60 | 293.55 |
| prculs | | |
| j lh | 351.92 | 169.98 |
| ya lh | 428.85 | 189.51 |
| oa lh | 394.94 | 177.06 |
| j rh | 343.65 | 192.04 |
| ya rh | 403.65 | 185.88 |
| oa rh | 386.03 | 189.75 |
| prcus-p | | |
| j lh | 249.93 | 138.99 |
| ya lh | 209.86 | 155.11 |
| oa lh | 181.25 | 104.19 |
| j rh | 242.93 | 143.95 |
| ya rh | 204.99 | 125.99 |
| oa rh | 166.31 | 106.63 |
| prcus-i | | |
| j lh | 328.17 | 156.50 |
| ya lh | 286.42 | 155.12 |
| oa lh | 236.79 | 131.24 |
| j rh | 332.92 | 163.39 |
| ya rh | 306.71 | 156.50 |
| oa rh | 278.53 | 158.74 |
| prcus-a | | |
| j lh | 282.38 | 155.26 |
| ya lh | 274.86 | 206.74 |
| oa lh | 275.26 | 138.84 |
| j rh | 268.38 | 143.63 |
| ya rh | 231.62 | 137.72 |
| oa rh | 221.72 | 168.71 |
| sps | | |
| j lh | 480.22 | 170.72 |
| ya lh | 442.32 | 212.44 |
| oa lh | 427.49 | 243.23 |
| j rh | 447.66 | 227.96 |
| ya rh | 410.97 | 183.18 |
| oa rh | 391.29 | 204.93 |
| spls | | |
| j lh | 501.00 | 132.04 |
| ya lh | 510.28 | 146.03 |
| oa lh | 478.24 | 135.24 |
| j rh | 507.47 | 164.05 |
| ya rh | 491.22 | 143.83 |
| oa rh | 436.35 | 134.23 |
| mcgs | | |
| j lh | 813.00 | 220.27 |
| ya lh | 804.42 | 175.57 |
| oa lh | 793.69 | 213.04 |
| j rh | 789.97 | 188.75 |
| ya rh | 734.32 | 201.28 |
| oa rh | 783.26 | 164.19 |
| sspls | | |
| j lh | 45.08 | 32.46 |
| ya lh | 46.25 | 26.19 |
| oa lh | 44.61 | 36.25 |
| j rh | 43.50 | 29.98 |
| ya rh | 32.37 | 29.67 |
| oa rh | 43.56 | 23.76 |
| ifrms | | |
| j lh | 79.97 | 42.92 |
| ya lh | 67.18 | 31.91 |
| oa lh | 61.60 | 46.50 |
| j rh | 82.36 | 49.34 |
| ya rh | 76.56 | 39.03 |
| oa rh | 56.96 | 30.13 |
| icgs-p | | |
| j lh | 47.78 | 44.92 |
| ya lh | 34.44 | 20.52 |
| oa lh | 36.41 | 23.02 |
| j rh | 43.81 | 25.52 |
| ya rh | 37.09 | 20.09 |
| oa rh | 51.46 | 30.70 |

*Note.* Surface area values are in squared millimeters. Abbreviations are as follows: juvenile (j), young adult (ya), older adult (oa), left hemisphere (lh), right hemisphere (rh).

**Supplementary Table 17**

*Mean ± sd cortical thickness of each PMC sulcus between human age groups*

| | mean | sd |
|---|---|---|
| pos | | |
| j lh | 2.56 | 0.20 |
| ya lh | 2.39 | 0.14 |
| oa lh | 2.15 | 0.20 |
| j rh | 2.63 | 0.16 |
| ya rh | 2.45 | 0.13 |
| oa rh | 2.20 | 0.18 |
| prculs | | |
| j lh | 2.40 | 0.31 |
| ya lh | 2.24 | 0.20 |
| oa lh | 1.99 | 0.25 |
| j rh | 2.34 | 0.32 |
| ya rh | 2.27 | 0.21 |
| oa rh | 2.01 | 0.25 |
| prcus-p | | |
| j lh | 2.78 | 0.44 |
| ya lh | 2.52 | 0.29 |
| oa lh | 2.32 | 0.29 |
| j rh | 2.71 | 0.44 |
| ya rh | 2.55 | 0.30 |
| oa rh | 2.34 | 0.34 |
| prcus-i | | |
| j lh | 2.44 | 0.32 |
| ya lh | 2.40 | 0.27 |
| oa lh | 2.23 | 0.34 |
| j rh | 2.46 | 0.36 |
| ya rh | 2.40 | 0.25 |
| oa rh | 2.19 | 0.33 |
| prcus-a | | |
| j lh | 2.53 | 0.32 |
| ya lh | 2.41 | 0.35 |
| oa lh | 2.09 | 0.28 |
| j rh | 2.54 | 0.38 |
| ya rh | 2.52 | 0.36 |
| oa rh | 2.16 | 0.33 |
| sps | | |
| j lh | 2.13 | 0.21 |
| ya lh | 2.12 | 0.17 |
| oa lh | 1.96 | 0.23 |
| j rh | 2.16 | 0.28 |
| ya rh | 2.16 | 0.18 |
| oa rh | 1.97 | 0.25 |
| spls | | |
| j lh | 2.91 | 0.21 |
| ya lh | 2.65 | 0.16 |
| oa lh | 2.46 | 0.20 |
| j rh | 2.92 | 0.24 |
| ya rh | 2.71 | 0.18 |
| oa rh | 2.45 | 0.18 |
| mcgs | | |
| j lh | 2.42 | 0.17 |
| ya lh | 2.25 | 0.13 |
| oa lh | 2.05 | 0.21 |
| j rh | 2.36 | 0.17 |
| ya rh | 2.24 | 0.13 |
| oa rh | 2.02 | 0.17 |
| sspls | | |
| j lh | 3.67 | 0.35 |
| ya lh | 3.17 | 0.33 |
| oa lh | 2.87 | 0.39 |
| j rh | 3.81 | 0.39 |
| ya rh | 3.18 | 0.36 |
| oa rh | 2.93 | 0.40 |
| ifrms | | |
| j lh | 3.81 | 0.44 |
| ya lh | 3.43 | 0.36 |
| oa lh | 2.92 | 0.35 |
| j rh | 3.71 | 0.40 |
| ya rh | 3.44 | 0.34 |
| oa rh | 2.99 | 0.36 |
| icgs-p | | |
| j lh | 3.57 | 0.52 |
| ya lh | 3.17 | 0.41 |
| oa lh | 2.78 | 0.46 |
| j rh | 3.66 | 0.33 |
| ya rh | 3.20 | 0.38 |
| oa rh | 2.84 | 0.44 |

*Note.* Cortical thickness values are in millimeters. Abbreviations are as follows: juvenile (j), young adult (ya), older adult (oa), left hemisphere (lh), right hemisphere (rh)

**Supplementary Table 18**

*Scanning parameters of the healthy older adult participants*

| participants | scanner manufacturer | magnetic field strength (T) | TR (ms) | TE (ms) | voxel size ($mm^3$) |
|---|---|---|---|---|---|
| 26 | Siemens | 3 | 2300.0 | 3.0 | 1 x 1 x 1 |
| 8 | Siemens | 3 | 2300.0 | 3.0 | 1.1 x 1.1 x 1.2 |
| 6 | Philips | 3 | 6.5 | 2.9 | 1 x 1 x 1 |
| 6 | GE | 1.5 | 8.6 | 3.8 | 0.9 x 0.9 x 1.2 |
| 5 | Philips | 3 | 6.8 | 3.2 | 1 x 1 x 1.2 |
| 5 | Siemens | 3 | 2300.0 | 3.0 | 1 x 1 x 1.2 |
| 4 | GE | 1.5 | 8.9 | 3.9 | 0.9 x 0.9 x 1.2 |
| 2 | Siemens | 1.5 | 2400.0 | 3.5 | 1.3 x 1.3 x 1.2 |
| 2 | GE | 1.5 | 9.2 | 4.1 | 0.9 x 0.9 x 1.2 |
| 1 | Philips | 3 | 6.8 | 3.1 | 1 x 1 x 1.2 |
| 1 | Philips | 1.5 | 8.6 | 4.0 | 0.9 x 0.9 x 1.2 |
| 1 | Philips | 1.5 | 8.5 | 4.0 | 0.9 x 0.9 x 1.2 |
| 1 | GE | 1.5 | 9.2 | 4.0 | 0.9 x 0.9 x 1.2 |
| 1 | GE | 1.5 | 9.1 | 4.0 | 0.9 x 0.9 x 1.2 |
| 1 | GE | 1.5 | 7.0 | 3.9 | 0.9 x 0.9 x 1.2 |
| 1 | GE | 3 | 7.0 | 3.0 | 1 x 1 x 1.2 |
| 1 | Siemens | 1.5 | 3000.0 | 3.6 | 1.3 x 1.3 x 1.2 |

*Note.* This table illustrates the different scanning parameters used for each of our randomly-selected, healthy older adult participants from the Alzheimer's Disease Neuroimaging Initiative (ADNI) online database (http://adni.loni.usc.edu). For each different set of parameters, the number of participants, scanner manufacturer, magnetic field strength in Teslas (T), repetition time (TR) in ms, time to echo (TE) in ms, and voxel size in $mm^3$ are provided.
